# Parallelization of single-molecule binding kinetic measurements via protein barcode sequencing

**DOI:** 10.1101/2025.06.05.658126

**Authors:** Sebastian Hutchinson, Ghada H. Mansour, Ahmed Rehan, Adeline Pichard-Kostuch, Ellyn Redheuil, Brian D. Reed, Andrew D. Griffiths, Marco Ribezzi-Crivellari

## Abstract

Screening protein variants for desired functions has long relied on coupling of genotype (gene sequence) to phenotype (protein function), limiting the use of powerful single-molecule (SM) techniques. Here, we introduce a scalable SM screening method that bypasses this constraint by linking SM functional analysis to protein identity through SM protein sequencing. Protein variants are tagged with unique C-terminal peptide barcodes and loaded onto a semiconductor chip containing millions of nanowells. Protein-ligand interactions are monitored in real time at the SM level, and a dye-cycling strategy extends the measurable dynamic range, enabling quantification of slow dissociation rates typical of high-affinity interactions. After functional analysis, each protein molecule is identified by sequencing its barcode. We apply this method to 20 barcoded nanobodies spanning over 1,000-fold in affinity, yielding results consistent with published values and individual SM measurements. Our approach should accelerate protein engineering by enabling rapid, multiplexed SM screening of protein libraries.

## Introduction

Selecting and screening proteins with new or improved activities, such as ligand binding, is critical for the development of novel therapeutic, diagnostic, industrial, and research applications. Coupling genotype (gene sequence) to phenotype (protein function) is essential for functional assessment, and can be achieved by expressing and analyzing variants one by one in microtiter plates, or by physically coupling each gene to the protein it encodes, for example using phage or yeast display^1,2^.

Although binding affinity is a common metric for interaction strength, efficacy is often driven by association and dissociation kinetics^3–8^. High-throughput methods to measure affinities are widespread^9,10^, but kinetic measurements remain largely low-throughput. SM techniques can analyze interaction kinetics by tracking the binding and dissociation of fluorescently-labelled ligands to surface-immobilized proteins, producing characteristic telegraph-like signals^11^. SM techniques use small quantities of reagents, do not require assessment of the active fraction, and can access subtle phenotypes that are inaccessible to bulk methods^12^. SM studies of longlived interactions, such as high-affinity antibody-antigen interactions are impeded by photobleaching, despite improvements in dye chemistry^13^ and photo-protection^14,15^. However, dye-cycling, wherein dyes are cyclically replenished via short-lived interactions^16–19^, extends the dynamic range.

SM methods can analyze large numbers of molecules, but typically only one variant at a time^12,20,21^. Parallelization efforts have used DNA probes^22,23^ or DNA sequencing platforms to study protein-DNA or DNA-DNA interactions^24–26^. However, the inability to couple genotype to phenotype at the SM level makes it difficult to adapt such approaches for parallel SM analysis of multiple protein variants in complex mixtures.

Here, we demonstrate a parallelized SM analysis system that eliminates the need for a physical linkage between genotype and phenotype. Using the commercially available Platinum^®^ platform^27^, we measure the ligand-binding kinetics of individual protein molecules and then identify the proteins by Next-Gen Protein Sequencing™ (NGPS™)^27^. Platinum detects SM fluorescence using a semiconductor chip containing 2 million nanoscale reaction chambers, with integrated optical waveguide for evanescent illumination with a 532 nm pulsed laser and integrated complementary metal-oxide-semiconductor (CMOS) sensors for fluorescence detection (Fig. 1a)^27^. Compatible fluorophores are distinguished by fluorescence intensity and lifetime^27^. In our method, we express protein variants (here nanobodies) that are fused to C-terminal peptide barcodes and couple them to a loading complex containing streptavidin (Fig. 1b)^28^. Following attachment of the barcoded proteins in reaction chambers, we measure SM lig-and-binding kinetics, then cleave the proteins by specific proteolysis, revealing the peptide barcode of each molecule. Finally, we sequence the peptide barcodes using NGPS^27^ to determine the genotype of the variant in each reaction chamber (Fig. 1c). NGPS relies on measuring SM interactions between N-terminal amino acids (NAAs) and one of six fluorescently labelled NAA recognizers. These interactions are identified by software analysis and annotated as recognition segments (RSs)—clusters of pulsing from the repeated on-off binding of recognizers to their target NAAs. Freely diffusing aminopeptidases sequentially remove NAAs, revealing subsequent NAAs for detection^27^.

**Figure 1.**
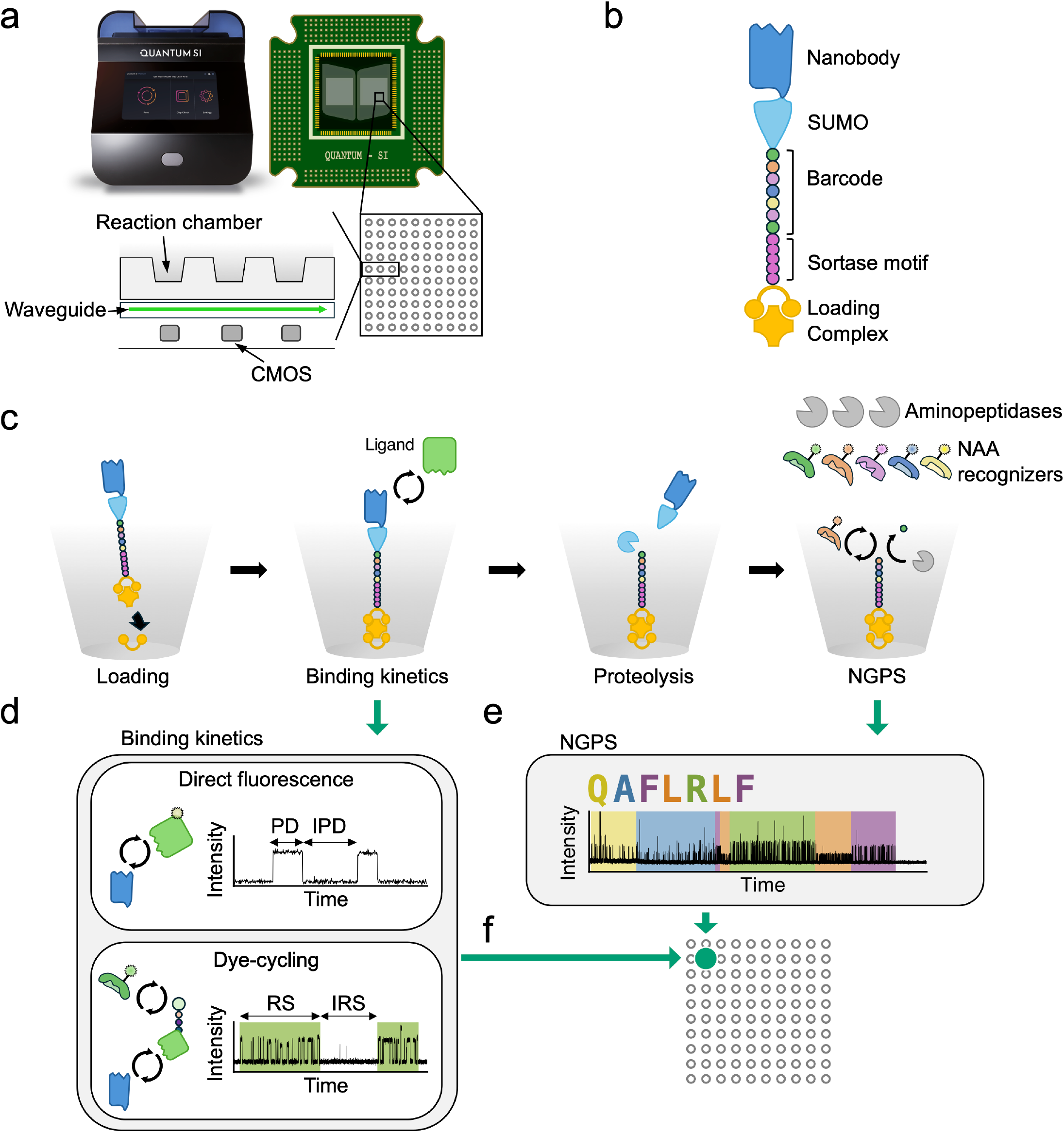
Parallel single molecule binding kinetics and protein barcode sequencing workflow. a) Platinum instrument (top left), and integrated semiconductor chip (top right). The chip contains 2 million nanowell reaction chambers organized in a 2D array (bottom right). The vertical cross-section (bottom left) shows a series of reaction chambers above, and integrated waveguide and CMOS detectors beneath. b) Recombinant nanobody construction. The nanobody is fused to the protein barcode via a SUMO tag. The C-terminal Sortase motif (LPETG) allows site-specific conjugation to the Platinum loading complex, a streptavidin-oligonucleotide conjugate carrying a DBCO moiety. c) Workflow. Loading: barcoded nanobodies are coupled via biotin-streptavidin interaction at the bottom of reaction chambers. Binding kinetics: labelled ligand is flowed into the reaction chamber, and bimolecular interactions are recorded. Proteolysis: the N-terminus of the barcode is exposed by proteolysis of SUMO by UlpI. NGPS: the peptide barcode is sequenced. The N-terminal residue is identified by the characteristic pulsing patterns of NAA recognizers. Aminopeptidases then expose subsequent residues for recognition. d) Detection regimes for SM binding kinetics. Direct fluorescence, the ligand is coupled to a fluorophore complex (yellow star). The trajectory (right) illustrates the telegraph signal, with PD and IPD shown. Dye-cycling: the ligand displays an NAA (green circle on ligand) that is bound by an NAA recognizer coupled to a fluorophore complex (green star). The trajectory (right) shows rapid pulsing due to repeated binding of NAA recognizers (black trace) to the ligand. RSs, regions of statistically similar pulsing behavior, are indicated as green boxes on the trajectory. Intervals between binding events (IRS) are indicated. e) NGPS results in dense pulsing patterns (black trace), which are annotated by primary analysis (colored boxes) based on binding kinetics and photonic properties, followed by peptide alignment, identifying the specific barcode sequence. f) Correlation of binding kinetic data and barcode sequence by mapping to the reaction chamber.

We developed two approaches to detect interactions: direct fluorescence and dye-cycling (Fig. 1d). For direct fluorescence, the ligand is coupled to a dye complex compatible with the Platinum system. Here, the inverse average pulse duration (PD) corresponds to the dissociation rate constant (k_*off*_) and the inverse average interpulse duration (IPD) corresponds to the association rate constant (k_*on*_) times the ligand concentration^11^ (Fig. 1d, top panel). Because this approach is limited by photobleaching, we developed a dye-cycling approach to measure low k_*off*_ values. We label ligands with peptide tags that are detected by fluorescently labeled NAA recognizers. The NAA recognizer functions as a fast-on, fast-off secondary reporter (Fig. 1d, bottom panel). We detect ligand-binding events as RSs, as for NGPS. Here, the duration of RSs correlates with the dissociation rate (k_*off*_), and inter-RS (IRS) duration with association rate (k_*on*_). This approach extends the dynamic range of the assay enabling measurement of binding events lasting thousands of seconds.

After aligning NGPS data to barcode sequences (Fig. 1e), we assign the protein variant in each reaction chamber to the SM binding kinetics data from the same reaction chamber (Fig. 1f) and perform statistical analysis on combined data for each variant. We applied this approach to a set of 20 nanobodies, with affinities varying more than 1,000-fold^29^. We show that parallel measurements correlate with both published values^29^ and our individual SM measurements.

## Results

### Measurement of SM binding kinetics by direct ligand labelling

To demonstrate SM binding kinetics measurement on Platinum, we separately analyzed nine anti-green fluorescent protein (GFP) nanobodies with affinities (K_*D*_) ranging from 6.7 × 10^−7^ - 6.9 × 10^−10^ M, (k_*off*_ 1.5 × 10^0^ - 1.1 × 10^−3^ s^−1^, k_*on*_ 3.2 × 10^5^ - 3.9 × 10^6^ M^−1^s^−1^) when measured using surface plasmon resonance (SPR)^29^, and a negative control nanobody, LaM1, that binds mCherry but not GFP^29^. We expressed and functionalized each nanobody and immobilized them at the bottoms of reaction chambers on a Platinum chip (Extended Data Fig. 1a-c, see Methods). We labelled GFP with a Platinum-compatible dye-complex^27^ (Fig. 2a, Extended Data Fig. 1d-f, see Methods) and added it to the chip to initiate the binding reaction. We first confirmed the expected concentration dependence of the association rate, and concentration independence of the dissociation rate, by measuring several concentrations of GFP binding with LaG42 (Extended Data Fig. 2a-e, Supplementary Information).

**Figure 2.**
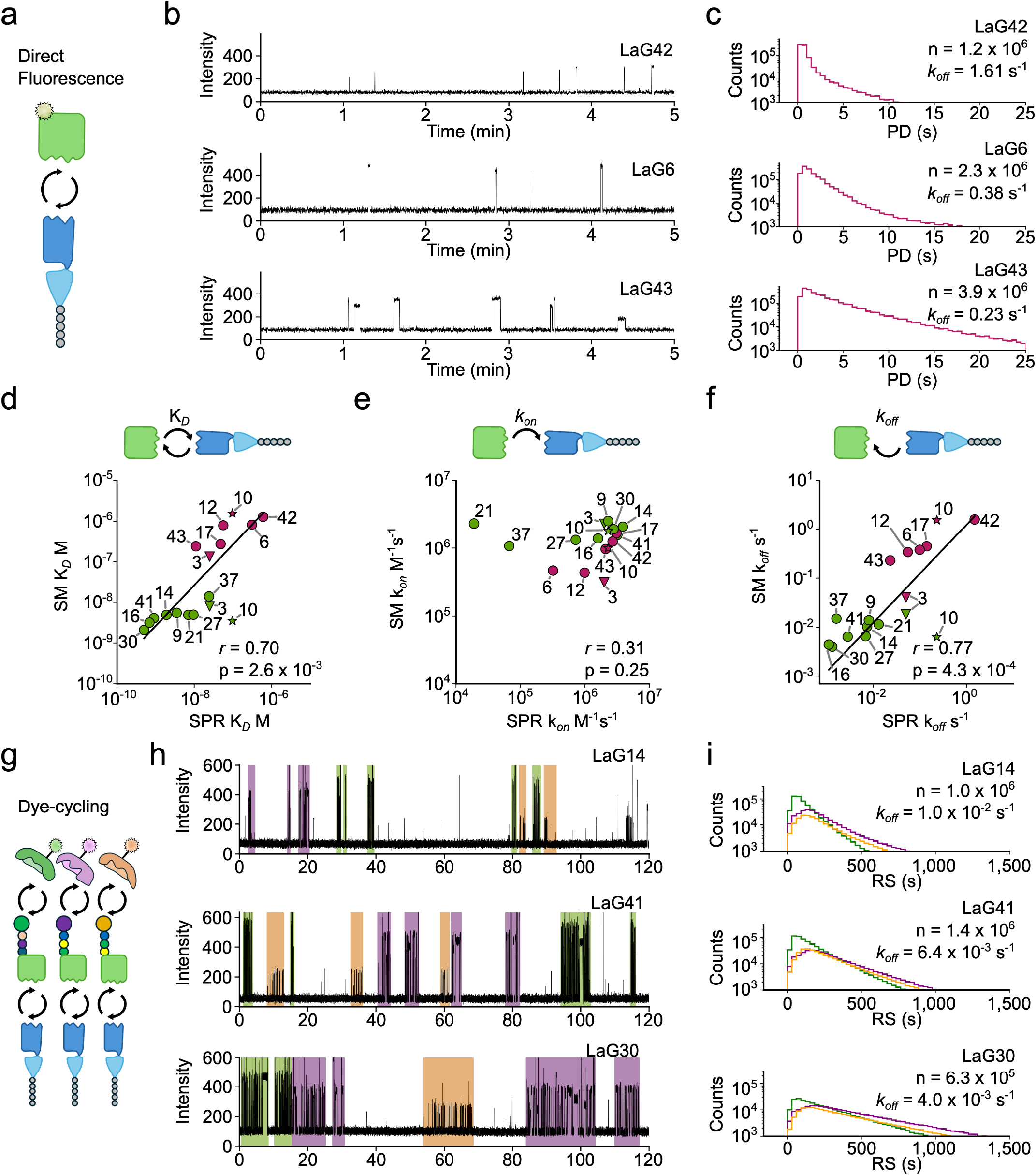
Single molecule binding kinetics on Platinum. a) Schematic of direct fluorescence experiments. GFP ligands are labelled with a fluorophore complex (yellow star) compatible with Platinum. b) Representative SM trajectories for the anti-GFP nanobodies LaG42, LaG6 and LaG43 with 7.5 nM GFP, measured by direct fluorescence. c) Histograms of pulse durations for LaG42, LaG6 and LaG43. The number of pulses (n) and the calculated k_*off*_ values are indicated. d) Comparison of K_*D*_ values from SM measurements to bulk SPR values^29^. Pearson correlation coefficient, *r*, and associated p-value are indicated. Solid line indicates fit by linear regression to y = ax, a = 2.62. Purple datapoints are from direct labelling and green datapoints are from dye cycling. Values for LaG10 and LaG3, which were measured by both direct labelling and dye cycling, are indicated as a star or a triangle, respectively (see main text). e,f) as for d but comparing k_*on*_ and k_*off*_, respectively. Solid line (f) indicates fit by linear regression to y = ax, a = 1.24. g) Schematic of dye-cycling experiments. NAA-GFP ligands are detected by their respective NAA recognizer. R/F/L-GFP are indicated with green, purple or orange, respectively. h) Representative SM trajectories for the anti-GFP nano-bodies LaG14, LaG41 and LaG30 with 7.5 nM GFP, measured by dye-cycling. Colored blocks indicate binding where the primary analysis algorithm called an RS. i) Histograms of RS durations for LaG14, LaG41 and LaG30. Line color corresponds to each of the three NAA-GFP types, as in h. The number of binding events (n) and the calculated k_*off*_ values are indicated.

We then recorded SM binding events for each nanobody for 2 h at 25 °C with 7.5 nM labelled GFP. We observed pulsing for anti-GFP nanobodies, but not LaM1 as expected (Extended Data Fig. 2f,g). Representative trajectories for three nanobodies, LaG42, LaG6 and LaG43 (Fig. 2b), and corresponding PD distributions (Fig. 2c) show clear differences, consistent with values measured by SPR^29^. Two nanobodies, LaG16 and LaG14, could not be measured due to photobleaching, which was quantified using the observed over expected number of pulses per nanowell (Extended Data Fig. 2h,i, see Methods). For the remaining nanobodies (LaG43, LaG6, LaG42, LaG10, LaG12, LaG17 and LaG3), we collected signal from a mean of 6.5 × 10^4^ nan-owells (5.9 × 10^3^ to 1.3 × 10^5^), with a mean of 1.4 × 10^6^ ligand binding events per variant (1.7 × 10^5^ to 3.9 × 10^6^) (Supplementary Table 1). We fit histograms of IPD and PD to a single-exponential decay model to obtain association k_*on*_ and dissociation k_*off*_, rate constants, respectively, and calculated the dissociation constant (K_*D*_ = k_*off*_/k_*on*_) (Fig. 2d-f, Extended Data Fig. 2j,k).

### Measurement of SM binding kinetics by dye-cycling

To measure high-affinity, slowly dissociating interactions relevant in most biomedical applications, we developed a dye-cycling approach, leveraging the rapid pulsing rate of NGPS NAA recognizers^27^. We engineered recombinant NAA-GFPs in which GFP was fused to either arginine (R-GFP), leucine (L-GFP), or phenylalanine (F-GFP) N-terminal peptides (Extended Data Fig. 3a-c). We confirmed recognition by NAA-recognizers by directly immobilizing NAA-GFPs in nanowells (Extended Data Fig. 3d-g, Supplementary Information).

We then implemented dye-cycling for SM measurement of binding kinetics. By using equimolar mixtures of three NAA-GFP ligands in solution (Fig. 2g), we ensured that consecutive binding events have different pulsing properties two-thirds of the time, facilitating and increasing the parameter space for RS annotation (see Methods)^27^. Dye-cycling data are not fit optimally by single-exponential distributions due to missed binding and unbinding events in trajectories and imprecision in the determination of bound times due to pulsing rates (Extended Data Fig. 4). We therefore designed a model to analyze dye-cycling data that accounts for these sources of error (see Methods). We applied this model to measurements of various concentrations of NAA-GFP binding to LaG16, confirming the expected concentration dependence of the association rate, and concentration independence of the dissociation rate (Extended Data Fig. 5, Supplementary Information).

We prepared thirteen anti-GFP nanobodies with affinities (K_*D*_), measured by SPR, ranging from 5.0 × 10^−10^ - 3.1 × 10^−7^ M (k_*off*_ 1.1 × 10^−3^ s^−1^ - 2.3 × 10^−1^ s^−1^, k_*on*_ 1.9 × 10^4^ - 3.9 × 10^6^ M^−1^s^−1^), and the anti-mCherry nanobody LaM1^29^, and measured SM ligand binding using dye-cycling (2 h, 25 °C and 7.5 nM NAA-GFP) (Extended Data Fig. 6). We recovered just 246 RSs for LaM1 and filtered out anti-GFP nanobodies that produced < 100-times this value of RSs from a single flow-cell (Supplementary Table 1). The three nanobodies filtered out—LaG6, LaG12 and LaG43—produced a mean of 6,100 RSs, and showed rapid dissociation in direct labelling experiments (Fig. 2d-f). For these three nanobodies the ratio of RS pulses to total pulses per nanowell, was lower than for the other nanobodies (≤ 0.323, Extended Data Fig. 6c). This dye-cycling efficiency metric decreases as dissociation rate becomes too fast to allow RS detection (Extended Data Fig. 6d). For the remaining ten anti-GFP nanobodies— LaG30, LaG16, LaG37, LaG41, LaG14, LaG9, LaG21, LaG27, LaG3, and LaG10—we collected signal from a mean of 1.5 × 10^5^ nanowells with a mean of 5.3 × 10^5^ RS per flow-cell (Supplementary Table 1).

Representative trajectories for LaG30, LaG41, and LaG14, are displayed in Fig. 2h. These trajectories show the three different types of RSs, for the three NAA-GFPs, each RS corresponding to a single GFP molecule binding. Distributions of RS durations correlate with SPR values, indicating that RSs report GFP binding events (Fig. 2i). We fit histograms of IRS and RS to the dye-cycling model to determine k_*on*_ and k_*off*_, respectively, and to calculate K_*D*_ (Fig. 2d-f).

### Combining direct ligand labelling and dye-cycling data

Combining the data from direct labelling and dye-cycling experiments allowed SM binding kinetics measurement over a wide dynamic range (Fig. 2d-f). K_*D*_ values varied over a 752-fold range, 1.6 × 10^−6^ M – 2.1 × 10^−9^ M, (Fig. 2d), and strongly correlated with SPR measurements^29^ (Pearson *r* = 0.70, *p* = 2.6 × 10^−3^). The k_*on*_ values spanned a 8.2-fold range, with Pearson *r* = 0.31, p = 0.25) compared to SPR measurements^29^ (Fig. 2e). This weak correlation is mostly due to two outliers, LaG37 and LaG21, with low k_*on*_ values in the SPR data^29^ that were not recapitulated in the SM measurements (after removing the outliers, *r* = 0.55, *p* = 0.040). Dissociation rate (k_*off*_) measurements spanned a 403-fold range, 1.61 × 10^0^ – 3.99 × 10^−3^ s^−1^, and showed strong correlation with SPR measurements (Pearson *r* = 0.77, *p* = 4.3 × 10^−4^) (Fig. 2f).

For two nanobodies, LaG3 and LaG10, k_*on*_ and k_*off*_ could be measured by both direct labelling and dye cycling. For LaG3, the k_*on*_ and k_*off*_ values were 4.1 × 10^−2^ s^−1^ (direct), and 1.8 × 10^−2^ s^−1^ (dyecycling) (Fig. 2e,f). For LaG10, k_*on*_ and k_*off*_ rates measured by direct labelling (1.0 × 10^6^ M^−1^s^−1^ and 1.6 s^−1^, respectively) were similar to reported values from SPR^29^. However, with dye-cycling we measured a similar k_*on*_ (1.8 × 10^6^ M^−1^s^−1^) but a k_*off*_ of 6.3 × 10^−3^ s^−1^. We tentatively propose that this result could be explained by conformational dynamics, where LaG10 is able to transition between different binding-competent states (see Discussion).

### Parallel Single-Molecule Binding Kinetics

We designed and characterized a set of 20 error-resistant, seven-amino-acid-long peptide barcodes encoding combinations of L, F, R, Q, or A for sequencing with Platinum v1 chemistry (Fig. 3a, Extended Data Fig. 7, Supplementary Information)^27^. We expressed and functionalized 19 anti-GFP nanobodies along with the anti-mCherry antibody LaM1^29^, each fused at the C-terminus with a distinct peptide barcode (Fig. 3a). We then pooled all 20 barcoded proteins at equimolar ratios and characterized ligand binding using both direct and dye-cycling measurements, each for 2 h at 25 °C, followed by NGPS barcode sequencing and peptide alignment. We detected a mean of 78.5% of residues within the aligned traces (Fig. 3b) and upon alignment of traces to barcodes, we could associate reaction chambers across the chip with specific nanobody variants (Fig. 3c). Representative trajectories in Figure 3d show linked binding kinetics data and barcode sequence data for three nanobodies measured by direct fluorescence and three measured by dye-cycling.

**Figure 3.**
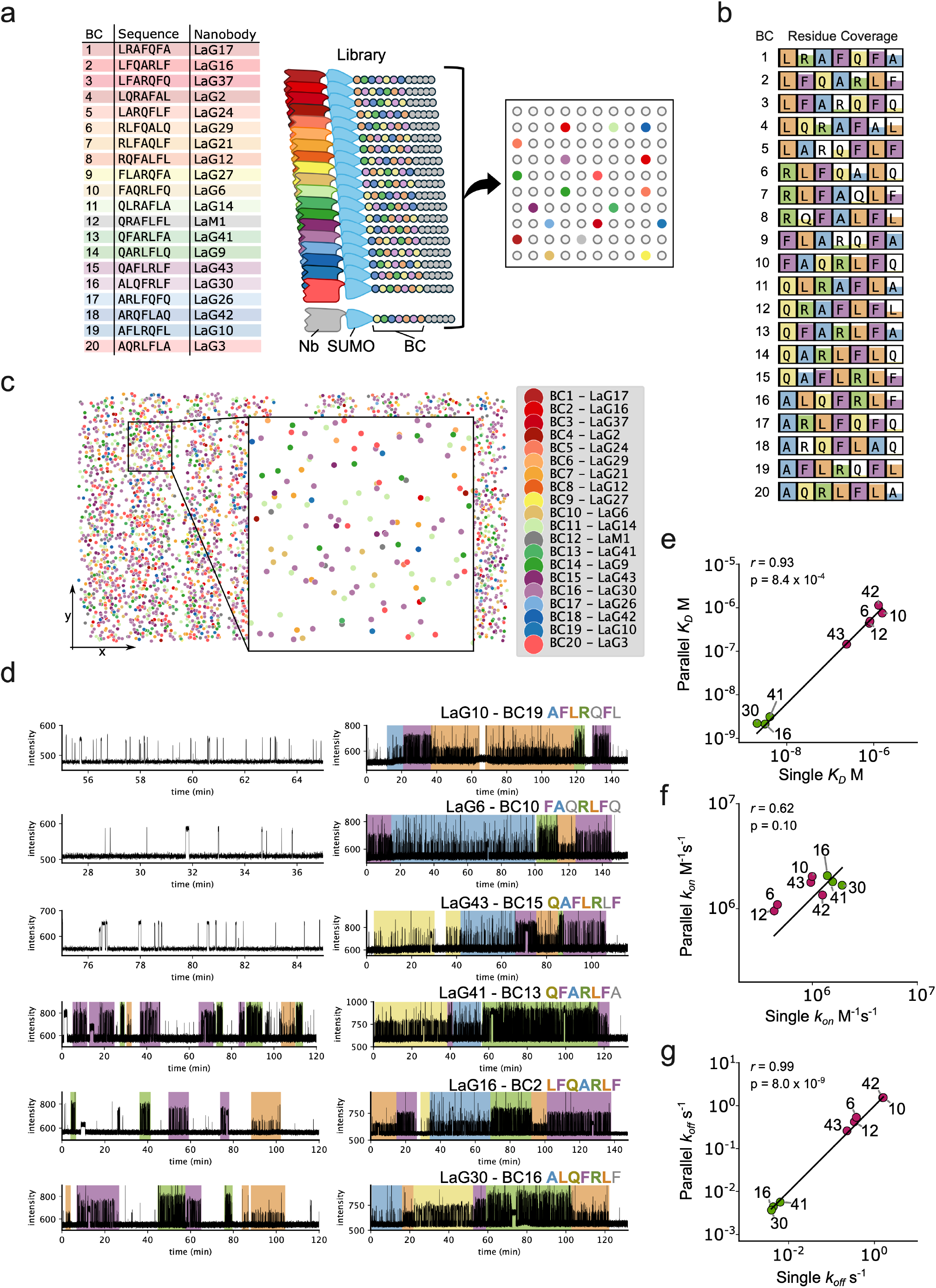
Parallel measurement of single molecule binding kinetics. a) Barcode number, sequences and linked nanobody (left) and experimental schematic (right). All barcoded nanobodies are mixed in equimolar ratios before being distributed into reaction chambers. b) Alignment coverage for each barcode, from a binding kinetics/NGPS experiment. Boxes show individual residues, and the proportion of the box that is filled indicates the fraction of alignments in which that residue was identified. c) Position of alignments for each barcode on the reaction chamber array from a single experiment. d) Linked SM trajectories for identical nanowells from binding kinetics and barcode readout by NGPS. Representative SM trajectories for ligand binding kinetics and associated barcode sequences of the anti-GFP nanobodies LaG10, LaG6 and LaG43 measured by direct labelling, and for LaG41, LaG16 and LaG30 measured by dye-cycling with 7.5 nM GFP. The barcode sequences are indicated. Detected residues are colored in bold and missed residues are in gray. e-g) Comparison of K_*D*_ (e), k_*on*_ (f) and k_*off*_ (g) values from parallel and individual measurements of nanobodies. Pearson correlation coefficient, *r*, and associated p-value is indicated. Solid lines indicate fit by linear regression to y = ax, where a = 0.63, 1.28 and 0.99, for K_*D*_, k_*on*_ and k_*off*_, respectively. Purple datapoints are from direct labelling and green datapoints are from dye cycling.

Filtering of parallelized data is complicated by cross-talk, derived from multiply loaded nanowells (Supplementary Information). We therefore developed one-sided Student’s t-tests comparing data for each anti-GFP nanobody to LaM1, filtering out variants that are not statistically significantly different from LaM1, which does not bind GFP^29^ (p > 0.01) (Extended Data Fig. 8a,b, Methods). After filtering, we obtained rate constants for 9 nanobodies: LaG42, LaG10, LaG12, LaG6, and LaG43 for direct labelling; and LaG30, LaG16, LaG41, and LaG24 for dye-cycling (Fig. 3e-g, Extended Data Fig. 8c-f, Supplementary Table 1). We observed a strong correlation between the K_*D*_, and k_*off*_ values, and weak correlation between k_*on*_ values from parallelized and individual variant experiments (Fig. 3e-g): Pearson *r* = 0.93 (p = 8.4 × 10^−4^) and *r* = 0.99 (p = 8.0 × 10^−9^), *r* = 0.62, (p = 0.10) respectively. Our analysis demonstrates that deconvolution of SM kinetic data using peptide barcode NGPS is efficient and reproducible.

## Discussion

Using the commercially available Platinum SM protein-sequencing instrument, we designed a simple workflow that measures SM binding kinetics for many protein variants in parallel. We load a variant library onto a flow cell, capture protein molecules in reaction chambers (nanowells), and monitor each molecule’s ligand-binding kinetics. We then identify variants by NGPS, removing the need for genotype-to-phenotype coupling. This yields an unprecedented amount of SM kinetic data: over 10^4^ coupled barcode NGPS-ligand binding traces per flow cell in a single experiment. The assay requires minimal hands-on time (~2 h), reagent preparation is implemented with basic molecular biology equipment and removes esoteric steps such as glass functionalization^30–33^. Data are processed automatically using Quantum-Si’s cloud computing architecture or locally^34^.

We developed a dye-cycling approach^16^ to measure interactions with slow dissociation rates in which ligands labeled with N-terminal peptides are detected by the same fluorescently labeled NAA recognizers used for NGPS^27^. Analysis of 19 anti-GFP nanobodies demonstrated that by combining direct ligand labelling and dye-cycling, we could measure k_*off*_ values across three orders of magnitude (1.6 × 10^0^ to 3.7 × 10^−3^ s^−1^), beyond the photobleaching limit. The k_*on*_ values varied only ~9-fold (3.1 × 10^5^ M^−1^s^−1^ to 2.9 × 10^6^ M^−1^s^−1^), consistent with lower variability in k_*on*_ generally observed for antibody-protein interactions^35^. Dissociation constants (K_*D*_) varied over three orders of magnitude (1.6 × 10^−6^ M to 2.1 × 10^−9^ M). The data recapitulate well bulk SPR measurements^29^.

With single nanobody measurements there is a small overlap in the dynamic range of direct labelling and dye-cycling: LaG3 could be characterized by both direct labelling and dye-cycling, with similar k_*off*_ values. However, in parallel experiments there is no overlap in dynamic range. Filtering excludes nanobodies with signals that do not exceed noise arising from nanowells containing multiple nanobodies, retaining nanobodies with the fastest dissociation rates for direct labelling and, conversely, the slowest for dye-cycling, hence removing variants with intermediate dissociation rates. In future we may be able to close this gap. For direct labelling, reducing photobleaching by modulating laser power or improving dye chemistry^13^ could allow measurement of longer interactions. For dye-cycling, increasing the pulsing rate, for instance by increasing the NAA recognizer concentration, or optimizing the NAA and penultimate residues for faster pulsing rates, would allow measurement of shorter interactions.

Binding kinetics of the anti-GFP nanobody LaG10 could also be measured by direct labelling and dye-cycling, both alone and in parallel. Measured k_*on*_ and k_*off*_ values from dye-cycling and k_*on*_ values from dye-cycling were similar to those measured by SPR^29^ (Fig. 2e,f, 3f,g); however, the value of k_*off*_ measured by dye-cycling was 244-248-times lower than by direct labelling (Fig. 2e,f, 3f,g). Based on the reported k_*off*_ from SPR^29^, the probability of an interaction lasting beyond the mean bound time in dyecycling of 160 s is 1.2 × 10^−16^. We propose that this discrepancy could be explained by conformational dynamics, where LaG10 molecules transition between different binding-competent states. While bulk approaches mask subtle sub-populations, SM techniques can reveal conformational dynamics^36,37^, but to our knowledge have not been applied to nanobody binding kinetics. Structural data suggests nanobody conformational plasticity is inversely correlated with binding affinity^38^. In some instances, however, introducing destabilizing mutations increased affinity^39^, and a direct link between conformation and binding kinetics has not been demonstrated. More generally, conformational isomerism, resulting in state-specific binding kinetics and multispecificity has been observed in antibodies and may be widespread^40,41^. Conformational dynamics have also been proposed to explain the behavior of single enzyme molecules^36,37,42^, including a DNA polymerase^43^.

Scaling up the number of barcode peptides from the 20 validated here will be key to expanding the utility of this technology. Our barcodes were designed for v1 Platinum chemistry, which recognizes 11 residues using 5 NAA recognizers. The newer Platinum v3 chemistry recognizes 13 residues, using 6 NAA recognizers. Projecting our current design scheme onto the new chemistry suggests 1,000-member barcode sets are possible (Supplementary Information). Existing chips, with 2 million nanowells, yield alignments from ~ 2 × 10^4^ nanowells per chip; at >100x coverage per variant, this would allow screening ~ 200 variants per experiment. Future technological developments will expand number of nanowells per chip by several orders of magnitude, allowing similar increases in the size of variant panels. Initial tests adapting this workflow to *in vitro* expression of variant libraries are also encouraging and suggest that the method will integrate well with synthetic biology workflows. It should also be possible to adapt our method for parallel SM measurement of enzymatic activity.

We have also co-developed a SM-parallelization method based on tagging each variant with an oligonucleotide barcode, decoded using the SM binding kinetics of complementary fluorescent single-stranded DNA probes^44^. However, when using genetically encoded peptide barcodes, coupling of oligonucleotide barcodes is not required. Instead, multiple genes encoding variant proteins with peptide barcodes can be co-expressed, greatly simplifying library preparation and streamlining the workflow.

The ability to generate large, high-quality SM datasets makes our method particularly well-suited for training or refining generative machine learning models focused on biophysical properties. Moreover, its simplicity lends itself to active learning frameworks involving iterative screening and refinement.

## Acknowledgements

We thank Margarida Gomes from Quantum-Si and members of the Laboratory of Biochemistry for helpful discussions. This work was financially supported by Quantum-Si.

## Author contributions

Conceptualization: SH, BR, ADG, MRC, Methodology: SH, GHM, APK, BR, ADG, MRC, Software: SH, MRC, Validation: SH, GHM, AR, ER, ADG, MRC, Formal analysis: SH, GHM, ADG, MRC, Investigation: SH, GHM, AR, APK, Resources: BR, ADG, MRC, Data Curation: SH, Writing - Original Draft: SH, GHM, MRC, Writing - Review & Editing: SH, GHM, BR, ADG, MRC, Visualization: SH, GHM, BR, ADG, MRC, Supervision: SH, BR, ADG, MRC, Project administration: SH, BR, ADG, MRC, Funding acquisition: BR, ADG, MRC.

## Materials and Methods

### Peptide-barcoded nanobody expression and functionalization

We constructed a modular plasmid, pPBC^Halo^, derived from the PURExpress DHFR Control Plasmid (New England Biolabs [NEB] N0424AVIAL) to facilitate recombinant protein expression in *Escherichia coli* and the protein functionalization and processing steps in our workflow. The plasmid contains an expression cassette, incorporating, from 5’ to 3’, a T7 RNA polymerase promoter, ribosome binding site (RBS), start codon, N-terminal Halo tag gene (the Halotag serves as a handle for affinity purification), tobacco etch virus (TEV) proteolysis sequence, two *Esp*3I restriction sites for scarless insertion of a protein of interest, a small ubiquitin like modifier (*SUMO*) gene, which is used as a proteolysis site to expose the N-terminus of the peptide barcode, two *Bbs*I restriction sites to scarlessly insert a peptide barcode sequence, followed by an LPETGG Sortase A motif^45,46^ for functionalization with the loading complex (K-linker, Quantum-Si), a hexahistidine tag, a stop codon and a T7 repressor sequence (Extended Data Fig. 1a). Barcoded nanobody expression plasmids were constructed in one step by Gibson Assembly (NEB E2611L). The pPBC^Halo^ backbone was amplified by PCR using Q5 Hot Start Polymerase (NEB M0493L) with primers Halo-SUMO_F and Halo-SUMO_R (Supplementary Table 2) to include the necessary flanking regions. The PCR conditions were as follows: for a 50 µL PCR reaction, the components included 10–50 ng of template DNA, 1 µL 10 mM dNTPs, 2.5 µL of 10 µM forward and reverse primers, 0.5 µL of Q5 Hot Start High-Fidelity DNA Polymerase in 10 µL 5X Q5 Hot Start Polymerase buffer supplemented with 5 µL 5X Q5 High GC Enhancer and nuclease free water to 50 µL. The PCR thermocycling conditions were as follows: initial denaturation at 98 °C for 30 seconds, followed by 25 cycles of denaturation at 98 °C for 30 seconds, annealing at 60 °C for 20 seconds, and extension at 72 °C for 2 minutes, with a final extension at 72 °C for 2 minutes. The PCR product was purified using a PCR cleanup kit (Macherey Nagel). Synthetic DNA constructs encoding the nanobody fused to SUMO and the peptide barcode (BC) at the C-terminus were synthesized (eBlocks, Integrated DNA Technologies) in a format compatible with insertion after the TEV proteolysis sequence (Supplementary Information). Vector backbone and insert were assembled in a 1:7 ratio with Gibson Assembly Master Mix (NEB) and incubated at 50°C for 1 hour. Plasmids were transformed into SHuffle T7 Express competent *E. coli* (NEB C3029J) according to the manufacturers protocol and assembly confirmed by Sanger sequencing using sequencing primer Sanger_R (Eurofins Genomics), resulting in pPBC^Halo^-BC#^−^LaG#, where # refers to either the barcode or LaG number (Supplementary Information).

Recombinant protein was produced by inoculating LB broth, supplemented with 100 µg/mL ampicillin, with a glycerol stock of the transformed colony containing the correctly assembled plasmid and incubating overnight at 30 °C with shaking at 180 r.p.m. The culture was then diluted to OD_600nm_ 0.1-0.2 in 10 mL LB broth and grown at 30 °C until OD_600nm_ 0.4 - 0.6. Protein expression was induced by addition of isopropyl-beta-D-thiogalactoside (IPTG) (Sigma Aldrich) at 0.5 mM concentration and bacteria incubated at 30 °C overnight with shaking at 180 r.p.m. Cells were pelleted at 4,000 x g for 10 minutes and lysed using NEBexpress cell lysis reagent (NEB P8116L) for 30 min at 25 °C, before clarifying the lysate by centrifugation at 10,000g for 10 minutes. Recombinant nanobody proteins were purified from the crude lysate using His-Pur™ Ni-NTA Resin (Thermo Fisher 88222). Before purification, 300 µL HisPur™ Ni-NTA Resin per sample was equilibrated by washing 3 x in 500 µL of binding buffer (50 mM HEPES pH 7.3, 150 mM NaCl and 10 mM imidazole). Clarified bacterial lysate was incubated with the equilibrated Ni-NTA Resin for 30 minutes with end-over-end mixing, then washed 3 x in 1 mL washing buffer (50 mM HEPES pH 7.3, 150 mM NaCl, 10 mM imidazole) by centrifuging at 1000 r.p.m. for 2 minutes and aspirating washing buffer. Bound proteins were eluted in 500 uL of elution buffer (50 mM HEPES, 150 mM NaCl, 300 mM imidazole) for 10 minutes and the eluate was collected by centrifugation at 1000 r.p.m. for 2 minutes.

Purified, recombinant nanobodies were then functionalized for loading on Platinum chips. First, nanobodies were functionalized with azide using Sortase A mediated transpeptidation^45,46^. Sortase A reactions consisted of 5-10 µg of the purified nanobody protein, 1 µM Sortase A tetramutant^45^, and 1 mM 3-azido-1-propylamine^47^ (Sigma Aldrich 762016) in 1 x Sortase A buffer^46^ (50 mM Tris-HCl (pH 7.5), 10 mM CaCl_2_, and 150 mM NaCl). Reactions were incubated at 37 °C for 1 hour. Following the Sortase A reaction, recombinant protein was purified on Magne HaloTag Beads (Promega G7281). The beads were first equilibrated in equilibration buffer consisting of 50 mM HEPES, 150 mM NaCl, and 0.005 % Igepal CA-630 (pH 7.3). The azide-labelled proteins were added and incubated for 1 hour at room temperature. The beads were then washed 4 × 200 µL equilibration buffer, and the conjugation reaction was set up consisting of 0.5 µL 250 µM cetyltrimethylammonium bromide (CTAB), 2 µM DBCO loading complex (K-linker, Quantum-Si) in equilibration buffer. The K-linker loading complex consists of a streptavidin molecule coupled to a DNA oligonucleotide by a bis-biotin moiety. The DNA oligonucleotide is linked to a Cy3 fluorophore and DBCO molecule for click chemistry. The click conjugation reaction was incubated overnight at 37 °C, followed by 4 x washes in 200 µL equilibration buffer to remove unconjugated DBCO complex. Finally, the barcoded nanobodies were eluted from the beads by digestion with 1.5 µL HaloTEV protease (Promega G6601) in equilibration buffer supplemented with 1 mM dithiothreitol (DTT). Digests were incubated, in 1.5 mL Eppendorf Protein LoBind Tubes, for 1 h at 37 °C with mixing at 700 rpm. Eluted loading complexes were quantified using Cy3b fluorescence on a CFX96 Touch Real-Time PCR Detection System with excitation at 515-535 nm and emission at 560-580 nm against a standard curve.

### Fluorescent labelling of Recombinant Ligands

We assembled a plasmid construct, pFluor^Halo^, to allow us to functionalize recombinant protein ligands with Quantum-Si dye complexes^27^ (Extended Data Fig. 1d, Supplementary Information). We modified pPBC^Halo^ by replacing the peptide barcode and Sortase A motif with a Bis-Avitag sequence^48^ for labelling with a streptavidin fluorescent dye construct. We retained a dual *Esp*3I site for insertion of the ligand gene. A *GFP* gene was amplified with compatible restriction sites from a gene fragment encoding a Halo-GFP fusion gene by PCR using Q5 Hot Start Polymerase (NEB M0493L) under the following conditions: For a 50 µL PCR reaction, the components included 10 µL of 5X Q5 Hot Start Polymerase buffer, 1 µL of 10 mM dNTPs, 2.5 µL each of GFP_BbsI_F (10 µM) and GFP_BbsI_R (10 µM) primers, 5 µL of 5x Q5 High GC Enhancer, 0.5 µL of Q5 Hot Start High-Fidelity DNA Polymerase, and 10– 50 ng of template DNA. The PCR thermocycling conditions were as follows: initial denaturation at 98 °C for 30 seconds, followed by 25 cycles of denaturation at 98 °C for 30 seconds, annealing at 60 °C for 20 seconds, and extension at 72 °C for 30 seconds, with a final extension at 72 °C for 2 minutes. The amplified PCR product was purified with the NucleoSpin Gel and PCR Clean-up Mini Kit (Macherey-Nagel, 740609.50) to remove excess primers and nucleotides. The purified PCR product and the pFluor^Halo^ vector were digested with *Esp*3I (NEB R0734) at 37 °C for 1 hour, producing compatible ends for cloning. Both digested products were purified using the Nucleospin Gel and PCR Clean-up Kit (Macherey-Nagel) to eliminate reagents and small DNA fragments. Vector and insert were ligated using the T4 Rapid Ligase Kit (ThermoFisher, K1422) at a 1:7 molar ratio for 15 minutes at 25 °C, transformed into SHuffle T7 Express competent *E. coli* according to the manufacturers protocol and the new construct, pFluor^Halo^-GFP, confirmed by Sanger sequencing (Eurofins Genomics) (Extended Data Fig. 1d, Supplementary Information).

SHuffle T7 Express competent *E. coli* transformed with pFluor^Halo^-GFP plasmid were cultured overnight in LB broth with 100 µg/mL ampicillin at 30 °C with shaking at 180 r.p.m., followed by dilution to OD_600nm_ 0.1–0.2. After cell growth reached OD_600nm_ 0.4–0.6, protein expression was induced using 0.5 mM IPTG, and the incubation was maintained overnight at 30°C with shaking at 180 r.p.m. The recombinant protein was mixed with 50 µL Magne HaloTag Beads (Promega G7281) equilibrated in 50 mM HEPES (pH 8), 2 mM DTT, and 0.2 % Tween-20. After binding for 2 hours at room temperature with shaking at 700 r.p.m., beads were washed 3 x in 200 µL 50 mM HEPES (pH 8), 2 mM DTT, and 0.2 % Tween-20, the unbound supernatant was collected for analysis. Biotinylation of the recombinant protein was then carried out on the Halo beads using the BirA enzyme (Sigma CS0008). The reaction mixture contained 1 μL of BirA enzyme (1 mg/mL), 5 mM MgCl_2_, 150 µM D-biotin, and 5 mM ATP in equilibration buffer to a final volume of 50 μL. The reaction was incubated at 25 °C for 2 hours with shaking at 700 r.p.m. After incubation, the beads were washed twice with 200 µL equilibration buffer on a magnetic rack and resuspended in 50 uL of 50 mM HEPES pH 8, 250 mM KCl, 0.1% Tween-2. Biotinylated proteins were incubated with 1 µL of 100 µM Quantum-Si Sg4cy3 dye complex^27^ for 1 hour at 20 °C with shaking at 700 r.p.m. After the labelling reaction, the beads were washed to remove unbound dye, and the supernatant was removed before the elution step. Labelled GFP was eluted from the beads with 1.5 µL Halo-TEV protease (Promega G6601) in 1 x wash buffer (Quantum-Si) diluted with nuclease free water and supplemented with 1 mM DTT for 1.5 hours at 25 °C with shaking in an Eppendorf Thermomix at 180 r.p.m. Eluted labelled GFP was analyzed by electrophoresis on a 4-12% Bis-tris polyacrylamide gel (ThermoFisher NP0321BOX) an imaged using a Vilber Fusion FX6 instrument with C480 filter. Labelled GFP was quantified by measuring the Sg4Cy3 dye fluorescence with a Qubit fluorometer (ThermoFisher) using a blue excitation filter (475 nm), green emission (510–580 nm) and the red range (665–720 nm).

### Preparation of NAA Ligands for Dye-cycling

Three short peptides, each comprising four amino acid residues, were designed to label the N-terminus of GFP. We constructed pNAA^Halo^ to genetically encode NAA-GFP and allow recombinant expression and purification (Extended Data Fig. 3a-c, Supplementary Information). The vector pPBC^Halo^ and synthetic gene fragments encoding GFP with an N-terminal dye-cycling peptide sequence (either RLFA, FAQR or LARQ) separated by a GGGS linker, and C-terminal 2x FLAG tag (Extended Data 3a) were digested with *Bbs*I-HF (NEB R3539) and *Xho*I (NEB R0146) at 37 °C for 1 hour. The NAA-GFP was positioned immediately downstream of the SUMO tag to enable exposure of the desired NAA upon cleavage with UlpI. Digested products were purified using the NucleoSpin Gel and PCR Clean-up Mini Kit (Macherey-Nagel, 740609.50) and ligated into the pPBC^Halo^ vector using the T4 Rapid Ligase Kit at a 1:7 molar ratio of vector to insert at 37°C for 15 minutes.

The three pNAA^Halo^ plasmids were transformed into SHuffle T7 Express competent *E. coli* and expressed as for pFluor^Halo^-GFP. Recombinant NAA-GFP proteins were purified using 300 µL equilibrated Ni-NTA resin (Thermo Fisher Scientific 88222) for 30 minutes with endover-end mixing. Purified NAA-GFP was then attached to Magne Halo-Tag Beads (Promega G7281) as described above for 1 hour at room temperature, the beads were then washed 4 x in 200 µL equilibration buffer. The NAA-GFP was then eluted using SUMO protease, by incubating the beads with 1 µL UlpI protease (Invitrogen 12588018) in equilibration buffer supplemented with 1 mM DTT at 37 °C for 1 hour with shaking (Extended Data 3b,c). GFP fluorescence was quantified using a a Qubit fluorometer (ThermoFisher) with a blue excitation filter (475 nm) and green emission (510–580 nm).

To directly attach GFP ligands to the Platinum chip, we constructed versions of pNAA^Halo^ in which a Sortase A motif (LPETGG) was included in place of C-terminal 2x FLAG tag (Supplementary Information). To assemble these constructs, synthetic gene fragments encoding NAA-GFP (Supplementary Information) were amplified using PCR conditions described for pFluorHalo but with annealing at 70 °C for 20 seconds. The forward primers used to amplify the R-GFP, F-GFP, and L-GFP synthetic gene fragments were GFP_RLFA_BbsI_TGGT_F, GFP_FAQR_BbsI_TGGT_F and, GFP_LARQ_BbsI_TGGT_F respectively and GFP_BbsI_TTGC_R was used as a reverse primer for the three constructs (Supplementary Information). The amplified PCR products and the pPBCHalo vector were then digested with BbsI-HF (NEB R3539) and ligated as described above. Sortase A reactions were performed as above to functionalize NAA-GFP with C-terminal azide, purified on HaloTag magnetic beads, clicked to the loading complex (K-linker, Quantum-Si) (Extended Data Fig. 3d,e). Ligands were eluted with UlpI, simultaneously exposing the NAA for each ligand. This facilitated the functionalization of the pNAA^Halo^ GFP with azide, which was then used to click the complex (K-linker, Quantum-Si) as described above.

### Single-molecule binding kinetics on the Platinum system

Binding kinetics experiments consisted of a single recognition segment on the Platinum system of defined duration, with the run script executed on the Quantum-Si cloud platform. Following chip check, we followed the standard Platinum loading protocol. Briefly, flow cells were rehydrated by washing 3 times with 50 µL 70 % Isopropanol and 3 times with 50 µL 1 x Wash Buffer, diluted in nuclease free water (Quantum-Si). The chip was removed from the Platinum instrument and each flow cell was loaded for 15 minutes with 30 µL of barcoded nanobody coupled to the loading complex (K-linker, Quantum-Si), diluted in 1x wash buffer at a concentration 2-3 nM for V2 chemistry and 0.05-0.2 nM for V3 chemistry. Imaging Solution was prepared by mixing 2 x Wash Buffer, nuclease-free water, and additives Trolox, Gox, and Catalase following the manufacturer’s instructions. After loading, residual nanobody complexes were removed with 6 x washes with 1 x Wash Buffer, Imaging Solution (30 μL) was added to each flow cell, and the chip returned to the Platinum for loading quantification. For kinetic characterization, the Recognition Solution was prepared following the Quantum-Si protocol by combining 33 μL of 2 x Wash Buffer, 6.6 μL of Additive 1, 3.3 μL of Additive 2 and 3.3 μL of Additive 3. For direct labelling experiments, the recognition solution was supplemented with Sg4Cy3-labeled GFP. For dye-cycling experiments, the Recognition Solution was supplemented with a mixture of NAA-GFP (with the volume of nuclease free water reduced) and 8.8 µL of NAA recognizer mix (Reagent A, Quantum-Si). Excess imaging solution was aspirated, 30 µL of Recognition Solution added to the flow cells, and the chip was sealed and reinserted into the instrument. Platinum runs collected binding kinetics data for 2 h.

### On-chip proteolysis

After the binding kinetics run, the chip was removed from the Platinum instrument, washed four times with 1 x Wash Buffer, then incubated in 1 x Wash Buffer, supplemented with 1 mM DTT with 5 U UlpI protease (Invitrogen 12588018) per 70 µL reaction for 2 hours at room temperature. Following this on-chip digestion step, the chip was washed six times with 50 µL 1 x Wash Buffer, after which it was prepared for NGPS.

### Peptide sequencing using Quantum-Si technology

NGPS was performed according to Quantum-Si’s standard protocol, commencing the protocol after loading quantification. Recognition Solution was prepared according to the standard Quantum-Si protocol, 27 µL was applied to each flow cell of the chip, and the chip inserted into the instrument for peptide sequencing. After the 15-minute recognition step, the chip was removed, and 3 µL of Aminopeptidase solution was added to the bottom reservoir of each flow cell. A volume of 20 µL was carefully aspirated from the reservoir and mixed through the hole into each flow cell 10 times. NGPS data were analyzed using primary analysis v2.5.0 and peptide alignment v2.5.0 analysis tools on the cloud platform^34^, producing alignment data consisting of nanowell identification numbers and alignment information, including the residues that were detected and the alignment score.

### Binding kinetics data analysis

Initial pulse calling for all data was performed in real-time on the Platinum system^27^. Pulse data are then analyzed using the Quantum-Si primary analysis (v2.5.0). Resulting data were then retrieved and analyzed locally in a Juypiter environment using Python notebooks. We modified the qsi_algo python library^27^ for use with binding kinetics data^49^.

### Direct fluorescence

Nanowell properties were computed using the Quantum-Si cloud primary analysis workflow (v2.5.0). Nanowells were filtered by mean pulse duration (>0.3s), mean bin ratio (> 0.23, < 0.30), mean fluorescence intensity (> 35, < 100, arbitrary units), signal to noise ratio (SNR, ≥ 15), number of pulses (≥ 5 in 2 h period), and by bleach steps (= 1). Following nanowell level filtering, pulse data were retrieved for the corresponding nanowells. Pulses were filtered by pulse duration (≥ 0.3s), bin ratio (> 0.18, < 0.35), fluorescence intensity (> 40, < 120, arbitrary units). Following pulse filtering, inter-pulse durations were recalculated. We developed a filter to remove nanobodies with dissociation rates beyond the photobleaching rate, without knowledge of their dissociation rate. For each nanowell we computed the observed number of pulses, *P*_*obs*_, and calculated the expected number of pulses, *P*_*exp*_, given the mean pulse duration 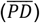 and mean inter-pulse duration 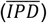:

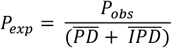

Nanobody variants where the median *P*_*obs*_: *P*_*exp*_ ratio per nan-owell was below 0.8 considered to be unmeasurable due to photobleaching and filtered out (Extended Data Fig. 2h,i).

For the remaining nanobodies, densities of PD or IPD were calculated for bins between 0 s and 100 s for PD, or 0 s and 1,000 s for IPD. Resulting PD and IPD densities were then fit using Scipy curve fit to a single-exponential decay model to determine k_*off*_ and k_*on*_, respectively.

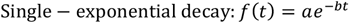

Dissociation constants, K_D_, were calculated according to the formula:

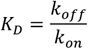

### Dye-cycling

First, data are analyzed using Quantum-Si’s primary analysis software, which segments trajectories into RSs and annotates them based on their pulsing and dye properties. We filter RSs for each NAA recognizer by the mean PD and IPD. Mean PD filtering parameters are 0.5 < R-GFP < 2.0 s, 0.7 < L-GFP < 3.0 s and 2.0 < F-GFP < 10.0 s. Mean IPD filtering parameters are 1.0 < R-GFP < 6.0 s, 6.0 < L-GFP < 20.0 s and 4.0 < F-GFP < 20.0 s.

We implemented a specific model to analyze dye-cycling data, which addresses systematic errors arising from missed events and dye pulsing kinetics, which impacts dissociation (k_*off*_) and association (k_*on*_) rate fitting. Unlike previous work^50^ where it was possible to define exact analytical solutions for probability densities that correct for missed events, we opted for a numerical solution to model dye-cycling.

We assume a two state Markov process in which the binder (in this instance a nanobody) is either bound or unbound by its ligand. The distributions of bound and unbound times are described by single-exponential distributions, parameterized by the association and dissociation rates, k_*on*_ and k_*off*_. Our model varies three parameters, k_*off*_, k_*on*_, and fraction skipped (f_*s*_). The third parameter, f_*s*_, allows the model to account for ligand binding events skipped arbitrarily, for instance due a failure of the RS caller to correctly classify the RS, exclusion of an RS due to data filtering steps, or if a ligand molecule does not correctly cycle the NAA recognizer. In these instances, the bound duration would be recorded as an interval between recorded ligand binding events. The model also utilizes the pulsing properties of the NAA recognizers, the total ligand (GFP) concentration, the p value set by the RS splitting software to split a protoRS, the buffer length used to split RS (see below), and the minimum pulses to constitute an RS. The RS calling algorithm creates a data-frame describing all RS observed in each experiment. This dataset is information rich, containing statistics for each RS on the nanowell, start, end and duration of each RS, as well as summary properties of the pulsing characteristics for each RS such as mean PD and IPD, and the associated dye properties, and an NAA annotation.

### 1. Dissociation rate

Three main sources of systematic error were identified in dissociation rate analysis, which could lead to signal distortion (Extended Data Fig. 4).

#### 1.1. Imprecision in the determination of bound times

When a ligand binds, a dye molecule is either be bound or unbound, with a certain probability given by the relationship between PD and IPD. Therefore, an RS may begin and end with a “bound” (B) or “unbound” (U) state, the four combinations thereof being B-B, U-B, B-U, and U-U. The probability of a ligand being in the bound state can be calculated from the means the pulsing parameters 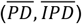:

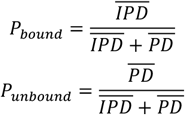

These probabilities are used to scale PDFs by the likelihood of observing bound/unbound dye combinations at the start and end of the RS. We correct for dye pulsing rate by modulating the expected rate-constant by the mean dye IPD.

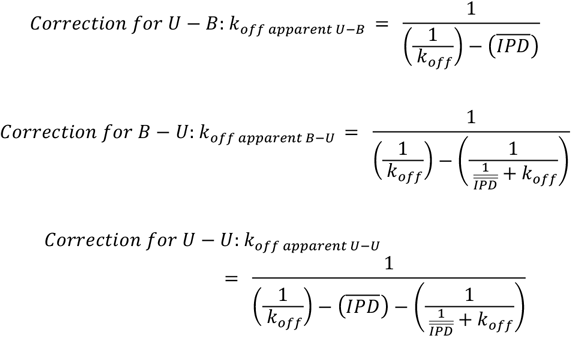

We then compute the probability density functions (PDFs) for the corrected apparent rates (k_app_) according to:

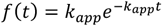

PDFs of apparent dissociation rates are scaled by the probability of observing each case, for instance, *P* _*B* −_*U* = *P*_*bound*_· *P*_*unbound*_, and summed, yielding the corrected probability density^49^.

#### 1.2. Missed unbound times, can occur due to high-frequency ligand (GFP) binding events, resulting in two independent ligand binding events being indistinguishable

This type of error is reduced by the use of multiple NAA ligand types, however we also provide a correction for consecutive binding events of the same NAA type. We provide a brief explanation of how the RS calling algorithm segments trajectories, for a fuller description see^34^. The RS caller first calls “protoRS”, defined as temporal regions of sustained pulsing, irrespective of other pulsing characteristics. ProtoRS are then separated into RS by applying a multiparametric statistical test on PD, IPD, bin ratio, intensity to candidate split points. A split point is any two consecutive pulses. A “buffer” of *α* pulses before and after each putative split point form each tested group. The algorithm iteratively applies the test to each potential split point within a protoRS. Here, we use a buffer size of *α* = 10. If the statistical test returns a P value < 0.0001 for the PD, IPD, bin ratio or intensity, the split point is considered valid, and the protoRS is split. If an RS is immediately followed by a binding event of the same type (e.g. R-GFP…R-GFP), in principle, protoRS would only split using the IPD parameter, as all other parameters are derived from the same distributions. If two binding events occur in rapid succession, such that the IPD of the dye pulsing and the interval between RS are likely to be derived from the same distribution, the test would not return a significant P value, and the protoRS would not be split.

To model splitting of RS, we estimate split points given the IPD of a dye. Naively, a single value (t_split_) can be computed using the following equation:

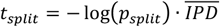

However, this does not accurately reflect the process of RS splitting, in which the RS caller samples *α* values from the IPD distribution, where *α* = buffer size. We can therefore model split times, given the P value, buffer size (*α*) and dye IPD as a gamma distribution to obtain the CDF the likelihood of splitting RSs:

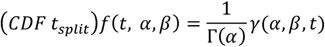

where

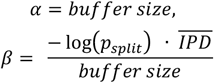

The likelihood of missing an interRS event between identical NAA ligand events is then (1-CDF *t*_*split*_). Multiplying the PDF of inter-RS events (single-exponential distribution parameterized by *k*_*on*_) by (1-CDF *t*_*split*_) gives the estimated PDF of missed inter-RS (*g*(*t*)). We estimate the probability of missing an interRS event as the integral of *g*(*t*). The distribution of two fused RS is then the convolution of (*f*(*t*) *\* g*(*t*) *\* f*(*t*)). The convolution is repeated n times. The resulting PDFs are then scaled by the respective probabilities of missing n RS. We scale the probability density from (1) by the probability of correctly terminating an RS, and sum with the scaled PDFs of fused events.

#### 1.3. Missed bound times

short ligand binding events are missed if the minimum of 3 pulses are not observed during the binding event. We correct for this by applying a loss function to the PDF of binding event durations. This loss function is the cumulative density function that describes 3 pulses, parameterized by the mean PD and IPD for each given dye. We use this distribution to infer the minimum likelihood of observing an RS of a given duration. We compute PDFs, parameterized by the reciprocal of the mean PD and IPD. Then we obtain the PDF of a three-pulse burst by convolving the PD (3 x) and IPD (2 x) PDFs. By integration we obtain the cumulative density, which we use to infer the likelihood of observing a binding event at each time point *f*(*t*). This is our loss function to correct for missed events. The PDF of ligand binding events is then obtained by multiplying the probability density from (1.2) by this loss function.

These processes result in a single probability density for RS durations. Distributions of RS durations are then fit to this probability density using the differential evolution algorithm from Scipy^51^, using an objective function computes the sum of the squared residuals.

### 2. Association rate

For the analysis of association rates, we apply corrections to three sources of systematic error (Extended Data Fig. 4).

#### 2.1. Imprecision in the determination of bound times

As for the dissociation rate fitting, NAA recognizer pulsing rates can distort the interval measured between RSs. We compute a correction for each pair of NAA recognizers preceding and following a binding interval. We compute the probabilities (P_bound_ and P_unbound_) that the NAA recognizer was bound or unbound as above and scale the distributions accordingly. We perform signal convolution to compute the probability density of concatenated signals. We compute this for all possible combinations of preceding and following NAA types. In the case that the preceding and following RS are identical, we first begin by calculating the distribution of observed intervals, which we calculate as above, using N*CDF t*_*split*_). We convolve this distribution with a single-exponential distribution parameterized by the mean IPD for the dye. Where the preceding and following RS are of different types, we assume that all inter-RS events are detectable, as the RS caller can split protoRS based on PD, fluorescence intensity, bin ratio and IPD. In this instance, we convolve a single-exponential distribution, parameterized by k_*on*_, with single-exponential distributions of each NAA recognizer IPD, parameterized by the reciprocal of the mean IPD. Resulting distributions are scaled by the product of the relevant probabilities P_bound_ and P_unbound_, summed to obtain a single density, and scaled such that 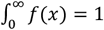.

#### 2.2. Missed binding events: short or missed binding events will be recorded as inter-RS durations

These timings correspond to the concatenation of n-1 missed events, together with n intervals. Such timings will be mixed with valid binding intervals and appear as a secondary component in histograms of inter-RS events. The correction for these types of events is the convolution of their predicted PDFs, weighted by the probability of their occurrence. We compute the PDF of skipped short events in a similar way to the correction for short events above (section 1.3). We calculate the cumulative density of three-pulse RSs *f*(*t*). The PDF of ligand binding events is computed as above 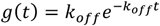. The PDF of skipped events is then given by *RS*_*miss*_ (*t*) = (1 − *f*) · *g*(*t*). We then compute the PDF of inter-RS events as a single-exponential distribution parameterized by k_*on*_. The PDF of skipped events can then be obtained by convolution of PDFs for combinations of n skipped events or f_s_ events with n+1 inter-RS events. The probability f_s_ is selected by the objective function between 0-1, and the probability of skipping a short RS (*p*_*miss*_) is given by the integral of *h*(*t*). We compute this correction for all possible combinations of short and f_s_ type event skipping. To limit computational time, we set an upper limit for event skipping at 10 concurrent skipped events. We find that for practical purposes, the probability of observing events this long is rather low, as it is the product of each skipped event probability (i.e. for 10 skipped f_s_ events, where typical fitted values are around 0.25, the probability would be f_s_ ^10^, which in practical terms is we estimate is in the order of 10^−6^ or below). We compute the probability of observing each combination as the product of the constituent skipped events, multiplied by the binomial coefficient (to scale for the number of possible combinations). To illustrate with an example: for *n* short events and *m* f_s_ events, the probability is 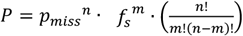. We then scale the probability density obtained by signal convolution by the probability *P*.

#### 2.3. Missed unbound events

RS splitting depends on a statistically significant difference in the properties of events either side of the split point. If the inter-RS duration is too brief, and each binding event is mediated by the same NAA-ligand, the ligand binding events will appear as fused RS. We correct for the expected loss of short inter-RS binding events in an analogous way to section 1.2 of the dissociation rate model. Once we obtain the PDF of t_split_, we can compute t_miss_ (1-t_split_ CDF) and apply it as a loss function by multiplication, resulting in a single probability density for IRS durations. For python code detailing these corrections see [49].

Distributions of IRS durations were then fit to these PDFs using the differential evolution algorithm from Scipy^51^ using an objective function computes the sum of the squared residuals. We use a global objective function which optimizes both k_*on*_, k_*off*_ and f_s_ simultaneously. For a given set of parameters, the objective function calculates the sum of the squared residuals for both RS and IRS distributions to their respective probability densities, then sums and returns a single value.

##### Linking binding kinetics data to NGPS barcode alignments

Platinum data are digitally addressed, such that each nanowell on a given chip is associated with both spatial coordinates and a unique identification number. To couple the data sets, we merge data frames containing binding kinetic information with NGPS alignments. First, binding kinetics data are filtered (see above for filtering parameters for direct fluorescence and dye-cycling experiments), resulting in valid binding kinetics data. Alignments from Platinum are filtered with the following parameters: alignment score >=4, read length >= 5, and the first reference residue must have been recognized. Filtered data are merged into a unified data frame using Pandas merge function.

To filter cross-talk between binders due to multiple loading of nanowells (Supplementary Information), we developed methods to filter binders which could not be distinguished from the non-GFP binding nanobody, LaM1, comparing statistics for each nanowell. For direct fluorescence experiments, we calculate the ratio between the observed and expected number of pulses for each nanowell (as for single nanobody experiments, see above). For dye-cycling experiments, we calculate the dye-cycling efficiency, the ratio of pulses within RSs to the total number of pulses per nanowell. We then performed Student’s t-tests comparing these data for each anti-GFP nanobody with LaM1. Variants where p > 0.01 were filtered from the analysis. Data for each variant passing filtering were pooled and fit either to single-exponential distributions (for direct-fluorescence) or the dye-cycling model.

**Extended Data Figure 1.**
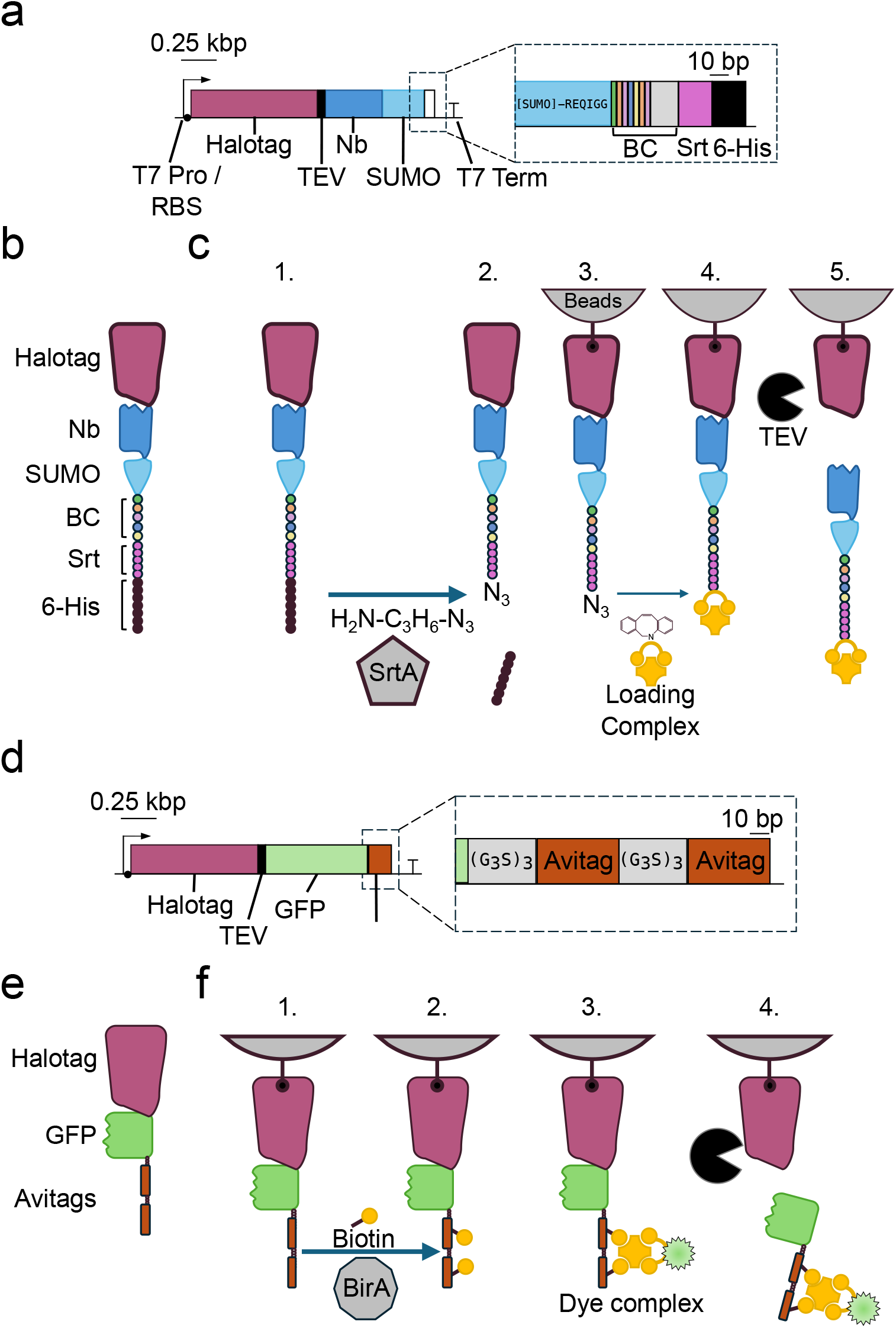
Construction of nanobodies and ligands. a) Schematic shows the expression cassette from pPBC^Halo^ for nanobody barcoding and recombinant expression in *E. coli*. Expanded region shows the C-terminus of SUMO (sequence shown), immediately followed by the protein barcode sequence, Sortase A motif (LPETGG), and a six-histidine tag. b. Schematic of recombinantly expressed protein. c. Schematic depicts the preparation of nanobodies for loading on chip. 1. Expressed nanobody proteins are initially purified using Ni-NTA affinity resin. 2. Sortase A mediated transpeptidation reaction functionalizes the C-terminus with an Azide (N_3_) motif. 3. Recombinant proteins are bound to Halotag magnetic beads to remove excess azide and Sortase A enzyme. 4. Loading complex is coupled to the C-terminus by copper-free DBCO-click chemistry. 5. Excess loading complex is removed and the loading product is eluted by proteolysis with TEV. d) pFluor^Halo^ construction for assembly of ligands in direct fluorescence experiments. e) Schematic of recombinantly expressed ligand protein. f) Ligand labelling scheme. 1. Expression and attachment to HaloTag magnetic beads. 2. Enzymatic biotinylation using BirA. 3. Labelling with fluorescent dye-complex compatible with 532-nm pulsed laser on Platinum. 4. Elution of fluorophore-labeled GFP from beads using TEV protease.

**Extended Data Figure 2.**
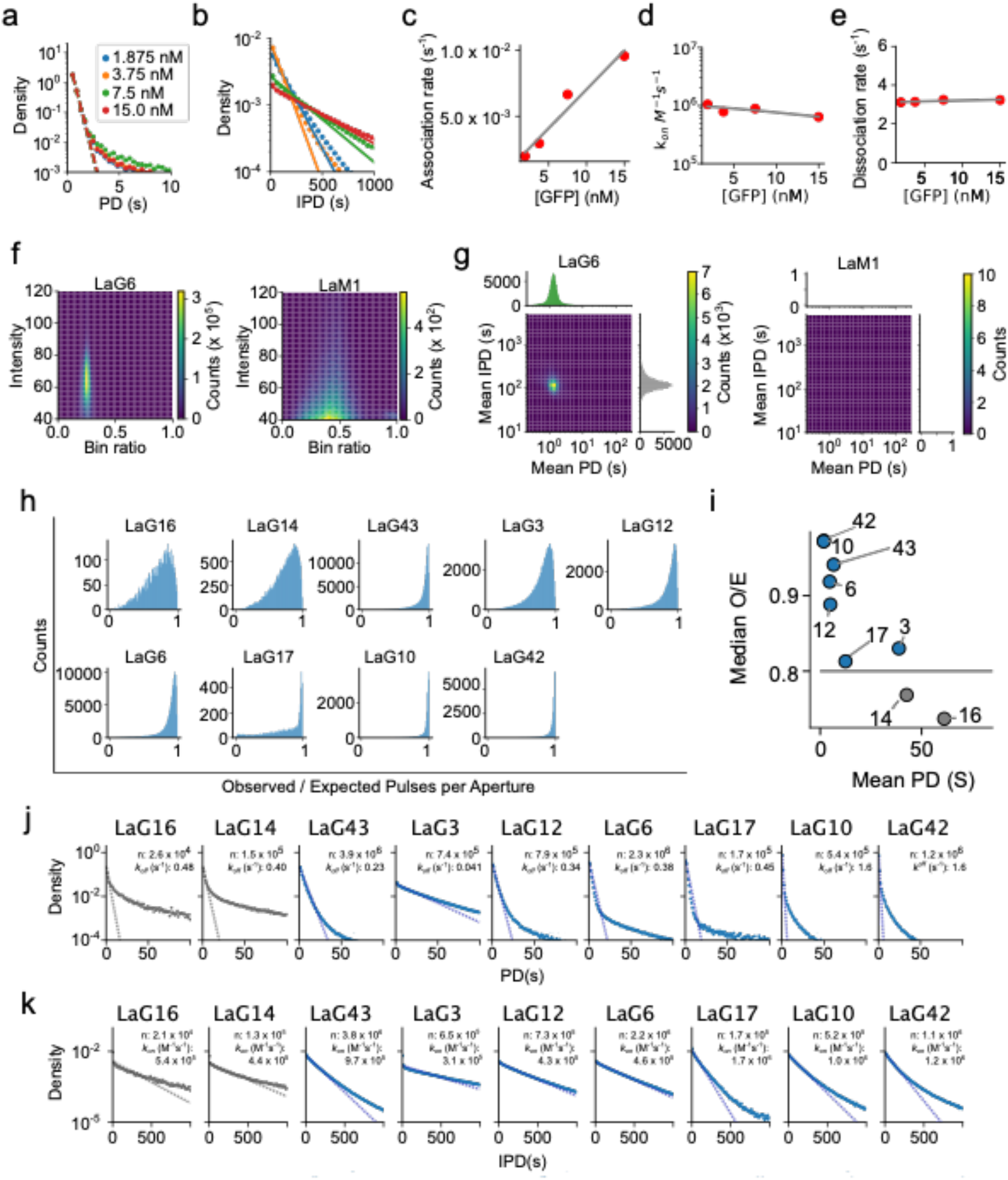
Direct fluorescence SM binding kinetics measurement on Platinum. a-e) SM binding kinetics of LaG42 with variable GFP concentration. One chip was loaded with LaG42 on each side and run two times. First with 1.875 nM (left) and 7.5 nM (right) GFP. In the second run the chip was washed and 3.5 nM and 15 nM GFP was added. a) PD histograms at each concentration. Dashed line indicates fit to a single exponential. b) IPD histograms at each concentration of GFP. Dashed line indicates fit to a single exponential. Legend as for a. c-e) Plots of first order association rate (s^−1^) (c), second order association rate constant, (k_*on*_, M^−1^s^−1^) (d) and dissociation rate (k_off_, s^−1^) (e) as function of GFP concentration. Lines indicate fit by linear regression. d) Second order association rate constant, k_*on*_ (M^−1^s^−1^), plotted as function of ligand concentration. Line indicates fit by linear regression. e) Dissociation rate, k_*off*_, as function of ligand concentration. Line indicates fit by linear regression. f) Unfiltered pulsing data for anti-GFP LaG6 and anti-mCherry LaM1 nanobodies with 7.5 nM GFP. 2D histogram shows pulse bin ratio (fluorescence lifetime) and intensity. g) 2D histograms show mean PD and IPD per nanowell for LaG6 and LaM1 with 7.5 nM GFP, after nanowell-filtering. Associated histograms show distributions of nanowell-mean PD (green) and nanowell-mean IPD (grey). h) Photobleaching filter: nine nanobody variants were measured separately with 7.5 nM GFP. Histograms show the distribution of the observed:expected number of pulses per nanowell for each nanobody. Expected number of pulses is calculated as run length/(mean PD + mean IPD). Photobleaching will result in an apparent deviation from a two state Markov process, and the ratio will deviate from 1. i) Correlation between median observed:expected number of pulses and mean PD. Gray horizontal line indicates photobleaching cut-off. Blue data points indicate binders that passed filters. j) Histograms of PD, aggregated over all filtered nanowells. Blue dashed line indicates fit to a single exponential. The number of pulses, n, and the calculated k_*off*_ values are indicated. k) Same as j but for IPD. Dashed line indicates single fit to a single exponential.

**Extended Data Figure 3.**
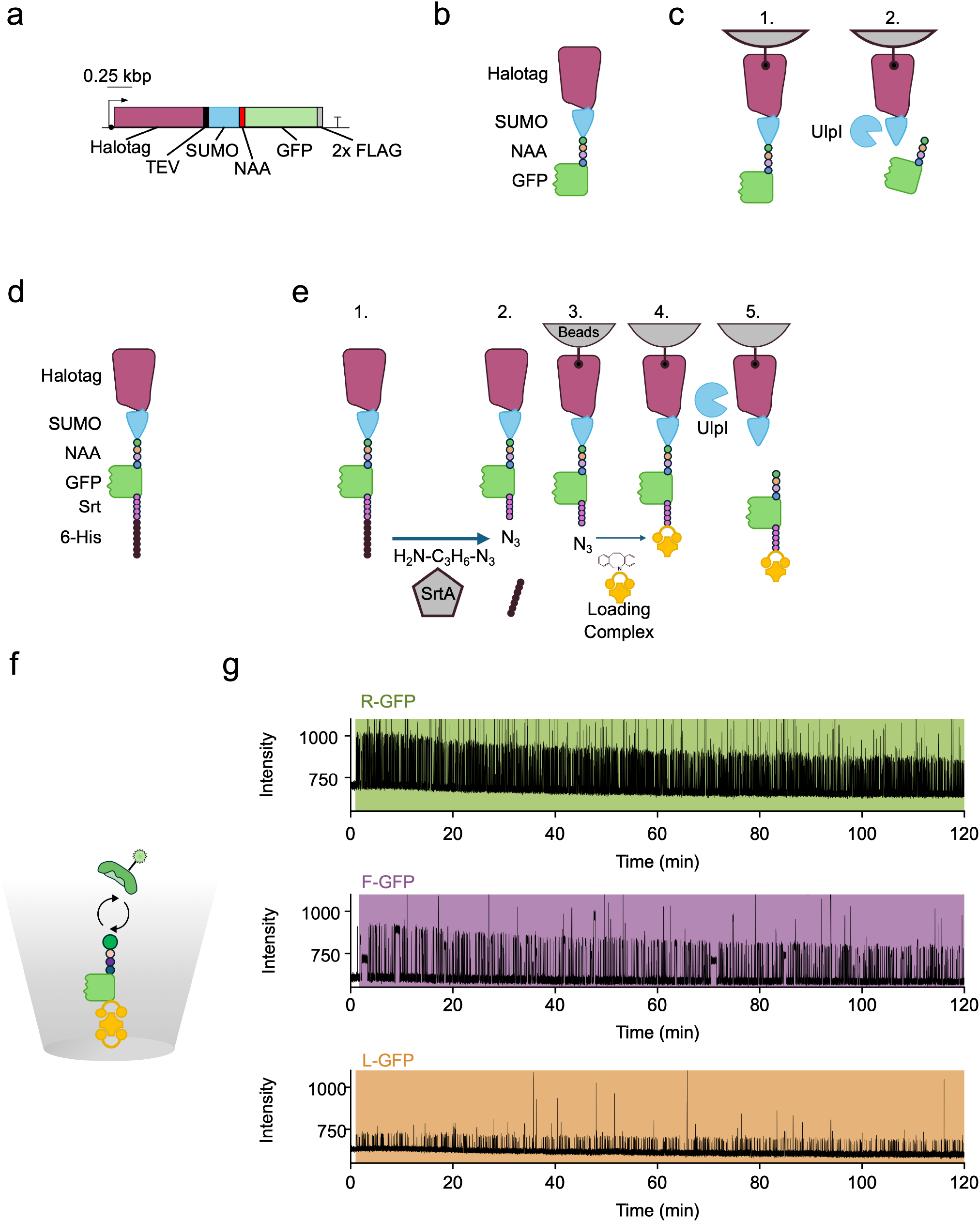
Dye-cycling on NAA-GFP ligands. a) Schematic of pNAA^Halo^ for recombinant expression of dye-cycling ligands. b) Schematic of recombinant NAA-GFP ligand. c) Preparation of NAA-GFP ligands. 1. Recombinant protein is expressed in *E. coli* then purified on Halotag magnetic beads. 2. Ligands displaying a distinct (R, L or F) N-terminal residue are eluted using UlpI protease. d) Schematic shows recombinant NAA-GFP-Sortase ligands for immobilization on chip. e) workflow to prepare NAA-GFP-Sortase for loading on chip. 1. Ligand is expressed and purified using Ni-NTA affinity resin. 2. Recombinant proteins are azide-functionalized at the C-terminus by Sortase A mediated transpeptidation. 3. The loading complex is clicked by azide-DBCO copper free click chemistry. 4. Excess loading complex is removed by washing and 5. Loading-ready complex is cleaved, exposing desired NAA with UlpI. f) Schematic of experiment. NAA-GFP-Sortase is loaded onto the Platinum chip and NAA recognizer mix flowed in. g) Trajectories for each R-GFP, F-GFP and L-GFP are annotated by the primary analysis algorithm for their respective RS.

**Extended Data Figure 4.**
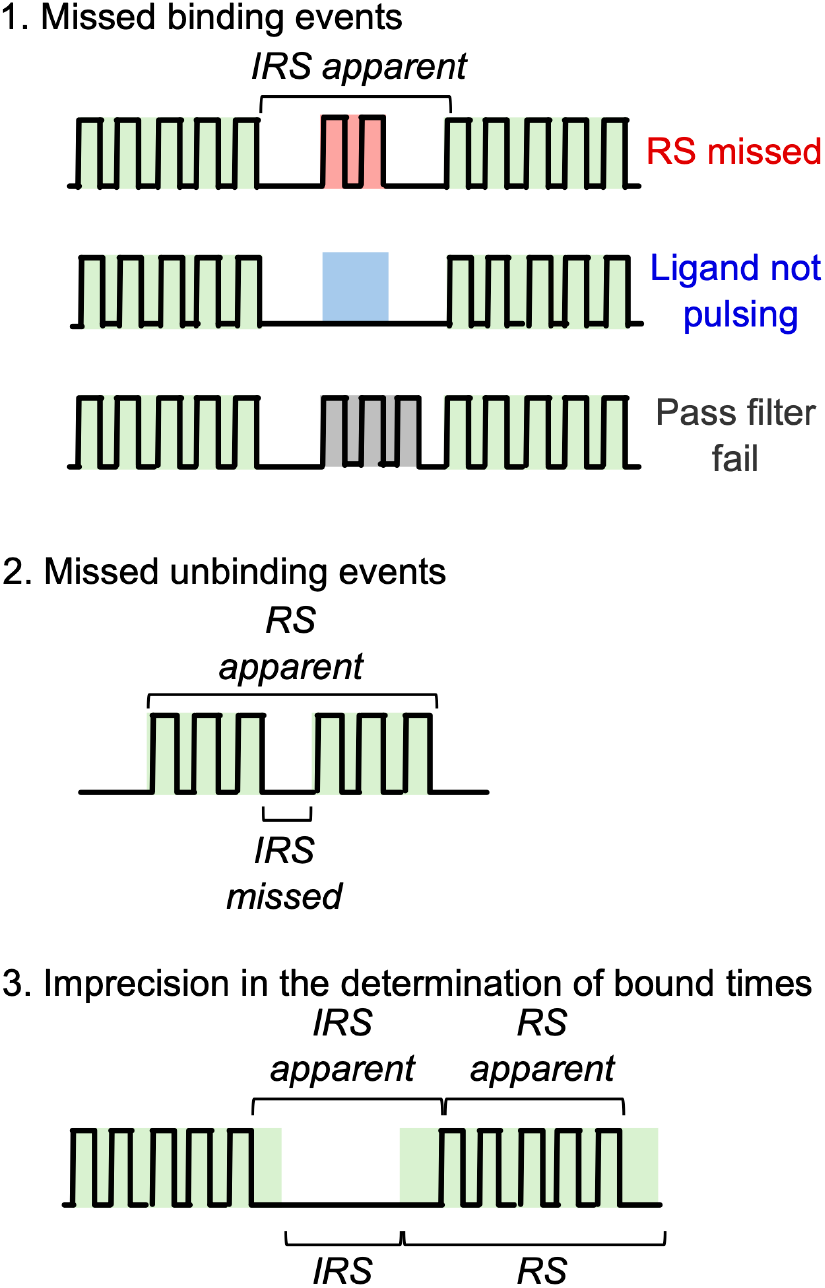
Dye-cycling model. Types of missed event and corrections accounted for in the dye-cycling model. Ligand binding events are shown as colored boxes and black lines depict NAA recognizer pulses. Green binding events are recognized as RSs. Red, blue and grey events are not recorded by primary analysis software.

**Extended Data Figure 5.**
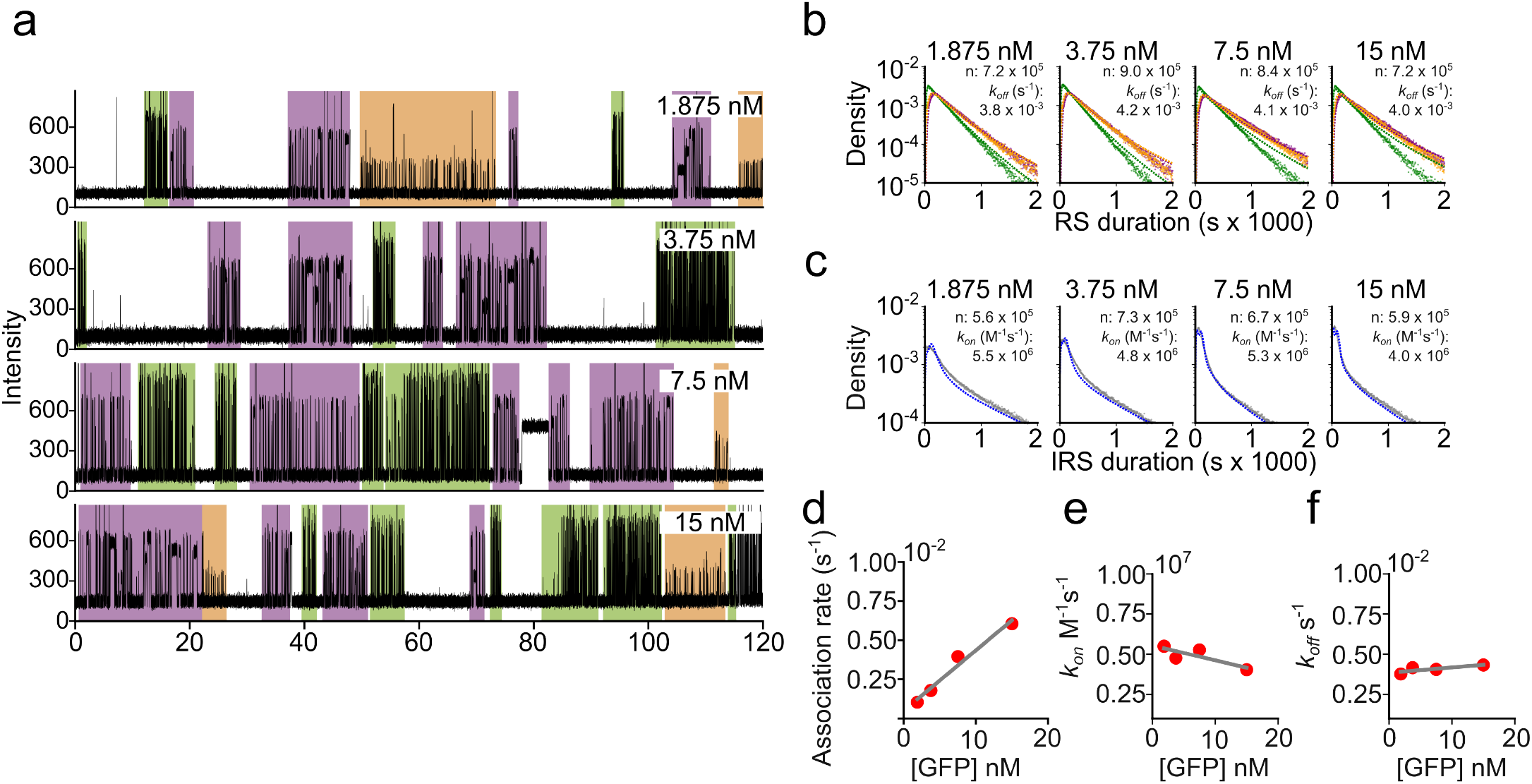
Characterization of dye-cycling over a range of ligand concentrations. a) Trajectories show dye-cycling of various concentrations of NAA-GFP on the anti-GFP nanobody LaG16. Pulses are shown as black traces and RS annotations as colored boxes. R-GFP: green, L-GFP: orange, F-GFP: Purple. b) Histograms of RS durations for each NAA-GFP concentration and type. Coloring as in (a). Dashed lines indicate fit to the dye-cycling model. The number of RS, n, and the calculated k_*off*_ values are indicated. c) Histograms of IRS duration histograms for each NAA-GFP concentration. Blue dashed line indicates fit to the to the dye-cycling model. The number of IRS, n, and the calculated k_*off*_ values are indicated. d-f) Plots of first order association rate (s^−1^) (d), second order association rate constant, (k_*on*_, M^−1^s^−1^) (e) and dissociation rate (k_off_, s^−1^) (f) at each concentration of NAA-GFP. Lines indicate fit by linear regression.

**Extended Data Figure 6.**
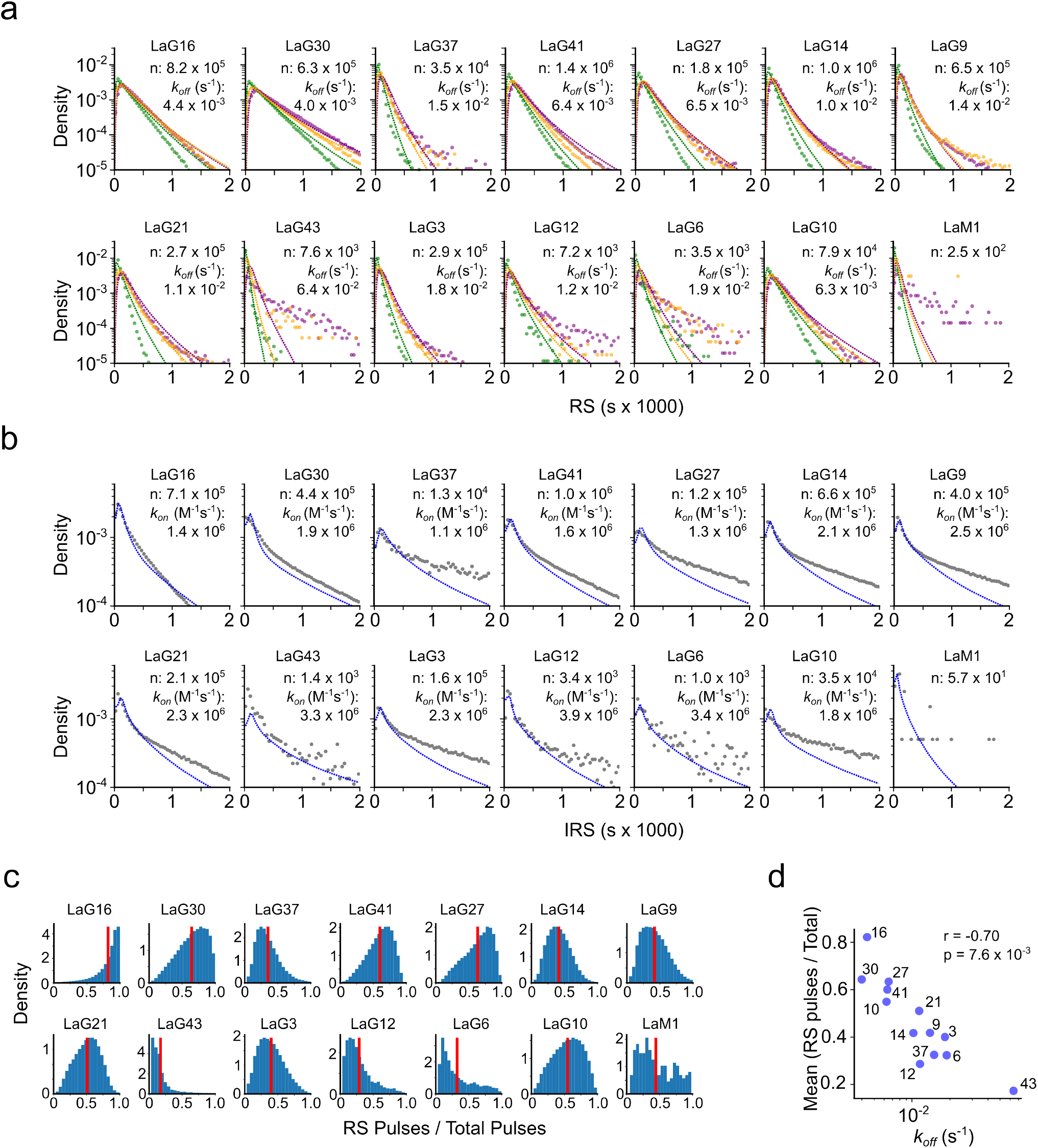
SM binding kinetics of 13 individual anti-GFP nanobodies by dye-cycling with 7.5 nM NAA-GFP. a) Histograms of RS distributions for each anti-GFP nanobody and the anti-mCherry nanobody LaM1 for each NAA-GFP type. R-GFP: green, L-GFP: orange, and F-GFP: Purple. Dashed lines indicate fits to the dye-cycling model. The number of RS, n, and the calculated k_*off*_ values are indicated. b) Histograms of inter-RS durations for the same nanobodies as in (a) (gray points). Blue dashed lines indicate fits to the dye-cycling model. The number of IRS, n, and the calculated k_*off*_ values are indicated. c) Establishment of metric for dye-cycling efficiency, using single nanobody runs. For each nanowell, we compute the ratio of RS pulses:total pulses. Higher values indicate more efficient recording of each binding event. Lower values indicate a larger relative fraction of binding events are missed. Histograms show distributions of this efficiency metric over valid nanowells. d) Correlation between dye-cycling efficiency (mean RS [pulses:total pulses]) and koff measured by dye-cycling.

**Extended Data Figure 7.**
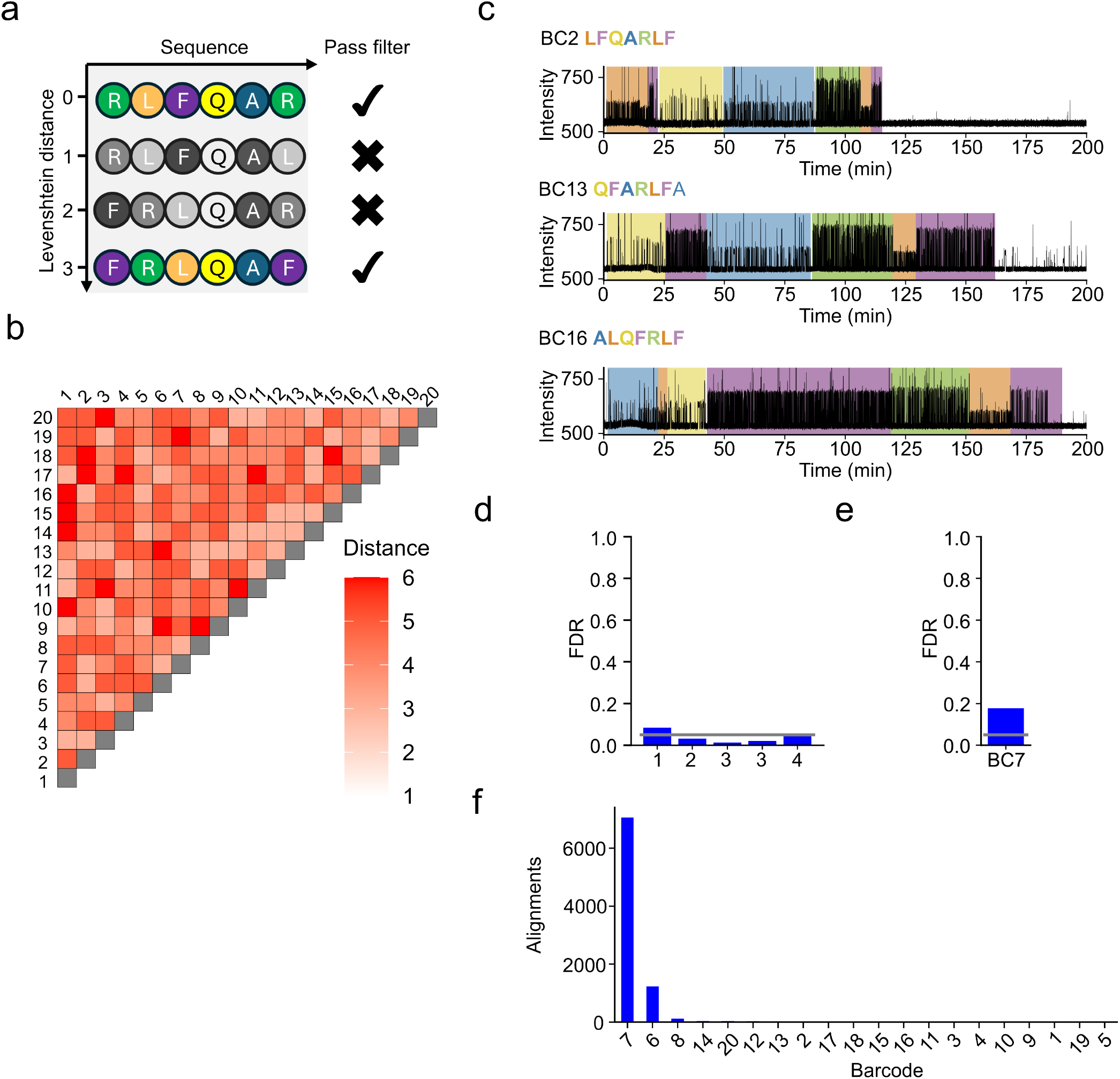
Single molecule NGPS decoding of protein barcodes. a) Design space for error resistance in protein barcodes. Levenshtein distances between sequences are used to construct peptide barcode sets. All barcodes in the set are separated by a distance ≥3. The cartoon depicts four putative barcode sequences. The first barcode (RLFQAR) is the seed barcode for the set, and the remaining barcodes are placed by their Levenshtein distance relative to the seed barcode. Barcodes with Levenshtein distances <3 are not included in the set (grayed sequences), whereas FRLQAF, with Levenshtein distance ≥3, in color, is added to the set. For more details on protein barcode design see [28]. (b) Heat map showing all-versus-all Levenshtein distances between the sequences for the 20 barcode sequences in this study. c) example trajectories for a subset of peptide barcodes. d) False discovery rates (FDR), the ratio of on-target to off-target alignments, calculated for sub-sets of the 20 protein barcodes. Set 1: BC7, BC13, BC17, BC1, BC9. Set 2: BC5, BC9, BC6, BC16, BC15. Set 3: BC2, BC16, BC3, BC13, BC11, BC14, BC15, BC20, Set 4: BC16, BC12. e) FDR for BC7 loaded alone. f) Number of alignments to all 20 barcode sequences for BC7 loaded alone.

**Extended Data Figure 8.**
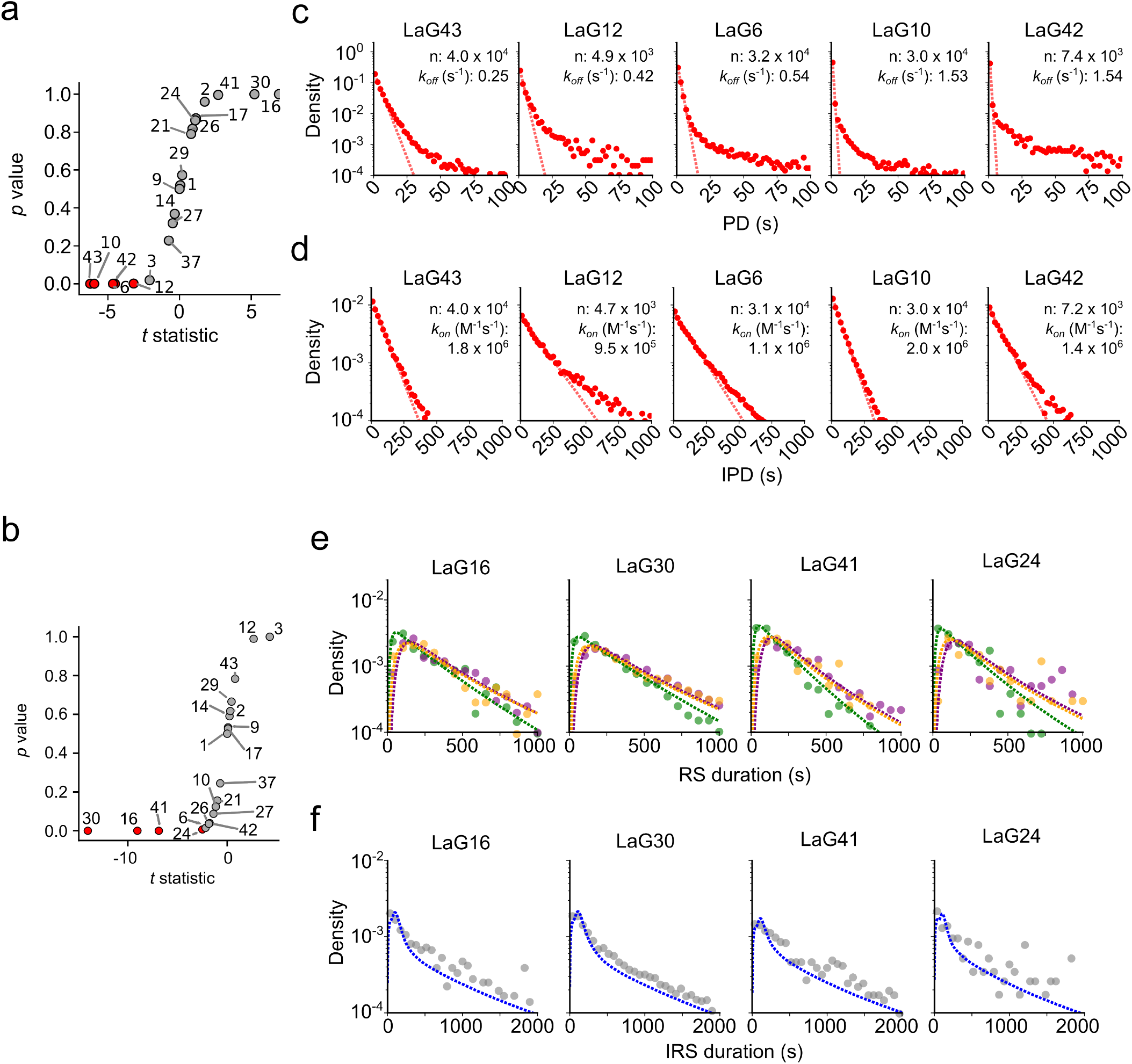
Parallel decoding of nanobody binding kinetics by direct fluorescence and dye-cycling for a 20-barcode set, with 7.5 nM GFP. a) Filtering of the direct labeling experiment. We calculate the ratio between observed and expected number of pulses per nanowell for each anti-GFP nanobody (see Extended Data Fig. 2h, i, Supplementary Information and Methods), and perform a one-sided Student’s t-test to test whether there is a statistically significant difference compared the data from the anti-mCherry nanobody LaM1. Binders passing the filter (p < 0.01) are indicated in red. b) Filtering of the dye-cycling experiment. Result of Student’s t-test comparing each anti-GFP variant to LaM1. We compute the dyecycling efficiency, the ratio of RS pulses to total pulses per nanowell (see Extended Data Fig. 6c, d) for each anti-GFP nanobody and compare to the dye-cycling efficiency metric for LaM1 nanowells (see Methods). Binders passing the filter (p < 0.01) are indicated in red. c) Direct fluorescence PD histograms for nanobodies passing filter. Dashed line indicates single-exponential fit to obtain k_*off*_. d) Direct fluorescence IPD histograms for nanobodies passing filter. Dashed line indicates single-exponential fit to obtain k_*on*_. e) RS duration histograms for each NAA-GFP type for nanobodies passing filter. Dashed lines indicate fits to the dye-cycling model to derive k_off_. R-GFP: green, L-GFP: orange, F-GFP: purple. f) Inter-RS duration histograms for nanobodies passing filter. Blue dashed lines indicate fits to the dye-cycling model to derive k_*on*_.

