## Supplementary_Information for "Parallelization of single-molecule binding kinetic measurements via protein barcode sequencing"

### Supplementary text

#### Direct fluorescence binding kinetics characterization

We sought to confirm the concentration dependence of the association rate ($k_{on}$, $M^{-1}s^{-1}$), and independence of concentration on dissociation rate ($k_{off}$, $s^{-1}$) in direct labeling experiments. We therefore measured four concentrations (1.875, 3.75, 7.5 and 15 nM) of fluorescently labelled GFP interacting with a single nanobody, LaG42. We loaded a single chip with LaG42 on both flow-cells and measured binding interactions for 2 h at 25 °C with 1.875 and 7.5 nM GFP in each flow-cell, respectively. We then extensively washed the flow cells and measured binding interactions for a subsequent 2 h at 25 °C with 3.75 and 15 nM GFP. We fit single exponential decays to the distributions of PD and IPD to obtain rate constants (Extended Data Fig. 2a,b). Comparison of both first and second order association rates revealed a concentration dependence (Extended Data Fig. c,d). Similarly, analysis of measured $k_{off}$ values revealed it is independent of ligand concentration.

##### Static ligand Dye-cycling

To determine whether exposed NAAs are detected when presented on GFP ligands, we constructed recombinant NAA-GFP proteins, displaying either Arginine, Phenylalanine or Leucine N-terminal residues, and fused at their C-termini with Sortase A motifs to facilitate loading on chip. We expressed, purified and conjugated the recombinant NAA-GFP to a Platinum loading complex and directly captured on chip (Extended Data Fig. 3a-c, see Methods). We then incubated with NAA recognizers used in NGPS sequencing for 2 h at 25 °C. We observed RSs for all species lasting the entire 2 h experiment (Extended Data Fig. 3d), confirming that NGPS recognizers interact with NAA-GFPs.

#### Dye cycling concentration response

As for our direct labeling experiments, we sought to initially confirm that dye-cycling faithfully reported binding kinetics by demonstrating the concentration dependence of the association rate, and independence of the dissociation rate. We loaded a high-affinity nanobody, LaG16, on Platinum and measured interactions by dye-cycling with 1.875, 3.75, 7.5 and 15 nM NAA-GFP. Trajectories from single nanowells show repeated binding and unbinding of individual GFP molecules (Extended Data Fig. 5a). We fit distributions of RS and inter-RS durations with our dye-cycling model (Extended Data Fig. 4, see Methods) to obtain $k_{off}$ and $k_{on}$ rates for each concentration (Extended Data Fig. 5b,c), confirming expected concentration dependence of the association rate, and concentration independence of the dissocation rate, with similar values of $k_{on}$ and $k_{off}$ calculated for all concentrations (Extended Data Fig. 5d-f).

### Description of protein barcode characterization

To characterize the set of 20 barcodes, we initially ordered the designs as an oPool (IDT), constructed a plasmid pool, expressed as recombinant proteins at the c-terminus of a Halotag-SUMO fusion, with a Sortase A motif for barcode functionalization. NGPS library prep was performed on this pool and barcodes were loaded on Platinum and sequenced. We obtained 11,636 alignments from a single Platinum flow cell. Next, we individually assembled plasmids encoding peptide barcoded-nanobody variants and sequenced subsets of the 20 barcodes to establish false discovery rates (FDR) within the set, by calculating the ratio between on-target to off target alignments (Extended data 7d). Of the four subsets, three had FDR below 0.05, the remaining subset FDR was 0.084 (Extended data 7d). We investigated the reason for this, identifying a clash between BC6 and BC7, which share the first three residues (RLF). This was confirmed by loading BC7 alone, which reported an FDR of 0.175, with BC6 the major component (Extended data 7e,f). Notably, the peptide alignment algorithm is frequently able to correctly distinguish BC7 from BC6 by the PD at a single residue. In BC6 ala in position 5 (RLFQALQPHDDDLPETG) has an expected PD of 1.45 s, vs 5.64 s for BC7 ala in position 4 (RLFAQLFHTDDDLPETG). These results demonstrate both the ability of the peptide alignment algorithm[34] to identify barcode sequences, and that our barcode set can discriminate protein variants on Platinum.

### Parallel data filtering due to cross-talk

We quantified crosstalk arising due to multiply loaded nanoapertures. We mixed anti-GFP LaG30 fused to barcode 16 (LaG30-BC16) and anti-mCherry LaM1 fused to barcode 12 (LaM1-BC12)[29], measured GFP binding kinetics by dye-cycling with 15 nM ligand, then cleaved the nanobody and sequenced the barcodes. We obtained 4,781 and 815 LaG30-BC16 alignments and 2,902 and 754 LaM1-BC12 alignments for each of two replicates (SFig. 1a). After intersecting NGPS with binding kinetics data, we retained 2,822 and 483 alignments for LaG30, but just 124 and 14 alignments for LaM1, with an average of 2.1, and 0.1 RSs per alignment for LaG30 and LaM1, respectively. Ligand binding rates observed for each binder were near-identical (SFig. 1b,c), demonstrating that RSs associated to LaM1 were derived from double aperture loading with LaG30.

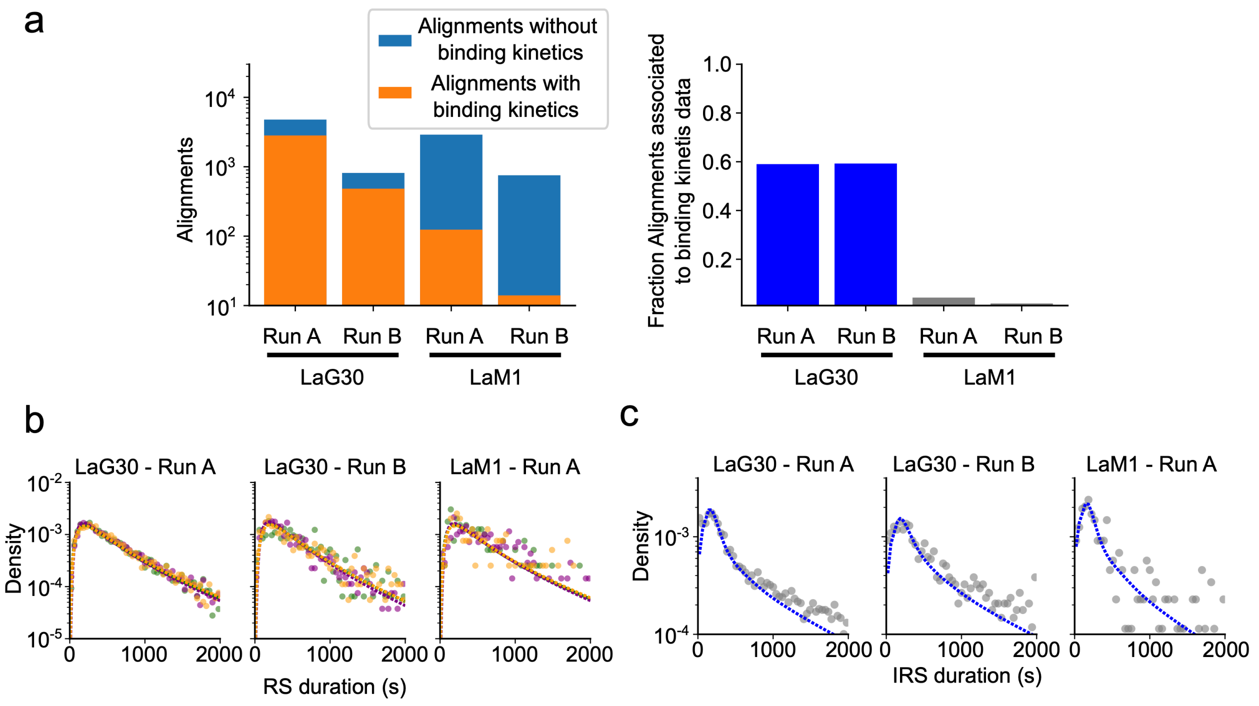

***Supplementary Figure 1: Characterization of parallel decoding of SM binding kinetics with the anti-GFP nanobody LaG30 and the anti-mCherry nanobody LaM1 measured using dye cycling*** *a) Alignment counts for parallel decoding of SM binding kinetics for LaG30 and LaM1. Left, NGPS alignments that were associated to binding kinetics data (orange) and NGPS alignments where binding kinetics data was not recorded (blue). Right, fraction of NGPS alignments associated to binding kinetics data. Data are from two runs. b) Histograms of RS durations for LaG30 and LaM1, from two runs. Insufficient data were retained after filtering for Run B - LaM1. R-GFP: Green, L-GFP: orange, F-GFP: purple. Dashed lines indicate fits to the dye-cycling model. c) Histograms of inter-RS durations for each nanobody. Dashed lines indicate fits to the dye-cycling model.*

In parallel experiments, we utilized the either the dye-cycling efficiency, or the obs/expected ratio for the number of pulses per nanowell to discriminate which variants could be differentiated from LaM1, which does not bind GFP. We calculate the respective metric for all nanowells, and apply a Student's t-test, comparing each variant to the non-GFP binding LaM1, which reports the average background from multiply loaded apertures on the chip (Extended Data Fig. 9b). If a binder produces signal significantly different from LaM1, it passes filtering and we analyse the binding kinetics.

### Barcode space expansion

We simulated peptide barcode sets using the current design methodology with a 6 NAA recognizer space to project how large a barcode set could reasonably be obtained. We construct the sets by computing all barcodes permissible under the stipulation that NAA recognizers cannot be repeated (i.e. L-L is not permitted), and that at least 5 recognizers must be used prior to repetition of any recognizer. There are 7.2 x $10^5$ combinations of NAA recognizers that satisfy these conditions with a barcode length of 8. Computing all-vs-all Levenshtein distances for this many barcodes is not computationally feasible, so we opted for a heuristic approach in which we randomize the entire barcode set and iteratively grow the barcode set, with each new candidate requiring a Levenshtein distance >2 with all other members in the set. Our results suggest that, with a barcode length of eight residues it is possible to construct barcode sets > 1,000 members (mean size of 10 sets: 1229) (Supplementary Figure 2). These barcode sets are not filtered by pulsing properties.

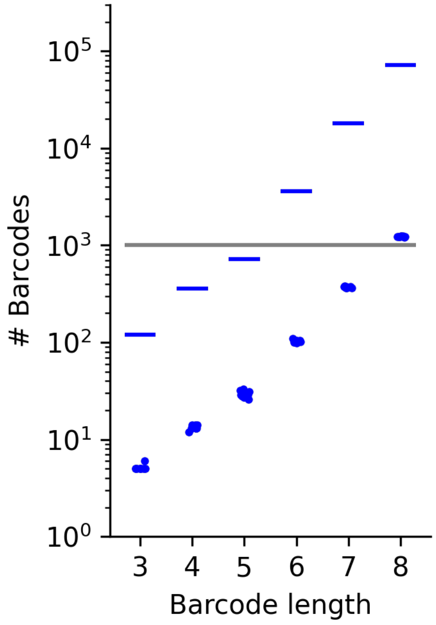

***Supplementary Figure 2: projected barcode sequence space with a 6 NAA recognizer mixture.*** *We simulated barcode sets with a Levenshtein distance > 2. Blue lines indicate the total size of the barcode space at each length. We created barcode sets by randomizing the barcode order, initializing a set with the first codeword and iteratively expanding the set, only adding new barcodes if they satisfy the Levenshtein distance > 2 for all current members. We repeated this process 10 times for each length. Grey line indicates 1,000 barcodes per set.*

### Supplementary methods

#### Kinetics characterization using Quantum-Si technology

The 20 barcoded nanobodies prepared in the previous steps were mixed at an equimolar ratio, and a 2-hour recognition experiment was planned on the cloud platform. The procedures were then performed according to the Quantum-Si protocol, as outlined below:

- The chip packaging was opened, and a dry chip check was performed using the Platinum device. The wash buffer was diluted in a 1:1 ratio with nuclease-free water.
- Each flow cell was washed twice with 50 µL of 70% isopropanol, with a 1-minute incubation between washes.
- Each flow cell was washed three times with 50 µL of 1X Wash Buffer. A volume of 50 µL of 1X Wash Buffer was added to each flow cell, allowing it to equilibrate between reservoirs for 30 seconds.
- The chip was inserted into the Platinum device, and a run plan was started.
- The sample was diluted to a concentration of 50-100 nM in 2X Wash Buffer. Subsequently, the Loading Solution for the barcoded nanobody mixture was prepared at a concentration of 0.05-5 nM in 1X Wash Buffer.
- The chip was removed from the Platinum device, and excess wash buffer was aspirated from the reservoirs.
- A volume of 30 µL of the Loading Solution was added to each flow cell and mixed 10 times. The mixture was incubated for 15 minutes.
- The Imaging Solution was prepared by adding 33 µL of 2X Wash Buffer, 15.4 µL of nuclease-free water, 8.8 µL of Additive 1, 4.4 µL of Additive 2, and 4.4 µL of Additive 3.
- After excess loading solution was removed from each reservoir, each side of the chip was washed six times with 50 µL of 1X Wash Buffer, alternating between the top and bottom reservoirs.
- A volume of 30 µL of Imaging Solution was added to each flow cell and mixed 10 times. The chip was returned to the instrument, and the run was continued.
- The Recognition Solution was prepared as follows:

**Note**: The Recognition Solution was prepared based on the type of kinetics characterization being performed. Reagent A was replaced with the ligand for the kinetics characterization experiment using a direct labelling approach, with a recommended final concentration of 1-100 nM (variable). The preparation details for each approach are shown in the table below.

| Component | General Protocol | Direct Labeling | Dye Cycling |
|---|---|---|---|
| **2X Wash Buffer** | 33 µL | 33 µL | 33 µL |
| **Nuclease-free water** | 4.4 µL | 4.4 µL | (Variable) µL |
| **Additive 1** | 6.6 µL | 6.6 µL | 6.6 µL |
| **Additive 2** | 3.3 µL | 3.3 µL | 3.3 µL |
| **Additive 3** | 3.3 µL | 3.3 µL | 3.3 µL |
| **Reagent A** | 8.8 µL | - | 8.8 µL |
| **GFP** | - | 8.8 µL | (Variable) µL |

- After the loading step was completed, the chip was removed from the Platinum device, and the excess Imaging Solution was aspirated.
- A volume of 27-30 µL of Recognition Solution was added to each flow cell and mixed 10 times.

- The plug was cleaned using isopropanol (IPA) on lint-free wipes followed by nuclease-free water. After cleaning, the surface interacting with the interior of the chip was not touched. The plug was then inserted and sealed gently using the plug pressure plate.
- The chip was returned to the instrument, the lid was closed, and the run continued.

### DNA Sequences

#### Oligonucleotides

**Supplementary Table 2. Oligonucleotides table**

| Primer | Sequence |
|---|---|
| GFP_BbsI_F | CGAAGAGAAGACctCGCTATGTCTAAAGGGGAAGAA |
| GFP_BbsI_R | CGAAGAGAAGACCTCACCCTTGTATAACTCATCCAT |
| Halo-SUMO_F | TTGCCTGAAACCGGCGGACA |
| Halo.SUMO_R | AGCGATTGCGTTGTCAGATTGGAA |
| Sanger_R | TAGAGGCCCCAAGGGGTTAT |
| N_GFP_F | gctaGAAGACctTGGT |
| N_GFP_R | TAACCTTACTCGAGTCATTT |
| GFP_RLFA_BbsI_TGGT_F | gctaGAAGACctTGGTCGTCTTTTTGCTGGTGGAGGCTCAATGTCTAAAGGGGAAGAA |
| GFP_FAQR_BbsI_TGGT_F | gctaGAAGACctTGGTTTTGCGCAGCGTGGTGGAGGCTCAATGTCTAAAGGGGAAGAA |
| GFP_LARQ_BbsI_TGGT_F | gctaGAAGACctTGGTTTGGCGCGTCAGGGTGGAGGCTCAATGTCTAAAGGGGAAGAA |
| GFP_BbsI_TTGC_R | tagcGAAGACctGCAACTTGTATAACTCATCCAT |

### Gene sequences

**Supplementary Table 3. Synthetic DNA gene fragments encoding Nb-SUMO-BC**

| Nanobody | Sequence |
|---|---|
| LaG2_BC4 | TTCCAATCTGACAACGCAATCGCTATGGCGCAGGTTCAACTGGTTGAAAGTGGGGGAGGGCTGGTACAGGCCGGAGGATCGCTTCGCTTATCTTGTGCGGCCTCGGGCCGTACTTTTTCCAATTATGCGATGGGGTGGTTCCGTCAAGCTCCCGGCAAAGAGCGTGAGTTTGTGGCCGCGATTTCTTGGACTGGTGTCTCAACGTACTATGCAGACTCCGTAAAAGGGCGTTTTACGATCTCTCGCGATAACGACAAGAACACGGTATATGTACAAATGAACTCTCTGATCCCTGAAGATACTGCCATCTATTATTGTGCGGCAGTCCGTGCGCGTTCTTTCTCAGACACTTACTCCCGTGTGAACGAATATGATTATTGGGGACAGGGGACCCAGGTGACCGTAGGTGGCGGATCTATGTCGGACTCAGAAGTCAATCAAGAAGCTAAGCCAGAGGTCAAGCCAGAAGTCAAGCCTGAGACTCACATCAATTTAAAGGTGTCCGATGGATCTTCAGAGATCTTCTTCAAGATCAAAAAGACCACTCCTTTAAGAAGGCTGATGGAAGCGTTCGCTAAAAGACAGGGTAAGGAAATGGACTCCTTAAGATTCTTGTACGACGGTATTAGAATTCAAGCTGATCAGACCCCTGAAGATTTGGACATGGAGGATAACGATATTATTGAGGCTCACAGAGAACAGATTGGTGGTCTGCAGCGTGCTTTCGCTCTGCATCCGGACGACGACTTGCCTGAaACCGGCGGACA |
| LaG3_BC20 | TTCCAATCTGACAACGCAATCGCTATGGCCCAGGTCCAGCTTGTGGAATCTGGCGGGGGTCTGGTCCAGGCTGGCGGGAGTCTTCGCGTTAGCTGTGCAGCTTCTGGTCGTACCTATTCTGATTATGCTATGGGTTGGTTCCGTCAAGCACCAGGCAAGGAACGTGACTTCGTAGCGGGGATCTCGGGTAGTGGCGGCGACACGTATTACGCAGACAGCGTGAAGGGCCGTTTTACGATCAGCCGTGATAATGCTAAGAACACGATGTACTTACAAATGAACTCTCTGAAGCCGGAGGATACCGCCGTATACTTCTGTGCGGCCCGCACTGGGACTGTTCTGTTCACCTCCCGTGTCGATTACCGTTATTGGGGTCAGGGTACACAGGTCACGGTCGGTGGCGGATCTATGTCGGACTCAGAAGTCAATCAAGAAGCTAAGCCAGAGGTCAAGCCAGAAGTCAAGCCTGAGACTCACATCAATTTAAAGGTGTCCGATGGATCTTCAGAGATCTTCTTCAAGATCAAAAAGACCACTCCTTTAAGAAGGCTGATGGAAGCGTTCGCTAAAAGACAGGGTAAGGAAATGGACTCCTTAAGATTCTTGTACGACGGTATTAGAATTCAAGCTGATCAGACCCCTGAAGATTTGGACATGGAGGATAACGATATTATTGAGGCTCACAGAGAACAGATTGGTGGTGCTCAGCGTCTGTTCCTGGCTCATCCGGACGACGACTTGCCTGAaACCGGCGGACA |
| LaG6_BC10 | TTCCAATCTGACAACGCAATCGCTATGGCCCAGGTGCAATTGGTCGAATCTGGTGGAGGGCTGGTGCAAGCGGGCGGATCTCTTCGTCTGTCATGCGCTGCATCCGGTCGCACGTTCTCGACCAGCGCAATGGCTTGGTTCCGCCAGGCACCTGGCAAGGAGCGTGAATTTGCCGCTGGTATCACTTGGATTAGCAGTTCAACGTACTATACGGATTCTGTTAAGGGCCGTTTTACGATCTCCCGCGACAACGCAAAAAATACCGTCTACCTTCAAATGAACTCATTGAAACCAGAAGATACCGCAGTCTACTATTGCGCCGCCAAGTCCGAAGGCTATTTTGGATTCCCCCGCGTTGAGAATGAATACCCGTATTGGGGACAAGGCACCCAAGTAACAGTAGGTGGCGGATCTATGTCGGACTCAGAAGTCAATCAAGAAGCTAAGCCAGAGGTCAAGCCAGAAGTCAAGCCTGAGACTCACATCAATTTAAAGGTGTCCGATGGATCTTCAGAGATCTTCTTCAAGATCAAAAAGACCACTCCTTTAAGAAGGCTGATGGAAGCGTTCGCTAAAAGACAGGGTAAGGAAATGGACTCCTTAAGATTCTTGTACGACGGTATTAGAATTCAAGCTGATCAGACCCCTGAAGATTTGGACATGGAGGATAACGATATTATTGAGGCTCACAGAGAACAGATTGGTGGTTTCGCTCAGCGTCTGTTCCAGGAACATGACGACGACTTGCCTGAaACCGGCGGACA |
| LaG9_BC14 | TTCCAATCTGACAACGCAATCGCTATGGCGGACGTACAGTTGGTTGAAAGCGGCGGTGGCCTGGTTCAAGCCGGGGGGAGTTTACGCCTTTCTTGTGCAGCGTCCGGGCGTACTTTCAGCACTAGCGCAATGGGCTGGTTCCGTCAGGCTCCTGGTAAGGAGCGTGAATTCGTAGCTCGTATTACTTGGTCTGCCGGGTATACAGCTTACTCAGATTCAGTTAAGGGCCGTTTCACCATTTCGCGTGATAAAGCAAAAAACACAGTTTATCTGCAGATGAATAGTCTTAAACCCGAGGATACTGCAGTATACTATTGTGCGTCACGCAGTGCAGGGTACTCCAGTAGTTTGACTCGCCGTGAGGACTACGCCTATTGGGGACAGGGAACTCAAGTAACTGTATCAGGTGGCGGATCTATGTCGGACTCAGAAGTCAATCAAGAAGCTAAGCCAGAGGTCAAGCCAGAAGTCAAGCCTGAGACTCACATCAATTTAAAGGTGTCCGATGGATCTTCAGAGATCTTCTTCAAGATCAAAAAGACCACTCCTTTAAGAAGGCTGATGGAAGCGTTCGCTAAAAGACAGGGTAAGGAAATGGACTCCTTAAGATTCTTGTACGACGGTATTAGAATTCAAGCTGATCAGACCCCTGAAGATTTGGACATGGAGGATAACGATATTATTGAGGCTCACAGAGAACAGATTGGTGGTCAGGCTCGTCTGTTCCTGCAGCCGCATGACGACGACTTGCCTGAaACCGGCGGACA |
| LaG10_BC19 | TTCCAATCTGACAACGCAATCGCTATGGCTCAGGTTCAGTTGGTTGAAAGTGGGGGCGGGCTGGTTCAGGCGGGTGATTCGTTACAACTTTCGTGTGCATTTAGCGGAGGAACCTTTTCAACCTATGCTATGGGGTGGTTCCGTCAAGCTCCGGGAAAGGAGCGCGAGTTTGTTGGAGGCATTTCGCG |

| | |
|---|---|
| | TTCGGGAGCGACAACAAATTACGAGGACAGTGTCAAGGGCCGCTTCACCATCTCAAAGGACAA<br>TACTAAGAACACAGTATATTTACAGTTAAACTCGCTTAAACCAGAGGACACTGCAGTGTATTATT<br>GTGCGGCTCGCAACAACATTTTACCAGTCACTACTATCGACAAGTACGAATACTGGGGTCAAG<br>GCACGCAGGTAACTGTCGGTGGCGGATCTATGTCGGACTCAGAAGTCAATCAAGAAGCTAAG<br>CCAGAGGTCAAGCCAGAAGTCAAGCCTGAGACTCACATCAATTTAAAGGTGTCCGATGGATCT<br>TCAGAGATCTTCTTCAAGATCAAAAAGACCACTCCTTTAAGAAGGCTGATGGAAGCGTTCGCTA<br>AAAGACAGGGTAAGGAAATGGACTCCTTAAGATTCTTGTACGACGGTATTAGAATTCAAGCTGA<br>TCAGACCCCTGAAGATTTGGACATGGAGGATAACGATATTATTGAGGCTCACAGAGAACAGAT<br>TGGTGGTGCTTTCCTGCGTCAGTTCCTGCATACTGACGACGACTTGCCTGAaACCGGCGGACA |
| LaG12_BC8 | TTCCAATCTGACAACGCAATCGCTATGGCCTCAGGTGCGGCGGGCGGAGGTCTTGGAGAGGG<br>ACTTGTACAAGCCGGTGGTAGCTTGCGCTTATCTTGTGCAGCTTCCGGTCGCACCTTTAATTC<br>CTACCCGATGGCATGGTTTCGTCAAGCTCCTGGAAAAGAACGTGAGTTTGTTGCCGCGTTGGG<br>CTGGTCCGGAGGGAGTACGGATTATGCGGACTCGGTAAAGGGCCGTTTTACTATTTCACGCG<br>ACAATTCGAAAAACACGGTATATTTAGAGATGAACTCGTTGAAACCGGACGACACTGGGGTTTA<br>CTACTGCGCCTTACGTCGTCGTGGCGGAGTGTACAACACCTATTCTGGAGAGAAAGATTACGA<br>CTACTGGGGGCAAGGCACCCAAGTGACAGTCAGTGGTGGCGGATCTATGTCGGACTCAGAAG<br>TCAATCAAGAAGCTAAGCCAGAGGTCAAGCCAGAAGTCAAGCCTGAGACTCACATCAATTTAA<br>AGGTGTCCGATGGATCTTCAGAGATCTTCTTCAAGATCAAAAAGACCACTCCTTTAAGAAGGCT<br>GATGGAAGCGTTCGCTAAAAGACAGGGTAAGGAAATGGACTCCTTAAGATTCTTGTACGACGG<br>TATTAGAATTCAAGCTGATCAGACCCCTGAAGATTTGGACATGGAGGATAACGATATTATTGAG<br>GCTCACAGAGAACAGATTGGTGGTCGTCAGTTCGCTCTGTTCCTGCATACTGACGACGACTTG<br>CCTGAaACCGGCGGACA |
| LaG14_BC1<br>1 | TTCCAATCTGACAACGCAATCGCTATGGCCCAAGTACAATTAGTTGAAAGTGGCGGAGGTTTA<br>GTCCAAGCTGGCGGTAGTCTGCGTTTGTCATGTGCAGCTAGTGGACGTACATACTCAATCTCT<br>GCCATGGGTTGGTTTCGTCAGGCACCAGGTAAAGAACGCGAGTTCGTAGCGGGCATTTCCCG<br>TTCCGGGGGGACGACTTACTACGCCGACCCTGTTAAGGGACGCTTCACGATTTCGCGCGACA<br>ATGCGAAAAATACCGTCTACCTGCAGATGAACAGCTTGAAACCAGAAGACACTGCCGTCTACT<br>ATTGTGCTGCTCGTGCCCGTGGCTGGACAACATTTCCTGCACGTGAGATCGAATACGATTATT<br>GGGGACAGGGCACACAGGTTACGGTCGGTGGCGGATCTATGTCGGACTCAGAAGTCAATCAA<br>GAAGCTAAGCCAGAGGTCAAGCCAGAAGTCAAGCCTGAGACTCACATCAATTTAAAGGTGTCC<br>GATGGATCTTCAGAGATCTTCTTCAAGATCAAAAAGACCACTCCTTTAAGAAGGCTGATGGAAG<br>CGTTCGCTAAAAGACAGGGTAAGGAAATGGACTCCTTAAGATTCTTGTACGACGGTATTAGAAT<br>TCAAGCTGATCAGACCCCTGAAGATTTGGACATGGAGGATAACGATATTATTGAGGCTCACAG<br>AGAACAGATTGGTGGTCAGCTGCGTGCTTTCCTGGCTCATCCGGACGACGACTTGCCTGAaAC<br>CGGCGGACA |
| LaG16_BC2 | TTCCAATCTGACAACGCAATCGCTATGGCCCAAGTACAGCTTGTCGAATCCGGCGGTCGCCTG<br>GTGCAAGCGGGCGATTCGTTGCGTTTAAGCTGTGCCGCCTCCGGTCGTACATTTTCGACGTCT<br>GCGATGGCGTGGTTTCGTCAAGCTCCTGGCCGCGAACGTGAGTTCGTGGCGGCCATTACGTG<br>GACTGTGGGGAACACAATCCTGGGGGATTCTGTGAAGGGTCGTTTTACAATTTCCCGTGACCG<br>TGCTAAAAACACCGTAGATTTGCAGATGGATAACCTGGAGCCAGAAGATACGGCCGTCTACTA<br>TTGCTCAGCCCGCTCGCGTGGTTATGTATTATCAGTCCTGCGTAGCGTTGATAGCTACGATTAT<br>TGGGGACAAGGGACTCAGGTGACAGTGTCAGGTGGCGGATCTATGTCGGACTCAGAAGTCAA<br>TCAAGAAGCTAAGCCAGAGGTCAAGCCAGAAGTCAAGCCTGAGACTCACATCAATTTAAAGGT<br>GTCCGATGGATCTTCAGAGATCTTCTTCAAGATCAAAAAGACCACTCCTTTAAGAAGGCTGATG<br>GAAGCGTTCGCTAAAAGACAGGGTAAGGAAATGGACTCCTTAAGATTCTTGTACGACGGTATT<br>AGAATTCAAGCTGATCAGACCCCTGAAGATTTGGACATGGAGGATAACGATATTATTGAGGCT<br>CACAGAGAACAGATTGGTGGTCTGTTCCAGGCTCGTCTGTTCCATACTGACGACGACTTGCCT<br>GAaACCGGCGGACA |
| LaG17_BC1 | TTCCAATCTGACAACGCAATCGCTATGGCCGACGTACAATTAGTAGAGAGCGGCGGTGGCTTG<br>GTCCAAGCGGGCGGCAGCTTACGTCTGAGCTGTGCAGCTAGTGGTCGCACTATCTCGATGGC<br>CGCTATGAGTTGGTTCCGTCAGGCACCGGGGAAAGAACGCGAGTTCGTTGCGGGTATTTCGC<br>GCTCCGCCGGTAGCGCGGTCCACGCCGACTCAGTGAAGGGCCGTTTTACGATTTCGCGCGAT<br>AACACCAAAAACACGTTGTACTTGCAAATGAACTCCCTTAAAGCGGAAGATACTGCTGTGTATT<br>ATTGTGCAGTACGTACCAGCGGTTTTTTCGGCTCCATCCCCCGTACCGGCACAGCCTTCGATT<br>ATTGGGGCCAGGGGACTCAAGTGACCGTATCAGGTGGCGGATCTATGTCGGACTCAGAAGTC<br>AATCAAGAAGCTAAGCCAGAGGTCAAGCCAGAAGTCAAGCCTGAGACTCACATCAATTTAAAG<br>GTGTCCGATGGATCTTCAGAGATCTTCTTCAAGATCAAAAAGACCACTCCTTTAAGAAGGCTGA<br>TGGAAGCGTTCGCTAAAAGACAGGGTAAGGAAATGGACTCCTTAAGATTCTTGTACGACGGTA<br>TTAGAATTCAAGCTGATCAGACCCCTGAAGATTTGGACATGGAGGATAACGATATTATTGAGGC<br>TCACAGAGAACAGATTGGTGGTCTGCGTGCTTTCCAGTTCGCTCATCCGGACGACGACTTGCC<br>TGAaACCGGCGGACA |

| | |
|---|---|
| LaG21_BC7 | TTCCAATCTGACAACGCAATCGCTATGGCTCAGGTGCAGTTAGTCGAATCCGGGGGAGGCTTGGTTCAAGCAGGCGGTTCATTACGCCTGAGCTGTGCCGCATCCGGGCCAACAGGAGCTATGGCGTGGTTCCGCCAGGCACCGGGGATGGAGCGTGAATTCGTAGGCGGTATCTCGGGGTCAGAGACAGACACATACTACGCTGATTTCGTCAAAGGCCGTCTTACTGTAGACCGTGACAATGTTAAAAACACGGTTGACCTTCAAATGAACTCTTTGAAGCCAGAAGATACAGCGGTCTACTACTGTGCCGCGCGTCGCCGCGTCACGTTATTCACGTCACGCGCGGATTACGATTTTTGGGGGCAGGGAACCCAAGTTACCGTGAGTGGTGGCGGATCTATGTCGGACTCAGAAGTCAATCAAGAAGCTAAGCCAGAGGTCAAGCCAGAAGTCAAGCCTGAGACTCACATCAATTTAAAGGTGTCCGATGGATCTTCAGAGATCTTCTTCAAGATCAAAAAGACCACTCCTTTAAGAAGGCTGATGGAAGCGTTCGCTAAAAGACAGGGTAAGGAAATGGACTCCTTAAGATTCTTGTACGACGGTATTAGAATTCAAGCTGATCAGACCCCTGAAGATTTGGACATGGAGGATAACGATATTATTGAGGCTCACAGAGAACAGATTGGTGGTCGTCTGTTCGCTCAGCTGTTCCATACTGACGACGACTTGCCTGAaACCGGCGGACA |
| LaG24_BC5 | TTCCAATCTGACAACGCAATCGCTATGGCAGATGTGCAGTTGGTAGAGTCGGGCGGTGGACTTGTTCAGCCAGGTGGTTCATTACGTCTGTCTTGCGCTGCCTCGGGGGAGATTGCTTCAATTATTGCCATCGGCTGGTATCGCCAAGCCCCTGGTAAGCAGCGTGAGAGCGTGGCGTTAATCACCCGCAGTGGGATGATTACCTATGGCGATAGCGCCCAGGGTCGTTTCACGATTAGTCGCGATGACGCCAAGAACACTGTATACCTTCACATGGATGATCTTGTGCCAGAGGACACCGCCGTGTATTACTGCAACGCGAAGAAGGTATCTTTCGGAGACTATTGGGGACAGGGGACTCAAGTTACGGTCAGCGGTGGCGGATCTATGTCGGACTCAGAAGTCAATCAAGAAGCTAAGCCAGAGGTCAAGCCAGAAGTCAAGCCTGAGACTCACATCAATTTAAAGGTGTCCGATGGATCTTCAGAGATCTTCTTCAAGATCAAAAAGACCACTCCTTTAAGAAGGCTGATGGAAGCGTTCGCTAAAAGACAGGGTAAGGAAATGGACTCCTTAAGATTCTTGTACGACGGTATTAGAATTCAAGCTGATCAGACCCCTGAAGATTTGGACATGGAGGATAACGATATTATTGAGGCTCACAGAGAACAGATTGGTGGTCTGGCTCGTCAGTTCCTGTTCACTCATGACGACGACTTGCCTGAaACCGGCGGACA |
| LaG26_BC17 | TTCCAATCTGACAACGCAATCGCTATGGCTCAAGTCCAGCTTGTGGAATCGGGTGGGGGGTTAGTGCAGGCAGGTGCAAGTATGCGTTTATCGTGCGCGGCTAGCGGGATCACTTTCTCGTTGTATCATTGGGTATGGTTCCGCCAGGCTGCTGGCCGTGAACACGAATTTGTGGCCGGTATCATCCGCTCAGGGGGTGAGACGCTTAGCGCTGACTCAGTCAAAGACCGTTTCATCATTTCCCGCGACGATGCCAAAAACACCCTGTACCTTCAGATGAATATGCTTCAGCCCGAGGATACGGCCACTTATTACTGCGCAGCGACACACCGTGCCGATTGGTATTCCTCTGCGTTCCGTGAATACATTTTTCGCGGTCAGGGGACGCAGGTCACAGTATCGGGTGGCGGATCTATGTCGGACTCAGAAGTCAATCAAGAAGCTAAGCCAGAGGTCAAGCCAGAAGTCAAGCCTGAGACTCACATCAATTTAAAGGTGTCCGATGGATCTTCAGAGATCTTCTTCAAGATCAAAAAGACCACTCCTTTAAGAAGGCTGATGGAAGCGTTCGCTAAAAGACAGGGTAAGGAAATGGACTCCTTAAGATTCTTGTACGACGGTATTAGAATTCAAGCTGATCAGACCCCTGAAGATTTGGACATGGAGGATAACGATATTATTGAGGCTCACAGAGAACAGATTGGTGGTGCTCGTCTGTTCCAGTTCCAGCATCCGGACGACGACTTGCCTGAaACCGGCGGACA |
| LaG27_BC9 | TTCCAATCTGACAACGCAATCGCTATGGCTGATGTTCAGTTGGTCGAGTCTGGTGGCGGCCTGGTACAGGCAGGTGGATCCTTGCGTCTTTCATGTACCGCCAGTGGTTTAACCATTTCTACATACAACATCGGCTGGTTCCGTCAAGCTCCAGGTAAGGAGCGCGAATTCGTCGGTATCATTATTCGTAACGGAGATACCACTTATTATGCCGACTCCGTTAAAGGTCGTTTTACGATCAGTCGTGACAACGCGAAAAATACGGTGTATTTACAAATGAATTCTGTAAAGCCCGCCGACGCAGCCGTATATTCATGTGGTGCAACTGTACGCGCAGGCGCTGCGGCGGAACAATATAACTCTTACATCTTCCGTGGGCAAGGCACTCAGGTTACCGTAGGTGGCGGATCTATGTCGGACTCAGAAGTCAATCAAGAAGCTAAGCCAGAGGTCAAGCCAGAAGTCAAGCCTGAGACTCACATCAATTTAAAGGTGTCCGATGGATCTTCAGAGATCTTCTTCAAGATCAAAAAGACCACTCCTTTAAGAAGGCTGATGGAAGCGTTCGCTAAAAGACAGGGTAAGGAAATGGACTCCTTAAGATTCTTGTACGACGGTATTAGAATTCAAGCTGATCAGACCCCTGAAGATTTGGACATGGAGGATAACGATATTATTGAGGCTCACAGAGAACAGATTGGTGGTTTCCTGGCTCGTCAGTTCGCTCATCCGGACGACGACTTGCCTGAaACCGGCGGACA |
| LaG29_BC6 | TTCCAATCTGACAACGCAATCGCTATGGCTCAAGTACAGCTTGTAGAGTCAGGAGGGGGTTTGGTTCAGGCTGGCGCAGCTCTTCGCTTATCTTGTGCGGCATCCGGTGGGACCTTCAGTTTCTATAATATGGGGTGGTTTCGCCAAGCGCCAGGGAAGGAGCGCGAGTTTGTTGTTTCGATCTCTCGCAGTGGCGGGGGGACTGCTTATGCTGATTCCGTTAAAGGACGTTTTACAATCTCGCGCGATAACGCTAAGAATACCGCCTACTTACAGATGAATTCCCTGAAACCGGAGGATACGGCAGTTTATTATTGCGCGGCAGGTTTACGCGACTGGGGTCGTGAAGGTGAGCCCCATTACTGGGGCCAGGGCACTCAGGTTACGGTTAGTGGTGGCGGATCTATGTCGGACTCAGAAGTCAATCAAGAAGCTAAGCCAGAGGTCAAGCCAGAAGTCAAGCCTGAGACTCACATCAATTTAAAGGTGTCCGATGGATCTTCAGAGATCTTCTTCAAGATCAAAAAGACCACTCCTTTAAGAAGGCTGATGGAAGCGTTCGCTAAAAGACAGGGTAAGGAAATGGACTCCTTAAGATTCTTGTACGACGGTATTAGAATTCAAGCTGATCAGACCCCTGAAGATTTGGACATGGAGGATAACGATATTATTGAGGCTCACAGAGAACAGATTGGTGGTCGTCTGTTCCAGGCTCTGCAGCCGCATGACGACGACTTGCCTGAaACCGGCGGACA |

| | |
|---|---|
| LaG30_BC16 | TTCCAATCTGACAACGCAATCGCTATGGCCCAGGTACAGTTAGTAGAATCTGGAGGTGGGCTTGTCCAAGCGGGTGGCTCCTTACGCTTATCGTGCGCAGCATCTGGACGTACATTCTCCACGAGTGCGATGGGATGGTTCCGCCAAGCACCAGGGCGTGAACGTGAGTTTGTCGCTGCCATCACGTGGACGGTGGGTAACACTATCTATGGCGACAGCATGAAGGGTCGCTTCACTATCTCGCGTGATCGTACTAAAAATACTGTAGACTTACAGATGGATTCGCTTAAACCCGAGGATACCGCCGTCTATTACTGTACTGCGCGTTCACGCGGATTTGTACTGTCCGACCTTCGTTCCGTCGACTCTTTCGACTACAAGGGGCAGGGAACGCAGGTCACCGTATCCGGTGGCGGATCTATGTCGGACTCAGAAGTCAATCAAGAAGCTAAGCCAGAGGTCAAGCCAGAAGTCAAGCCTGAGACTCACATCAATTTAAAGGTGTCCGATGGATCTTCAGAGATCTTCTTCAAGATCAAAAAGACCACTCCTTTAAGAAGGCTGATGGAAGCGTTCGCTAAAAGACAGGGTAAGGAAATGGACTCCTTAAGATTCTTGTACGACGGTATTAGAATTCAAGCTGATCAGACCCCTGAAGATTTGGACATGGAGGATAACGATATTATTGAGGCTCACAGAGAACAGATTGGTGGTGCTCTGCAGTTCCGTCTGTTCCATACTGACGACGACTTGCCTGAaACCGGCGGACA |
| LaG37_BC3 | TTCCAATCTGACAACGCAATCGCTATGGCCCAAGTACAGTTTGTAGAATCGGGAGGAGGCACGGTTCAGGATGGAGACTTCTTACGCTTATCGTGTACGGCCTCAGGCGACACATTCAGCAATTATCATGCGGGCTGGTTTCGCCAACCGCCAGGACGCGAACGTGAGTTCGTAGCAGCCATTAGCTGGACAGGTGAAGGTACTCTTTACGCCGATAGTGTCAAAGGCCAATTCACCATTTCACGTGACAACGCAAAGAACGCGATGTACTTACAAATGAATCGTTTGAAGCCAGAGGACACTGCAGTTTACTACTGCGCTGCAGCACGTTCTGTAGGGTTTACATGGCGCAGTAGCAAGTCCAACGATTATGCCTATTGGGGGCAAGGGACGCAGGTGACCGTAGGTGGCGGATCTATGTCGGACTCAGAAGTCAATCAAGAAGCTAAGCCAGAGGTCAAGCCAGAAGTCAAGCCTGAGACTCACATCAATTTAAAGGTGTCCGATGGATCTTCAGAGATCTTCTTCAAGATCAAAAAGACCACTCCTTTAAGAAGGCTGATGGAAGCGTTCGCTAAAAGACAGGGTAAGGAAATGGACTCCTTAAGATTCTTGTACGACGGTATTAGAATTCAAGCTGATCAGACCCCTGAAGATTTGGACATGGAGGATAACGATATTATTGAGGCTCACAGAGAACAGATTGGTGGTCTGTTCGCTCGTCAGTTCCAGCATCCGGACGACGACTTGCCTGAaACCGGCGGACA |
| LaG41_BC13 | TTCCAATCTGACAACGCAATCGCTATGGCAGACGTTCAATTAGTAGAGTCAGGCGGCGGCCTTGTACAGGCTGGTGGGTCGTTGCGTCTGTCTTGCGCTGCTTCTGGCCCCACTGGAGCTATGGCGTGGTTCCGTCAGGCTCCAGGGAAAGAGCGCGAATTTGTGGGTGGGATTTCCGGTTCTGAGACGGATACATACTATGTAGACTCTGTAAAGGGTCGCTTTACTGTCGACCGCGACAATGTCAAGAACACGGTTTACTTGCAAATGAACTCTCTTAAACCGGAGGACACCGCTGTATACTACTGCGCGGCCCGCCGTCGCATCACTCTTTTTACATCCCGTACCGATTACGACTTTTGGGGGCGCGGTACTCAGGTCACCGTAGGTGGCGGATCTATGTCGGACTCAGAAGTCAATCAAGAAGCTAAGCCAGAGGTCAAGCCAGAAGTCAAGCCTGAGACTCACATCAATTTAAAGGTGTCCGATGGATCTTCAGAGATCTTCTTCAAGATCAAAAAGACCACTCCTTTAAGAAGGCTGATGGAAGCGTTCGCTAAAAGACAGGGTAAGGAAATGGACTCCTTAAGATTCTTGTACGACGGTATTAGAATTCAAGCTGATCAGACCCCTGAAGATTTGGACATGGAGGATAACGATATTATTGAGGCTCACAGAGAACAGATTGGTGGTCAGTTCGCTCGTCTGTTCGCTCATCCGGACGACGACTTGCCTGAaACCGGCGGACA |
| LaG42_BC18 | TTCCAATCTGACAACGCAATCGCTATGGCAGACGTTCAGCTTGTCGAATCCGGGGGCGGGCTTGTCCAAGCTGGGGACTCCCTTCGCCTGTCATGCGCCGCTAGTGGTCCGACAGGAGCGATGGCATGGTTTCATCAAGGTCTTGGGAAGGAACGTGAATTCGTGGGCGGAATCTCACCGAGTGGAGATAACATTTACTATGCCGATTCGGTTAAAGGCCGTTTTACGATTGACCGCGACAACGCGAAAAATACAGTTAGCTTGCAAATGAATTCATTGAAGCCAGAAGATATGGGGGTATACTATTGTGCAGCCCGCCGTCGCGTCACACTTTTTACCTCACGTACTGATTACGAATTCTGGGGTCGTGGCACTCAGGTTACTGTTTCCGGTGGCGGATCTATGTCGGACTCAGAAGTCAATCAAGAAGCTAAGCCAGAGGTCAAGCCAGAAGTCAAGCCTGAGACTCACATCAATTTAAAGGTGTCCGATGGATCTTCAGAGATCTTCTTCAAGATCAAAAAGACCACTCCTTTAAGAAGGCTGATGGAAGCGTTCGCTAAAAGACAGGGTAAGGAAATGGACTCCTTAAGATTCTTGTACGACGGTATTAGAATTCAAGCTGATCAGACCCCTGAAGATTTGGACATGGAGGATAACGATATTATTGAGGCTCACAGAGAACAGATTGGTGGTGCTCGTCAGTTCCTGGCTCAGCCGCATGACGACGACTTGCCTGAaACCGGCGGACA |
| LaG43_BC15 | TTCCAATCTGACAACGCAATCGCTATGGCGGATGTCCAACTGGTAGAGTCGGGAGGCGGCCTGGTACAACCAGGTGGATCGTTACGTCTGAGCTGCGAAGCAAGCGGAGGTGCCTTTTCAACAGTGGCTATGGGTTGGTTCCGTCAAGCACCAGGAAAAGAGCGCGAATTCGTAGGGGCGATCACGTGGACGGCGGGGAGCACCTATTATGCAGACAGTGCAAAAGGCCGTTTCACCATTTCCCGTGATAACGCAAAGAACACGGTTCACCTGCAAATGAATAGTTTGAAGCCAGAGGACACCGCTGTCTACTATTGTGCTCAACGTGTGCGCGGGTTCTTCGGCCCGCTTCGCACTACTCCCTCTTGGTATGAATACTGGGGACAAGGAACCCAGGTCACTGTGAGCGGTGGCGGATCTATGTCGGACTCAGAAGTCAATCAAGAAGCTAAGCCAGAGGTCAAGCCAGAAGTCAAGCCTGAGACTCACATCAATTTAAAGGTGTCCGATGGATCTTCAGAGATCTTCTTCAAGATCAAAAAGACCACTCCTTTAAGAAGGCTGATGGAAGCGTTCGCTAAAAGACAGGGTAAGGAAATGGACTCCTTAAGATTCTTGTACGACGGTATTAGAATTCAAGCTGATCAGACCCCTGAAGATTTGGACATGGAGGATAACGATATTATTGAGG |

| | |
|---|---|
| | CTCACAGAGAACAGATTGGTGGTCAGGCTTTCCTGCGTCTGTTCCATACTGACGACGACTTGC<br>CTGAaACCGGCGGACA |
| LaM1_BC12 | TTCCAATCTGACAACGCAATCGCTATGGCTCAGGTTCAATTGGTCGAGTCAGGGGGTGGTCTG<br>GTCCAAGCCGGAGATTCTCTTCGTTTATCATGTGCAGCAAGCGGACGCACTTTTGAGAACTAT<br>GCTATGGGGTGGTTTCGTCAGGCCCCAGGTAAGGAACGCGAGTTTGTCGGTGCAGTTTCCTG<br>GGGCGGAGGTCGTACCTATTACGCTGACAATGTTAAAGGCCGTTTCACGATTTCCCGTGATAA<br>TGCAAAGAAAAACACCGTTTATTTGCAGATGAACTCGTTAAAGCCTGAAGATACCGCCGTCTAT<br>TACTGTGCCGCTAAATCCGTCCTGACAATCGCAACTATGCGCGTCCCGGACGAATATAACTAT<br>TGGGGCCAAGGCACTCAGGTGACCGTGTCCGGTGGCGGATCTATGTCGGACTCAGAAGTCAA<br>TCAAGAAGCTAAGCCAGAGGTCAAGCCAGAAGTCAAGCCTGAGACTCACATCAATTTAAAGGT<br>GTCCGATGGATCTTCAGAGATCTTCTTCAAGATCAAAAAGACCACTCCTTTAAGAAGGCTGATG<br>GAAGCGTTCGCTAAAAGACAGGGTAAGGAAATGGACTCCTTAAGATTCTTGTACGACGGTATT<br>AGAATTCAAGCTGATCAGACCCCTGAAGATTTGGACATGGAGGATAACGATATTATTGAGGCT<br>CACAGAGAACAGATTGGTGGTCAGCGTGCTTTCCTGTTCCTGCATACTGACGACGACTTGCCT<br>GAaACCGGCGGACA |

### Supplementary Table 4. Gene sequences encoding GFP ligands

| Name | Sequence |
|---|---|
| GFP | ATGTCTAAAGGGGAAGAACTTTTCACAGGGGTGGTCCCGATCTTGGTGGAACTTGACGGAGATGTAAATGGGCACAAATTTTCTGTTTCCGGTGAAGGAGAGGGTGATGCCACTTATGGTAAATTAACGTTAAAATTTATCTGCACAACCGGAAAACTGCCTGTGCCCTGGCCTACACTGGTCACCACGTTTAGTTACGGGGTTCAGTGCTTCAGTCGTTACCCAGACCACATGAAACAACATGATTTCTTTAAATCCGCGATGCCAGAGGGCTATGTCCAGGAGCGTACAATTTTCTTCAAAGATGACGGAAACTATAAGACTCGTGCCGAAGTAAAATTTGAGGGTGATACCCTTGTCAACCGTATCGAGCTGAAGGGCATTGACTTCAAGGAAGATGGTAACATCTTAGGTCACAAACTGGAGTACAACTATAACTCCCATAATGTGTACATCATGGCGGACAAGCAAAAGAACGGAATCAAGGTAAACTTCAAAATTCGTCATAATATTGAGGATGGAAGTGTACAGTTAGCCGACCACTACCAGCAAAACACCCCCATTGGGGATGGCCCTGTGCTGCTGCCTGACAACCACTATCTGTCGACACAGAGCGCATTGTCTAAGGACCCAAACGAGAAGCGTGATCACATGGTATTGCTGGAATTCGTAACGGCTGCTGGCATTACGCATGGAATGGATGAGTTATACAAG |
| RLFA_GS_GFP_2xFLAG | gctaGAAGACctTGGTCGCTTATTCGCCGGCGGGGGGTCCATGAGTAAGGGTGAAGAATTATTCACGGGGGTGGTGCCAATCTTGGTCGAACTGGACGGCGATGTAAACGGACACAAGTTTTCCGTGTCAGGTGAAGGTGAAGGCGATGCGACTTATGGCAAACTTACGCTTAAATTCATTTGCACAACCGGCAAACTGCCCGTTCCGTGGCCGACTTTGGTTACTACATTTAGTTACGGAGTTCAATGCTTCTCACGCTACCCCGACCATATGAAACAGCACGATTTTTTTAAATCGGCGATGCCCGAAGGCTATGTGCAAGAGCGTACAATTTTTTTTAAAGACGATGGAAATTACAAAACCCGCGCTGAGGTCAAGTTCGAGGGCGATACTCTTGTTAATCGCATCGAGCTGAAAGGTATTGATTTTAAGGAAGATGGCAATATTTTGGGACACAAGTTAGAATATAACTATAACTCACATAATGTCTATATTATGGCGGACAAACAGAAGAACGGCATTAAGGTGAACTTTAAAATTCGTCACAATATCGAGGACGGCTCCGTCCAATTAGCCGACCATTACCAACAAAATACGCCAATTGGTGACGGGCCGGTGTTGTTACCAGACAACCATTACCTTTCTACACAAAGCGCATTATCAAAAGATCCTAATGAAAAGCGCGATCACATGGTTTTGCTTGAATTTGTCACAGCCGCAGGCATCACGCACGGCATGGATGAGTTATATAAGGGTGGCGGCTCGGGCGGTTCAGATTACAAAGACGATGACGATAAGGGTGGCGGATCAGATTATAAGGACGATGACGACAAATGACTCGAGTAAGGTTA |
| FAQR_GS_GFP_2xFLAG | gctaGAAGACctTGGTTTTGCTCAACGCGGCGGTGGAAGTATGTCCAAGGGTGAAGAGCTTTTCACCGGAGTAGTGCCTATTCTTGTCGAACTTGATGGTGATGTGAACGGTCACAAGTTCTCTGTCTCGGGCGAAGGCGAAGGTGATGCGACATACGGGAAGTTGACCCTGAAGTTCATTTGCACAACCGGCAAATTACCGGTTCCATGGCCTACGTTGGTAACAACATTCTCTTACGGCGTACAATGTTTTTCACGCTATCCAGACCACATGAAGCAGCACGACTTTTTCAAATCTGCAATGCCGGAGGGTTACGTCCAGGAGCGTACAATTTTCTTCAAAGATGATGGGAATTACAAAACCCGTGCAGAGGTCAAGTTCGAGGGCGACACCTTGGTAAATCGCATCGAATTGAAGGGGATCGATTTCAAGGAAGATGGGAATATCTTGGGCCACAAATTGGAGTATAACTACAATAGCCACAATGTCTATATCATGGCAGACAAACAGAAGAACGGGATCAAAGTAAACTTCAAAATTCGTCACAATATCGAGGATGGTTCAGTTCAGTTAGCAGATCACTATCAGCAAAATACGCCAATCGGGGACGGACCAGTGCTTTTGCCGGACAATCACTATTTATCAACACAAAGCGCCTTAAGTAAAGACCCCAATGAAAAACGCGACCATATGGTCTTACTTGAATTCGTGACGGCTGCAGGTATCACCCACGGTATGGATGAGCTGTATAAGGGAGGCGGCAGCGGAGGTTCTGACTATAAAGACGACGATGACAAAGGCGGTGGCAGTGACTATAAAGACGACGACGACAAATGACTCGAGTAAGGTTA |
| LARQ_GS_GFP_2xFLAG | gctaGAAGACctTGGTCTTGCCCGTCAAGGAGGAGGTAGCATGAGTAAGGGTGAAGAGTTATTTACCGGTGTTGTCCCAATTCTTGTGGAGTTAGACGGTGATGTTAATGGCCATAAATTTAGTGTCTCTGGAGAAGGTGAAGGAGATGCTACTTATGGTAAGTTGACGCTGAAGTTTATTTGTACTACGGGAAAACTTCCGGTCCCGTGGCCTACCCTGGTTACGACGTTTAGTTATGGCGTCCAATGTTTTTCGCGCTATCCAGACCATATGAAACAACATGACTTTTTTAAGAGTGCCATGCCTGAGGGTTATGTCCAAGAACGCACCATTTTTTTCAAGGATGACGGGAACTACAAGACACGTGCTGAAGTAAAATTTGAGGGAGACACGTTAGTGAATCGCATTGAATTAAAAGGAATTGATTTTAAGGAAGATGGCAATATCCTTGGACACAAATTGGAGTACAATTACAATTCCCATAATGTCTACATCATGGCGGATAAGCAGAAGAATGGCATCAAGGTCAACTTTAAGATTCGCCATAACATCGAAGATGGCAGTGTACAGTTAGCAGACCATTACCAGCAAAATACACCTATTGGGGACGGTCCGGTGCTTTTGCCCGACAACCACTACTTATCTACTCAATCCGCGTTAAGTAAGGATCCAAATGAAAAGCGCGATCATATGGTACTGTTAGAGTTTGTTACAGCAGCCGGTATTACTCACGGCATGGACGAATTATACAAGGGAGGTGGTTCCGGAGGATCAGATTACAAAGACGATGACGACAAAGGTGGCGGTTCAGATTACAAAGACGACGATGATAAATGACTCGAGTAAGGTTA |
| RLFA_PA_GFP_2xFLAG | gctaGAAGACctTGGTCGTTTATTCGCTCCAGCGCCCGCTCCCGCGATGTCAAAGGGTGAGGAATTGTTCACCGGCGTAGTTCCTATTCTTGTGGAGCTGGATGGTGACGTGAATGGACATAAATTCTCTGTATCTGGTGAAGGAGAAGGTGATGCAACGTACGGGAAGCTGACGCTGAAGTTTATCTGTACAACTGGCAAGCTTCCAGTTCCGTGGCCTACACTTGTTACAACTTTTTCTTACGGGGTGCAGTGTTTCTCACGTTATCCTGACCACATGAAGCAGCATGATTTCTTTAAGAGTGCCATGCCCGAAGGATATGTCCAAGAACGTACTATCTTTTTTAAGGACGACGGCAATTACAAAACGCGCGCTGAGGTAAAGTTCGAGGGTGATACCCTTGTGAACCGTATTGAGCTGAAAGGGATCGACTTCAAGGAGGATGGTAACATTTTGGGACATAAGTTGG |

| | |
|---|---|
| | AGTATAATTACAATAGTCACAACGTATACATTATGGCGGATAAACAAAAGAACGGAATCAAAGTTAATTTCAAGATCCGCCATAACATCGAGGACGGCTCAGTTCAACTGGCCGATCACTACCAACAGAATACTCCAATTGGTGATGGGCCCGTTTTGCTTCCTGATAACCACTACCTGTCAACGCAAAGCGCCTTATCCAAGGACCCTAACGAAAAGCGCGATCATATGGTCTTATTAGAATTCGTAACGGCGGCGGGCATTACACACGGCATGGACGAACTGTATAAAGGGGGAGGCTCTGGTGGAAGCGATTACAAGGACGACGATGATAAGGGAGGCGGGAGTGACTATAAAGATGATGACGACAAATGACTCGAGTAAGGTTA |
| RLFA_GS_GFP | gctaGAAGACctTGGTCGTCTTTTTGCTGGTGGAGGCTCAATGTCTAAAGGGGAAGAACTTTTCACAGGGGTGGTCCCGATCTTGGTGGAACTTGACGGAGATGTAAATGGGCACAAATTTTCTGTTTCCGGTGAAGGAGAGGGTGATGCCACTTATGGTAAATTAACGTTAAAATTTATCTGCACAACCGGAAAACTGCCTGTGCCCTGGCCTACACTGGTCACCACGTTTAGTTACGGGGTTCAGTGCTTCAGTCGTTACCCAGACCACATGAAACAACATGATTTCTTTAAATCCGCGATGCCAGAGGGCTATGTCCAGGAGCGTACAATTTTCTTCAAAGATGACGGAAACTATAAGACTCGTGCCGAAGTAAAATTTGAGGGTGATACCCTTGTCAACCGTATCGAGCTGAAGGGCATTGACTTCAAGGAAGATGGTAACATCTTAGGTCACAAACTGGAGTACAACTATAACTCCCATAATGTGTACATCATGGCGGACAAGCAAAAGAACGGAATCAAGGTAAACTTCAAAATTCGTCATAATATTGAGGATGGAAGTGTACAGTTAGCCGACCACTACCAGCAAAACACCCCCATTGGGGATGGCCCTGTGCTGCTGCCTGACAACCACTATCTGTCGACACAGAGCGCATTGTCTAAGGACCCAAACGAGAAGCGTGATCACATGGTATTGCTGGAATTCGTAACGGCTGCTGGCATTACGCATGGAATGGATGAGTTATACAAGTTGCagGTCTTCgcta |
| FAQR_GS_GFP | gctaGAAGACctTGGTTTTGCGCAGCGTGGTGGAGGCTCAATGTCTAAAGGGGAAGAACTTTTCACAGGGGTGGTCCCGATCTTGGTGGAACTTGACGGAGATGTAAATGGGCACAAATTTTCTGTTTCCGGTGAAGGAGAGGGTGATGCCACTTATGGTAAATTAACGTTAAAATTTATCTGCACAACCGGAAAACTGCCTGTGCCCTGGCCTACACTGGTCACCACGTTTAGTTACGGGGTTCAGTGCTTCAGTCGTTACCCAGACCACATGAAACAACATGATTTCTTTAAATCCGCGATGCCAGAGGGCTATGTCCAGGAGCGTACAATTTTCTTCAAAGATGACGGAAACTATAAGACTCGTGCCGAAGTAAAATTTGAGGGTGATACCCTTGTCAACCGTATCGAGCTGAAGGGCATTGACTTCAAGGAAGATGGTAACATCTTAGGTCACAAACTGGAGTACAACTATAACTCCCATAATGTGTACATCATGGCGGACAAGCAAAAGAACGGAATCAAGGTAAACTTCAAAATTCGTCATAATATTGAGGATGGAAGTGTACAGTTAGCCGACCACTACCAGCAAAACACCCCCATTGGGGATGGCCCTGTGCTGCTGCCTGACAACCACTATCTGTCGACACAGAGCGCATTGTCTAAGGACCCAAACGAGAAGCGTGATCACATGGTATTGCTGGAATTCGTAACGGCTGCTGGCATTACGCATGGAATGGATGAGTTATACAAGTTGCagGTCTTCgcta |
| LARQ_GS_GFP | gctaGAAGACctTGGTTTGGCGCGTCAGGGTGGAGGCTCAATGTCTAAAGGGGAAGAACTTTTCACAGGGGTGGTCCCGATCTTGGTGGAACTTGACGGAGATGTAAATGGGCACAAATTTTCTGTTTCCGGTGAAGGAGAGGGTGATGCCACTTATGGTAAATTAACGTTAAAATTTATCTGCACAACCGGAAAACTGCCTGTGCCCTGGCCTACACTGGTCACCACGTTTAGTTACGGGGTTCAGTGCTTCAGTCGTTACCCAGACCACATGAAACAACATGATTTCTTTAAATCCGCGATGCCAGAGGGCTATGTCCAGGAGCGTACAATTTTCTTCAAAGATGACGGAAACTATAAGACTCGTGCCGAAGTAAAATTTGAGGGTGATACCCTTGTCAACCGTATCGAGCTGAAGGGCATTGACTTCAAGGAAGATGGTAACATCTTAGGTCACAAACTGGAGTACAACTATAACTCCCATAATGTGTACATCATGGCGGACAAGCAAAAGAACGGAATCAAGGTAAACTTCAAAATTCGTCATAATATTGAGGATGGAAGTGTACAGTTAGCCGACCACTACCAGCAAAACACCCCCATTGGGGATGGCCCTGTGCTGCTGCCTGACAACCACTATCTGTCGACACAGAGCGCATTGTCTAAGGACCCAAACGAGAAGCGTGATCACATGGTATTGCTGGAATTCGTAACGGCTGCTGGCATTACGCATGGAATGGATGAGTTATACAAGTTGCagGTCTTCgcta |

### Plasmid sequences

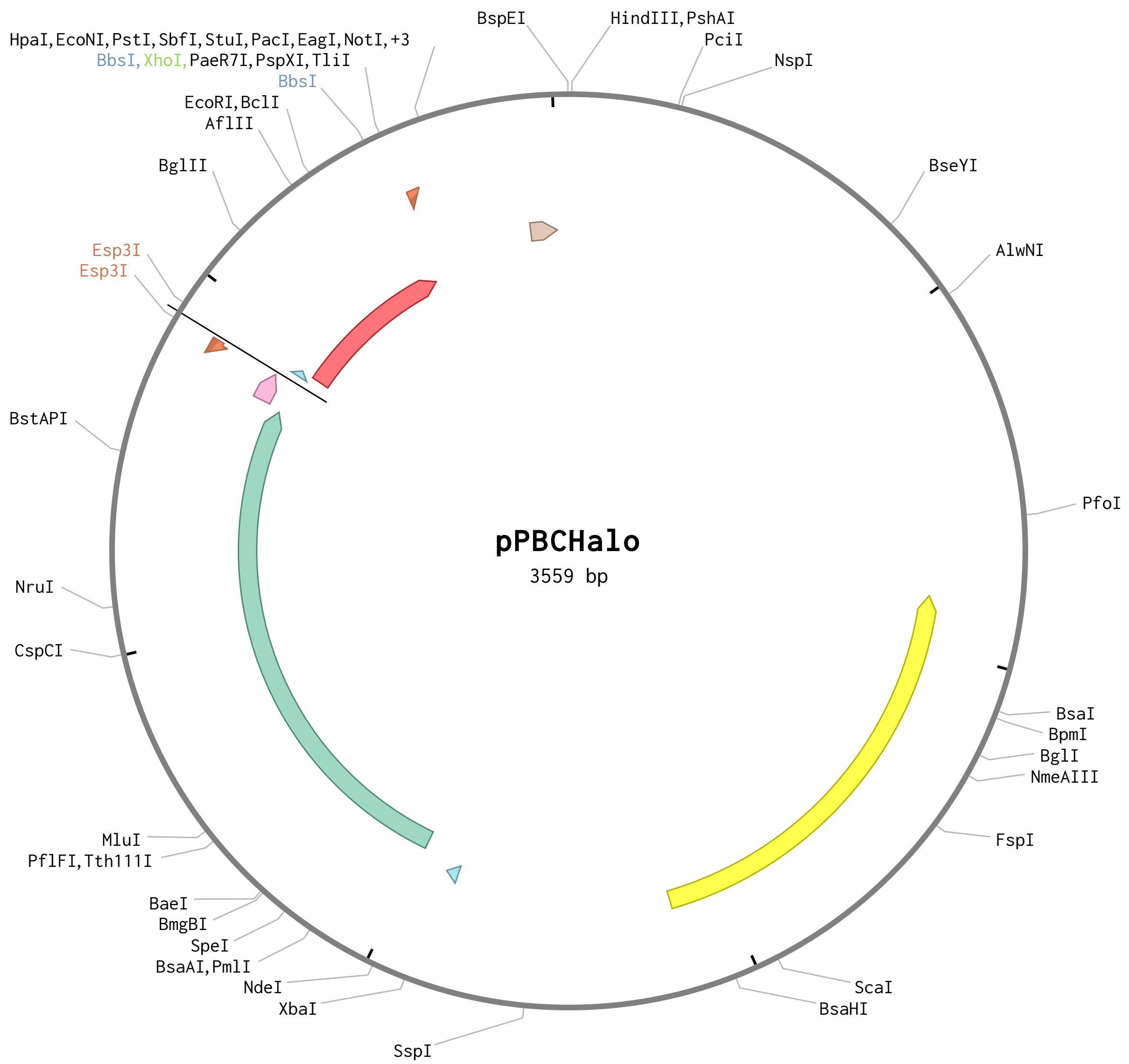

```
LOCUS       pPBCHalo                3559 bp ds-DNA     circular     06-MAR-2025
DEFINITION  .
COMMENT     Imported using the Genbank importer. File name:
            pNEB_Halo_TEV_SUMO_Sortase.ape ApEinfo:methylated:1
```

```
FEATURES             Location/Qualifiers

     CDS             complement(921..1580)

                     /label="AmpR"

                     /ApEinfo_revcolor="#ffff00"

                     /ApEinfo_fwdcolor="#ffff00"

                     /locus_tag="AmpR"

                     /label="AmpR"

                     /ApEinfo_graphicformat="arrow_data {{0 0.5 0 1 2 0 0 -1 0-0.5} {}
0} width 5 offset 0"

/translation="MSTFKVLLCGAVLSRIDAGQEQLGRRIHYSQNDLVEYSPVTEKHLTDGMTVRELCSAAITMSDNTAANLLLTT
IGGPKELTAFLHNMGDHVTRLDRWEPELNEAIPNDERDTTMPVAMATTLRKLLTGELLTLASRQQLIDWMEADKVAGPLLRSALPAG
WFIADKSGAGERGSRGIIAALGPDGKPSRIVVIYTTGSQATMDERNRQIAEIGASLIKHW"

     primer_bind     1927..1946

                     /label="T7"

                     /ApEinfo_revcolor="#85dae9"

                     /ApEinfo_fwdcolor="#85dae9"

                     /locus_tag="T7"

                     /label="T7"

                     /ApEinfo_graphicformat="arrow_data {{0 0.5 0 1 2 0 0 -1 0-0.5} {}
0} width 5 offset 0"

     misc_feature    1995..2882

                     /label="Halo"

                     /ApEinfo_revcolor="#75c6a9"

                     /ApEinfo_fwdcolor="#75c6a9"

     misc_feature    2889..2936

                     /label="Promega TEV optimized site"

                     /ApEinfo_revcolor="#ff9ccd"

                     /ApEinfo_fwdcolor="#ff9ccd"

     primer          complement(2913..2936)

                     /label="GhM50_pHalo.SUMO_Rev"

                     /note="sequence: AGCGATTGCGTTGTCAGATTGGAA"

                     /ApEinfo_revcolor="#f58a5e"

                     /ApEinfo_fwdcolor="#f58a5e"

     misc_feature    2955..2966

                     /label="GS linker"
```

```
                     /ApEinfo_revcolor="#85dae9"
                     /ApEinfo_fwdcolor="#85dae9"
                     /locus_tag="GS linker"
                     /label="GS linker"
                     /ApEinfo_graphicformat="arrow_data {{0 0.5 0 1 2 0 0 -1 0-0.5} {}
0} width 5 offset 0"
     misc_feature    2967..3260
                     /label="scSUMO"
                     /ApEinfo_revcolor="#ff3843"
                     /ApEinfo_fwdcolor="#ff3843"
                     /locus_tag="scSUMO"
                     /label="scSUMO"
                     /ApEinfo_graphicformat="arrow_data {{0 0.5 0 1 2 0 0 -1 0-0.5} {}
0} width 5 offset 0"
     primer          3279..3298
                     /label="GhM49_pHalo-SUMO_Fwd"
                     /note="sequence: TTGCCTGAaACCGGCGGACA"
                     /ApEinfo_revcolor="#f58a5e"
                     /ApEinfo_fwdcolor="#f58a5e"
     attenuator      3453..3499
                     /label="T7 terminator"
                     /ApEinfo_revcolor="#d6b295"
                     /ApEinfo_fwdcolor="#d6b295"
                     /locus_tag="T7 terminator"
                     /label="T7 terminator"
                     /ApEinfo_graphicformat="arrow_data {{0 0.5 0 1 2 0 0 -1 0-0.5} {}
0} width 5 offset 0"
ORIGIN
        1 CGCTGCGCTC GGTCGTTCGG CTGCGGCGAG CGGTATCAGC TCACTCAAAG GCGGTAATAC
       61 GGTTATCCAC AGAATCAGGG GATAACGCAG GAAAGAACAT GTGAGCAAAA GGCCAGCAAA
      121 AGGCCAGGAA CCGTAAAAAG GCCGCGTTGC TGGCGTTTTT CCATAGGCTC CGCCCCCCTG
      181 ACGAGCATCA CAAAAATCGA CGCTCAAGTC AGAGGTGGCG AAACCCGACA GGACTATAAA
      241 GATACCAGGC GTTTCCCCCT GGAAGCTCCC TCGTGCGCTC TCCTGTTCCG ACCCTGCCGC
      301 TTACCGGATA CCTGTCCGCC TTTCTCCCTT CGGGAAGCGT GGCGCTTTCT CATAGCTCAC
      361 GCTGTAGGTA TCTCAGTTCG GTGTAGGTCG TTCGCTCCAA GCTGGGCTGT GTGCACGAAC
```

421 CCCCCGTTCA GCCCGACCGC TGCGCCTTAT CCGGTAACTA TCGTCTTGAG TCCAACCCGC
481 TAAGACACGA CTTATCGCCA CTGGCAGCAG CCACTGGTAA CAGGATTAGC AGAGCGAGGT
541 ATGTAGGCGG TGCTACAGAG TTCTTGAAGT GGTGGCCTAA CTACGGCTAC ACTAGAAGAA
601 CAGTATTTGG TATCTGCGCT CTGCTGAAGC CAGTTACCTT CGGAAAAAGA GTTGGTAGCT
661 CTTGATCCGG CAAACAAACC ACCGCTGGTA GCGGTGGTTT TTTTGTTTGC AAGCAGCAGA
721 TTACGCGCAG AAAAAAAGGA TCTCAAGAAG ATCCTTTGAT CTTTTCTACG GGGTCTGACG
781 CTCAGTGGAA CGAAAACTCA CAGATCCGGG ATTTTGGTCA TGAGATTATC AAAAAGGATC
841 TTCACCTAGA TCCTTTTAAA TTAAAAATGA AGTTTTAAAT CAATCTAAAG TATATATGAG
901 TAAACTTGGT CTGACAGTTA CCAATGCTTA ATCAGTGAGG CACCTATCTC AGCGATCTGT
961 CTATTTCGTT CATCCATAGT TGCCTGACTC CCCGTCGTGT AGATAACTAC GATACGGGAG
1021 GGCTTACCAT CTGGCCCCAG TGCTGCAATG ATACCGCGAG ACCCACGCTC ACCGGCTCCA
1081 GATTTATCAG CAATAAACCA GCCAGCCGGA AGGGCCGAGC GCAGAAGTGG TCCTGCAACT
1141 TTATCCGCCT CCATCCAGTC TATTAATTGT TGCCGGGAAG CTAGAGTAAG TAGTTCGCCA
1201 GTTAATAGTT TGCGCAACGT TGTTGCCATT GCTACAGGCA TCGTGGTGTC ACGCTCGTCG
1261 TTTGGTATGG CTTCATTCAG CTCCGGTTCC CAACGATCAA GGCGAGTTAC ATGATCCCCC
1321 ATGTTGTGCA AAAAAGCGGT TAGCTCCTTC GGTCCTCCGA TCGTTGTCAG AAGTAAGTTG
1381 GCCGCAGTGT TATCACTCAT GGTTATGGCA GCACTGCATA ATTCTCTTAC TGTCATGCCA
1441 TCCGTAAGAT GCTTTTCTGT GACTGGTGAG TACTCAACCA AGTCATTCTG AGAATAGTGT
1501 ATGCGGCGAC CGAGTTGCTC TTGCCCGGCG TCAATACGGG ATAATACCGC GCCACATAGC
1561 AGAACTTTAA AAGTGCTCAT CATTGGAAAA CGTTCTTCGG GGCGAAAACT CTCAAGGATC
1621 TTACCGCTGT TGAGATCCAG TTCGATGTAA CCCACTCGTG CACCCAACTG ATCTTCAGCA
1681 TCTTTTACTT TCACCAGCGT TTCTGGGTGA GCAAAAACAG GAAGGCAAAA TGCCGCAAAA
1741 AAGGGAATAA GGGCGACACG GAAATGTTGA ATACTCATAC TCTTCCTTTT TCAATATTAT
1801 TGAAGCATTT ATCAGGGTTA TTGTCTCATG AGCGGATACA TATTTGAATG TATTTAGAAA
1861 AATAAACAAA TAGGGGTTCC GCGCACATTT CCCCGAAAAG TGCTAGTGGT GCtagccccg
1921 cgaaattaat acgACTCACT ATAGGGTCTA GAAGAAATAA TTTTGTTTAA CTTTAAGAAG
1981 GAGATATACA TATGgcggag attggcacag gatttccatt tgatccccac tatgttgagg
2041 ttcttggtga acgtatgcac tatgttgacg ttggaccacg tgacggaacc cctgttttat
2101 ttttgcatgg taatccgact agtagctacg tttggcgcaa catcatccca cacgtcgccc
2161 ctactcaccg ttgtatcgca ccggacttga tcggtatggg gaagtccgac aagccagact
2221 taggctattt ctttgacgac catgtccgtt tcatggacgc gtttatcgag gcccttgggt
2281 tagaagaggt tgtacttgtc attcacgact gggggtccgc tcttggcttt cattgggcta
2341 aacgtaaccc cgagcgcgtc aaggggatcg cctttatgga gtttatccgc cccatcccta
2401 cctgggatga gtggccggaa tttgcgcgtg aaacgtttca agccttccgc acaactgatg

2461 ttggccgcaa attaattatc gatcagaacg tgttcatcga ggggacattg ccaatgggag

2521 tggttcgtcc gttgaccgaa gttgaaatgg accattatcg cgaaccgttt ctgaacccgg

2581 tcgatcgtga acccttatgg cgctttccta acgaattgcc gattgctggt gaacctgcga

2641 acattgtcgc attagtagag gaatacatgg actggttaca tcaatccccc gttcctaaat

2701 tgttgttttg gggaacgcca ggcgtattaa tcccgccagc ggaggccgca cgccttgcga

2761 agtcacttcc taattgcaaa gcagtggata ttggcccagg cttgaatctt ttgcaagagg

2821 ataaccctga tctgatcggg tcggaaattg cgcgttggtt atcgaccctt gagatttcgg

2881 gcACCAGCGA ACCAACAACT GAGGACTTGT ATTTCCAATC TGACAACGCA ATCGCTaGAG

2941 ACGgctaCGT CTCgGGTGGC GGATCTATGT CGGACTCAGA AGTCAATCAA GAAGCTAAGC

3001 CAGAGGTCAA GCCAGAAGTC AAGCCTGAGA CTCACATCAA TTTAAAGGTG TCCGATGGAT

3061 CTTCAGAGAT CTTCTTCAAG ATCAAAAAGA CCACTCCTTT AAGAAGGCTG ATGGAAGCGT

3121 TCGCTAAAAG ACAGGGTAAG GAAATGGACT CCTTAAGATT CTTGTACGAC GGTATTAGAA

3181 TTCAAGCTGA TCAGACCCCT GAAGATTTGG ACATGGAGGA TAACGATATT ATTGAGGCTC

3241 ACAGAGAACA GATTGGTGGT agGTCTTCga GAAGACctTT GCCTGAaACC GGCGGACATC

3301 ACCACCATCA TCACTGACTC GAGTAAGGTT AACCTGCAGG AGGCCTTTAA TTAAGGTGGT

3361 GCGGCCGCGC TAGCGGTCCC GGGGGATCGA TCCGGCTGCT AACAAAGCCC GAAAGGAAGC

3421 TGAGTTGGCT GCTGCCACCG CTGAGCAATA ACTAGCATAA CCCCTTGGGG CCTCTAAACG

3481 GGTCTTGAGG GGTTTTTTGC TGAAAGGAGG AACTATATCC GGAAGCTTGG CACTGGCCGA

3541 CCGGGGTCGA GCACTGACT

//

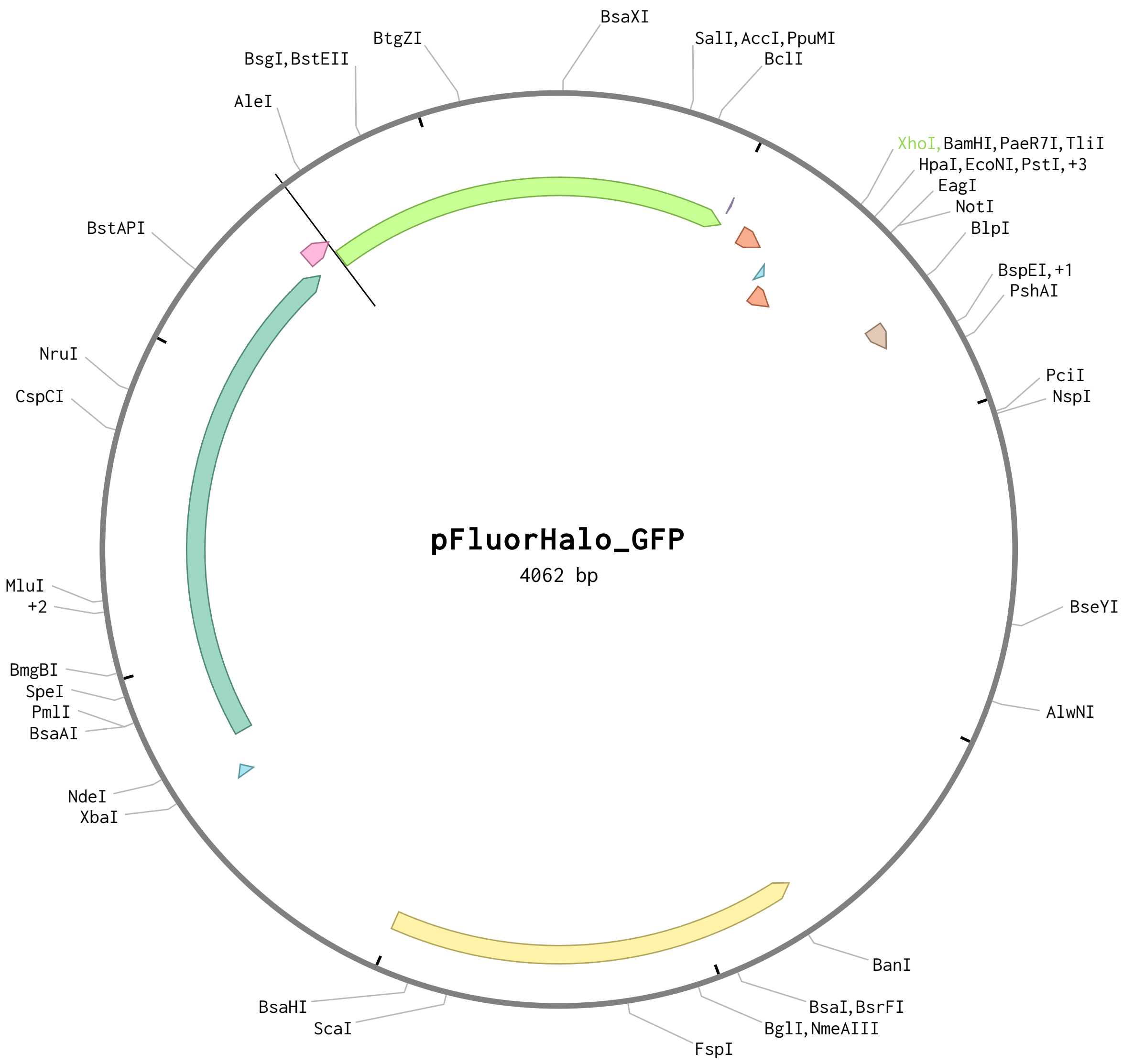

```
LOCUS       pFluorHalo_GFP          4062 bp ds-DNA     circular     06-MAR-2025
DEFINITION  .
COMMENT     Imported using the Genbank importer. File name:
            pNEB-halotag-tev-GFP-avitag.ape ApEinfo:methylated:1
FEATURES             Location/Qualifiers
```

primer_bind     286..302

/label="T7"

/ApEinfo_revcolor="#85dae9"

/ApEinfo_fwdcolor="#85dae9"

/locus_tag="T7"

/label="T7"

/ApEinfo_graphicformat="arrow_data {{0 0.5 0 1 2 0 0 -1 0-0.5} {} 0} width 5 offset 0"

CDS             348..2165

/ApEinfo_revcolor="#84b0dc"

/ApEinfo_fwdcolor="#84b0dc"

/translation="MAEIGTGFPFDPHYVEVLGERMHYVDVGPRDGTPVLFLHGNPTSSYVWRNIIPHVAPTHRCIAPDLIGMGKSD
KPDLGYFFDDHVRFMDAFIEALGLEEVVLVIHDWGSALGFHWAKRNPERVKGIAFMEFIRPIPTWDEWPEFARETFQAFRTTDVGRK
LIIDQNVFIEGTLPMGVVRPLTEVEMDHYREPFLNPVDREPLWRFPNELPIAGEPANIVALVEEYMDWLHQSPVPKLLFWGTPGVLI
PPAEAARLAKSLPNCKAVDIGPGLNLLQEDNPDLIGSEIARWLSTLEISGEPTTEDLYFQSDNAIAMSKGEELFTGVVPILVELDGD
VNGHKFSVSGEGEGDATYGKLTLKFICTTGKLPVPWPTLVTTFSYGVQCFSRYPDHMKQHDFFKSAMPEGYVQERTIFFKDDGNYKT
RAEVKFEGDTLVNRIELKGIDFKEDGNILGHKLEYNYNSHNVYIMADKQKNGIKVNFKIRHNIEDGSVQLADHYQQNTPIGDGPVLL
PDNHYLSTQSALSKDPNEKRDHMVLLEFVTAAGITHGMDELYKGGGSGGGSGGGSGLNDFFEAQKIEWHEGGGSGGGSGGGSGLNDF
FEAQKIEWHE*"

misc_feature    351..1238

/label="Halo"

/ApEinfo_revcolor="#75c6a9"

/ApEinfo_fwdcolor="#75c6a9"

/locus_tag="Halo"

/label="Halo"

/ApEinfo_graphicformat="arrow_data {{0 0.5 0 1 2 0 0 -1 0-0.5} {} 0} width 5 offset 0"

misc_feature    1239..1286

/label="Promega TEV optimized site"

/ApEinfo_revcolor="#ff9ccd"

/ApEinfo_fwdcolor="#ff9ccd"

/locus_tag="Promega TEV optimized site"

/label="Promega TEV optimized site"

/ApEinfo_graphicformat="arrow_data {{0 0.5 0 1 2 0 0 -1 0-0.5} {} 0} width 5 offset 0"

misc_feature    1287..2000

/label="GFP"

/ApEinfo_revcolor="#b1ff67"

```
                     /ApEinfo_fwdcolor="#b1ff67"
     misc_feature    2001..2002
                     /label="GS linker"
                     /ApEinfo_revcolor="#c7b0e3"
                     /ApEinfo_fwdcolor="#c7b0e3"
                     /locus_tag="GS linker"
                     /label="GS linker"
                     /ApEinfo_graphicformat="arrow_data {{0 0.5 0 1 2 0 0 -1 0-0.5} {}
0} width 5 offset 0"
     misc_feature    2040..2081
                     /label="AviTag"
                     /ApEinfo_revcolor="#f58a5e"
                     /ApEinfo_fwdcolor="#f58a5e"
     misc_feature    2106..2117
                     /label="GS linker"
                     /ApEinfo_revcolor="#85dae9"
                     /ApEinfo_fwdcolor="#85dae9"
     misc_feature    2121..2162
                     /label="AviTag"
                     /ApEinfo_revcolor="#f58a5e"
                     /ApEinfo_fwdcolor="#f58a5e"
     misc_feature    2321..2361
                     /label="T7 terminator"
                     /ApEinfo_revcolor="#d6b295"
                     /ApEinfo_fwdcolor="#d6b295"
                     /locus_tag="T7 terminator"
                     /label="T7 terminator"
                     /ApEinfo_graphicformat="arrow_data {{0 0.5 0 1 2 0 0 -1 0-0.5} {}
0} width 5 offset 0"
     CDS             complement(3342..4001)
                     /label="AmpR"
                     /ApEinfo_revcolor="#ffef86"
                     /ApEinfo_fwdcolor="#ffef86"
                     /locus_tag="AmpR"
                     /label="AmpR"
```

/ApEinfo_graphicformat="arrow_data {{0 0.5 0 1 2 0 0 -1 0-0.5} {} 0} width 5 offset 0"

/translation="MSTFKVLLCGAVLSRIDAGQEQLGRRIHYSQNDLVEYSPVTEKHLTDGMTVRELCSAAITMSDNTAANLLLTTIGGPKELTAFLHNMGDHVTRLDRWEPELNEAIPNDERDTTMPVAMATTLRKLLTGELLTLASRQQLIDWMEADKVAGPLLRSALPAGWFIADKSGAGERGSRGIIAALGPDGKPSRIVVIYTTGSQATMDERNRQIAEIGASLIKHW"

ORIGIN

1 TCGATGTAAC CCACTCGTGC ACCCAACTGA TCTTCAGCAT CTTTTACTTT CACCAGCGTT

61 TCTGGGTGAG CAAAAACAGG AAGGCAAAAT GCCGCAAAAA AGGGAATAAG GGCGACACGG

121 AAATGTTGAA TACTCATACT CTTCCTTTTT CAATATTATT GAAGCATTTA TCAGGGTTAT

181 TGTCTCATGA GCGGATACAT ATTTGAATGT ATTTAGAAAA ATAAACAAAT AGGGGTTCCG

241 CGCACATTTC CCCGAAAAGT GCTAGTGGTG CTAGCCCCGC GAAATTAATA CGACTCACTA

301 TAGGGTCTAG AAATAATTTT GTTTAACTTT AAGAAGGAGA TATACATATG gcggagattg

361 gcacaggatt tccatttgat ccccactatg ttgaggttct tggtgaacgt atgcactatg

421 ttgacgttgg accacgtgac ggaacccctg ttttattttt gcatggtaat ccgactagta

481 gctacgtttg gcgcaacatc atcccacacg tcgcccctac tcaccgttgt atcgcaccgg

541 acttgatcgg tatggggaag tccgacaagc cagacttagg ctatttcttt gacgaccatg

601 tccgtttcat ggacgcgttt atcgaggccc ttgggttaga agaggttgta cttgtcattc

661 acgactgggg gtccgctctt ggctttcatt gggctaaacg taaccccgag cgcgtcaagg

721 ggatcgcctt tatggagttt atccgcccca tccctacctg ggatgagtgg ccggaatttg

781 cgcgtgaaac gtttcaagcc ttccgcacaa ctgatgttgg ccgcaaatta attatcgatc

841 agaacgtgtt catcgagggg acattgccaa tgggagtggt tcgtccgttg accgaagttg

901 aaatggacca ttatcgcgaa ccgtttctga acccggtcga tcgtgaaccc ttatggcgct

961 ttcctaacga attgccgatt gctggtgaac ctgcgaacat tgtcgcatta gtagaggaat

1021 acatggactg gttacatcaa tcccccgttc ctaaattgtt gttttgggga acgccaggcg

1081 tattaatccc gccagcggag gccgcacgcc ttgcgaagtc acttcctaat tgcaaagcag

1141 tggatattgg cccaggcttg aatcttttgc aagaggataa ccctgatctg atcgggtcgg

1201 aaattgcgcg ttggttatcg acccttgaga tttcgggcGA ACCAACAACT GAGGACTTGT

1261 ATTTCCAATC TGACAACGCA ATCGCTATGT CTAAAGGGGA AGAACTTTTC ACAGGGGTGG

1321 TCCCGATCTT GGTGGAACTT GACGGAGATG TAAATGGGCA CAAATTTTCT GTTTCCGGTG

1381 AAGGAGAGGG TGATGCCACT TATGGTAAAT TAACGTTAAA ATTTATCTGC ACAACCGGAA

1441 AACTGCCTGT GCCCTGGCCT ACACTGGTCA CCACGTTTAG TTACGGGGTT CAGTGCTTCA

1501 GTCGTTACCC AGACCACATG AAACAACATG ATTTCTTTAA ATCCGCGATG CCAGAGGGCT

1561 ATGTCCAGGA GCGTACAATT TTCTTCAAAG ATGACGGAAA CTATAAGACT CGTGCCGAAG

1621 TAAAATTTGA GGGTGATACC CTTGTCAACC GTATCGAGCT GAAGGGCATT GACTTCAAGG

1681 AAGATGGTAA CATCTTAGGT CACAAACTGG AGTACAACTA TAACTCCCAT AATGTGTACA

1741 TCATGGCGGA CAAGCAAAAG AACGGAATCA AGGTAAACTT CAAAATTCGT CATAATATTG

1801 AGGATGGAAG TGTACAGTTA GCCGACCACT ACCAGCAAAA CACCCCCATT GGGGATGGCC

1861 CTGTGCTGCT GCCTGACAAC CACTATCTGT CGACACAGAG CGCATTGTCT AAGGACCCAA

1921 ACGAGAAGCG TGATCACATG GTATTGCTGG AATTCGTAAC GGCTGCTGGC ATTACGCATG

1981 GAATGGATGA GTTATACAAG GGTGGCGGGT CGGGTGGTGG ATCGGGAGGT GGGTCAGGTT

2041 TGAATGACTT TTTCGAGGCA CAGAAAATCG AATGGCATGA AGGCGGCGGG AGTGGAGGGG

2101 GGTCGGGTGG CGGATCTGGG TTGAATGATT TTTTTGAAGC CCAAAAAATT GAGTGGCATG

2161 AATGAGGATC CCGGGAATTC TCGAGTAAGG TTAACCTGCA GGAGGCCTTT AATTAAGGTG

2221 GTGCGGCCGC GCTAGCGGTC CCGGGGGATC GATCCGGCTG CTAACAAAGC CCGAAAGGAA

2281 GCTGAGTTGG CTGCTGCCAC CGCTGAGCAA TAACTAGCAT AACCCCTTGG GGCCTCTAAA

2341 CGGGTCTTGA GGGGTTTTTT GCTGAAAGGA GGAACTATAT CCGGAAGCTT GGCACTGGCC

2401 GACCGGGGTC GAGCACTGAC TCGCTGCGCT CGGTCGTTCG GCTGCGGCGA GCGGTATCAG

2461 CTCACTCAAA GGCGGTAATA CGGTTATCCA CAGAATCAGG GGATAACGCA GGAAAGAACA

2521 TGTGAGCAAA AGGCCAGCAA AAGGCCAGGA ACCGTAAAAA GGCCGCGTTG CTGGCGTTTT

2581 TCCATAGGCT CCGCCCCCCT GACGAGCATC ACAAAAATCG ACGCTCAAGT CAGAGGTGGC

2641 GAAACCCGAC AGGACTATAA AGATACCAGG CGTTTCCCCC TGGAAGCTCC CTCGTGCGCT

2701 CTCCTGTTCC GACCCTGCCG CTTACCGGAT ACCTGTCCGC CTTTCTCCCT TCGGGAAGCG

2761 TGGCGCTTTC TCATAGCTCA CGCTGTAGGT ATCTCAGTTC GGTGTAGGTC GTTCGCTCCA

2821 AGCTGGGCTG TGTGCACGAA CCCCCCGTTC AGCCCGACCG CTGCGCCTTA TCCGGTAACT

2881 ATCGTCTTGA GTCCAACCCG CTAAGACACG ACTTATCGCC ACTGGCAGCA GCCACTGGTA

2941 ACAGGATTAG CAGAGCGAGG TATGTAGGCG GTGCTACAGA GTTCTTGAAG TGGTGGCCTA

3001 ACTACGGCTA CACTAGAAGA ACAGTATTTG GTATCTGCGC TCTGCTGAAG CCAGTTACCT

3061 TCGGAAAAAG AGTTGGTAGC TCTTGATCCG GCAAACAAAC CACCGCTGGT AGCGGTGGTT

3121 TTTTTGTTTG CAAGCAGCAG ATTACGCGCA GAAAAAAAGG ATCTCAAGAA GATCCTTTGA

3181 TCTTTTCTAC GGGGTCTGAC GCTCAGTGGA ACGAAAACTC ACAGATCCGG GATTTTGGTC

3241 ATGAGATTAT CAAAAAGGAT CTTCACCTAG ATCCTTTTAA ATTAAAAATG AAGTTTTAAA

3301 TCAATCTAAA GTATATATGA GTAAACTTGG TCTGACAGTT ACCAATGCTT AATCAGTGAG

3361 GCACCTATCT CAGCGATCTG TCTATTTCGT TCATCCATAG TTGCCTGACT CCCCGTCGTG

3421 TAGATAACTA CGATACGGGA GGGCTTACCA TCTGGCCCCA GTGCTGCAAT GATACCGCGA

3481 GACCCACGCT CACCGGCTCC AGATTTATCA GCAATAAACC AGCCAGCCGG AAGGGCCGAG

3541 CGCAGAAGTG GTCCTGCAAC TTTATCCGCC TCCATCCAGT CTATTAATTG TTGCCGGGAA

3601 GCTAGAGTAA GTAGTTCGCC AGTTAATAGT TTGCGCAACG TTGTTGCCAT TGCTACAGGC

3661 ATCGTGGTGT CACGCTCGTC GTTTGGTATG GCTTCATTCA GCTCCGGTTC CCAACGATCA

3721 AGGCGAGTTA CATGATCCCC CATGTTGTGC AAAAAAGCGG TTAGCTCCTT CGGTCCTCCG

3781 ATCGTTGTCA GAAGTAAGTT GGCCGCAGTG TTATCACTCA TGGTTATGGC AGCACTGCAT

3841 AATTCTCTTA CTGTCATGCC ATCCGTAAGA TGCTTTTCTG TGACTGGTGA GTACTCAACC

3901 AAGTCATTCT GAGAATAGTG TATGCGGCGA CCGAGTTGCT CTTGCCCGGC GTCAATACGG

3961 GATAATACCG CGCCACATAG CAGAACTTTA AAAGTGCTCA TCATTGGAAA ACGTTCTTCG

4021 GGGCGAAAAC TCTCAAGGAT CTTACCGCTG TTGAGATCCA GT

//

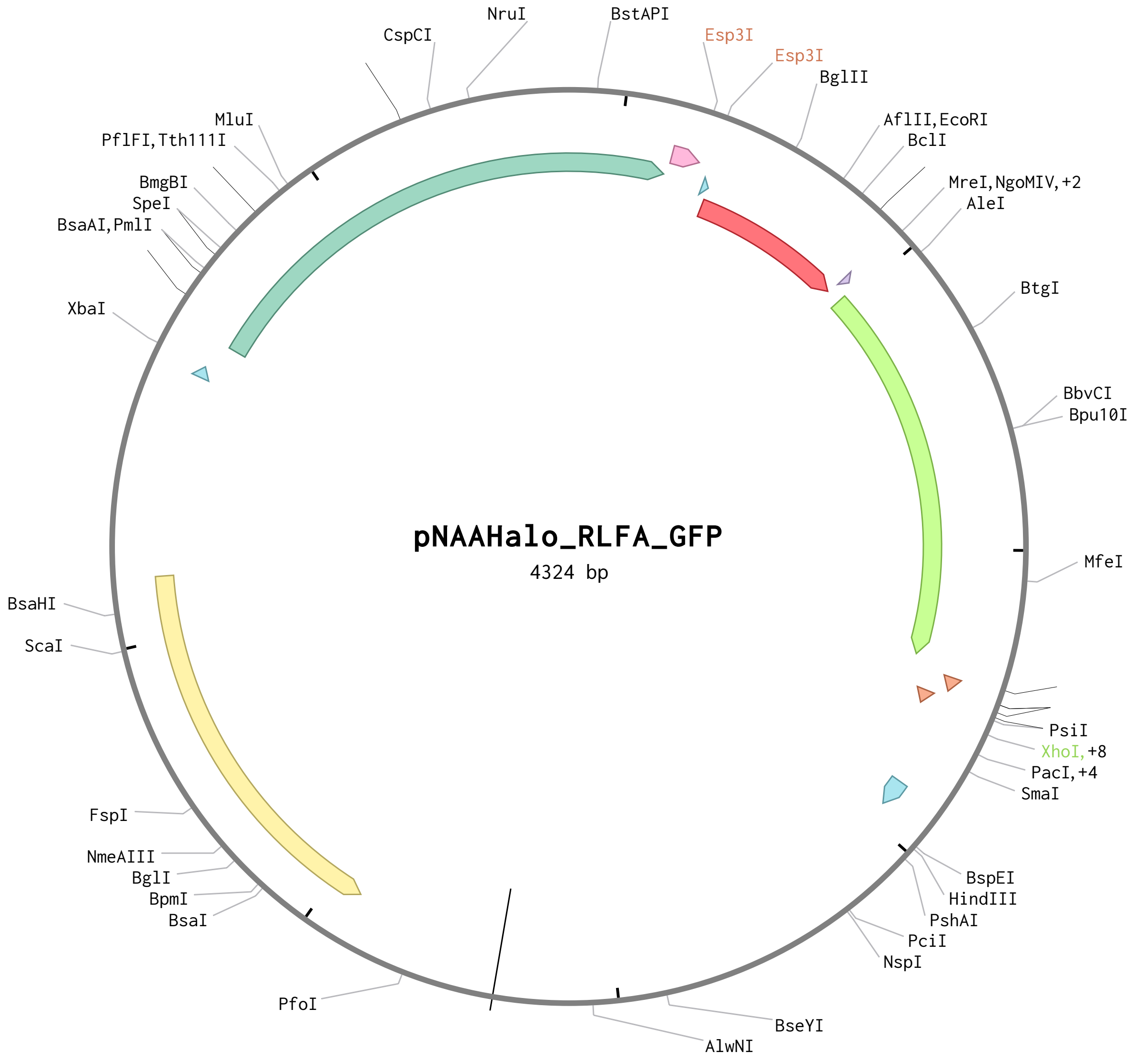

```
LOCUS       pNAAHalo_RLFA_GFP       4324 bp ds-DNA     circular     06-MAR-2025
DEFINITION  .
FEATURES             Location/Qualifiers
     primer_bind     128..147
                     /label="T7"
                     /ApEinfo_revcolor="#85dae9"
                     /ApEinfo_fwdcolor="#85dae9"
                     /locus_tag="T7"
                     /label="T7"
                     /ApEinfo_graphicformat="arrow_data {{0 0.5 0 1 2 0 0 -1 0-0.5} {}
0} width 5 offset 0"
     misc_feature    196..1083
                     /label="Halo"
                     /ApEinfo_revcolor="#75c6a9"
                     /ApEinfo_fwdcolor="#75c6a9"
                     /locus_tag="Halo"
                     /label="Halo"
                     /ApEinfo_graphicformat="arrow_data {{0 0.5 0 1 2 0 0 -1 0-0.5} {}
0} width 5 offset 0"
     misc_feature    1090..1137
                     /label="Promega TEV optimized site"
                     /ApEinfo_revcolor="#ff9ccd"
                     /ApEinfo_fwdcolor="#ff9ccd"
     misc_feature    1156..1167
                     /label="GS linker"
                     /ApEinfo_revcolor="#85dae9"
                     /ApEinfo_fwdcolor="#85dae9"
                     /locus_tag="GS linker"
                     /label="GS linker"
                     /ApEinfo_graphicformat="arrow_data {{0 0.5 0 1 2 0 0 -1 0-0.5} {}
0} width 5 offset 0"
     misc_feature    1168..1457
                     /label="scSUMO"
                     /ApEinfo_revcolor="#ff3843"
                     /ApEinfo_fwdcolor="#ff3843"
                     /locus_tag="scSUMO"
```

```
                     /label="scSUMO"
                     /ApEinfo_graphicformat="arrow_data {{0 0.5 0 1 2 0 0 -1 0-0.5} {}
0} width 5 offset 0"
     misc_feature    1462..1473
                     /label="RLFA"
                     /ApEinfo_revcolor="#c7b0e3"
                     /ApEinfo_fwdcolor="#c7b0e3"
     misc_feature    1486..2199
                     /label="GFP"
                     /ApEinfo_revcolor="#b1ff67"
                     /ApEinfo_fwdcolor="#b1ff67"
     misc_feature    2221..2244
                     /label="FLAG"
                     /ApEinfo_revcolor="#f58a5e"
                     /ApEinfo_fwdcolor="#f58a5e"
     misc_feature    2257..2280
                     /label="FLAG"
                     /ApEinfo_revcolor="#f58a5e"
                     /ApEinfo_fwdcolor="#f58a5e"
     attenuator      2419..2465
                     /label="T7 terminator"
                     /ApEinfo_revcolor="#85dae9"
                     /ApEinfo_fwdcolor="#85dae9"
                     /locus_tag="T7 terminator"
                     /label="T7 terminator"
                     /ApEinfo_graphicformat="arrow_data {{0 0.5 0 1 2 0 0 -1 0-0.5} {}
0} width 5 offset 0"
     CDS             complement(3446..4105)
                     /label="AmpR"
                     /ApEinfo_revcolor="#ffef86"
                     /ApEinfo_fwdcolor="#ffef86"
ORIGIN
        1 TTGAAGCATT TATCAGGGTT ATTGTCTCAT GAGCGGATAC ATATTTGAAT GTATTTAGAA
       61 AAATAAACAA ATAGGGGTTC CGCGCACATT TCCCCGAAAA GTGCTAGTGG TGCtagcccc
      121 gcgaaattaa tacgACTCAC TATAGGGTCT AGAAGAAATA ATTTTGTTTA ACTTTAAGAA
      181 GGAGATATAC ATATGgcgga gattggcaca ggatttccat ttgatcccca ctatgttgag
      241 gttcttggtg aacgtatgca ctatgttgac gttggaccac gtgacggaac ccctgtttta
```

301 tttttgcatg gtaatccgac tagtagctac gtttggcgca acatcatccc acacgtcgcc

361 cctactcacc gttgtatcgc accggacttg atcggtatgg ggaagtccga caagccagac

421 ttaggctatt tctttgacga ccatgtccgt ttcatggacg cgtttatcga ggcccttggg

481 ttagaagagg ttgtacttgt cattcacgac tgggggtccg ctcttggctt tcattgggct

541 aaacgtaacc ccgagcgcgt caaggggatc gcctttatgg agtttatccg ccccatccct

601 acctgggatg agtggccgga atttgcgcgt gaaacgtttc aagccttccg cacaactgat

661 gttggccgca aattaattat cgatcagaac gtgttcatcg aggggacatt gccaatggga

721 gtggttcgtc cgttgaccga agttgaaatg gaccattatc gcgaaccgtt tctgaacccg

781 gtcgatcgtg aacccttatg gcgctttcct aacgaattgc cgattgctgg tgaacctgcg

841 aacattgtcg cattagtaga ggaatacatg gactggttac atcaatcccc cgttcctaaa

901 ttgttgtttt ggggaacgcc aggcgtatta atcccgccag cggaggccgc acgccttgcg

961 aagtcacttc ctaattgcaa agcagtggat attggcccag gcttgaatct tttgcaagag

1021 gataaccctg atctgatcgg gtcggaaatt gcgcgttggt tatcgaccct tgagatttcg

1081 ggcACCAGCG AACCAACAAC TGAGGACTTG TATTTCCAAT CTGACAACGC AATCGCTaGA

1141 GACGgctaCG TCTCgGGTGG CGGATCTATG TCGGACTCAG AAGTCAATCA AGAAGCTAAG

1201 CCAGAGGTCA AGCCAGAAGT CAAGCCTGAG ACTCACATCA ATTTAAAGGT GTCCGATGGA

1261 TCTTCAGAGA TCTTCTTCAA GATCAAAAAG ACCACTCCTT TAAGAAGGCT GATGGAAGCG

1321 TTCGCTAAAA GACAGGGTAA GGAAATGGAC TCCTTAAGAT TCTTGTACGA CGGTATTAGA

1381 ATTCAAGCTG ATCAGACCCC TGAAGATTTG GACATGGAGG ATAACGATAT TATTGAGGCT

1441 CACAGAGAAC AGATTGGTGG TCGCTTATTC GCCGGCGGGG GGTCCATGAG TAAGGGTGAA

1501 GAATTATTCA CGGGGGTGGT GCCAATCTTG GTCGAACTGG ACGGCGATGT AAACGGACAC

1561 AAGTTTTCCG TGTCAGGTGA AGGTGAAGGC GATGCGACTT ATGGCAAACT TACGCTTAAA

1621 TTCATTTGCA CAACCGGCAA ACTGCCCGTT CCGTGGCCGA CTTTGGTTAC TACATTTAGT

1681 TACGGAGTTC AATGCTTCTC ACGCTACCCC GACCATATGA AACAGCACGA TTTTTTTAAA

1741 TCGGCGATGC CCGAAGGCTA TGTGCAAGAG CGTACAATTT TTTTTAAAGA CGATGGAAAT

1801 TACAAAACCC GCGCTGAGGT CAAGTTCGAG GGCGATACTC TTGTTAATCG CATCGAGCTG

1861 AAAGGTATTG ATTTTAAGGA AGATGGCAAT ATTTTGGGAC ACAAGTTAGA ATATAACTAT

1921 AACTCACATA ATGTCTATAT TATGGCGGAC AAACAGAAGA ACGGCATTAA GGTGAACTTT

1981 AAAATTCGTC ACAATATCGA GGACGGCTCC GTCCAATTAG CCGACCATTA CCAACAAAAT

2041 ACGCCAATTG GTGACGGGCC GGTGTTGTTA CCAGACAACC ATTACCTTTC TACACAAAGC

2101 GCATTATCAA AAGATCCTAA TGAAAAGCGC GATCACATGG TTTTGCTTGA ATTTGTCACA

2161 GCCGCAGGCA TCACGCACGG CATGGATGAG TTATATAAGG GTGGCGGCTC GGGCGGTTCA

2221 GATTACAAAG ACGATGACGA TAAGGGTGGC GGATCAGATT ATAAGGACGA TGACGACAAA

2281 TGACTCGAGT AAGGTTAACC TGCAGGAGGC CTTTAATTAA GGTGGTGCGG CCGCGCTAGC

2341 GGTCCCGGGG GATCGATCCG GCTGCTAACA AAGCCCGAAA GGAAGCTGAG TTGGCTGCTG

2401 CCACCGCTGA GCAATAACTA GCATAACCCC TTGGGGCCTC TAAACGGGTC TTGAGGGGTT

2461 TTTTGCTGAA AGGAGGAACT ATATCCGGAA GCTTGGCACT GGCCGACCGG GGTCGAGCAC

2521 TGACTCGCTG CGCTCGGTCG TTCGGCTGCG GCGAGCGGTA TCAGCTCACT CAAAGGCGGT

2581 AATACGGTTA TCCACAGAAT CAGGGGATAA CGCAGGAAAG AACATGTGAG CAAAAGGCCA

2641 GCAAAAGGCC AGGAACCGTA AAAAGGCCGC GTTGCTGGCG TTTTTCCATA GGCTCCGCCC

2701 CCCTGACGAG CATCACAAAA ATCGACGCTC AAGTCAGAGG TGGCGAAACC CGACAGGACT

2761 ATAAAGATAC CAGGCGTTTC CCCCTGGAAG CTCCCTCGTG CGCTCTCCTG TTCCGACCCT

2821 GCCGCTTACC GGATACCTGT CCGCCTTTCT CCCTTCGGGA AGCGTGGCGC TTTCTCATAG

2881 CTCACGCTGT AGGTATCTCA GTTCGGTGTA GGTCGTTCGC TCCAAGCTGG GCTGTGTGCA

2941 CGAACCCCCC GTTCAGCCCG ACCGCTGCGC CTTATCCGGT AACTATCGTC TTGAGTCCAA

3001 CCCGCTAAGA CACGACTTAT CGCCACTGGC AGCAGCCACT GGTAACAGGA TTAGCAGAGC

3061 GAGGTATGTA GGCGGTGCTA CAGAGTTCTT GAAGTGGTGG CCTAACTACG GCTACACTAG

3121 AAGAACAGTA TTTGGTATCT GCGCTCTGCT GAAGCCAGTT ACCTTCGGAA AAAGAGTTGG

3181 TAGCTCTTGA TCCGGCAAAC AAACCACCGC TGGTAGCGGT GGTTTTTTTG TTTGCAAGCA

3241 GCAGATTACG CGCAGAAAAA AAGGATCTCA AGAAGATCCT TTGATCTTTT CTACGGGGTC

3301 TGACGCTCAG TGGAACGAAA ACTCACAGAT CCGGGATTTT GGTCATGAGA TTATCAAAAA

3361 GGATCTTCAC CTAGATCCTT TTAAATTAAA AATGAAGTTT TAAATCAATC TAAAGTATAT

3421 ATGAGTAAAC TTGGTCTGAC AGTTACCAAT GCTTAATCAG TGAGGCACCT ATCTCAGCGA

3481 TCTGTCTATT TCGTTCATCC ATAGTTGCCT GACTCCCCGT CGTGTAGATA ACTACGATAC

3541 GGGAGGGCTT ACCATCTGGC CCCAGTGCTG CAATGATACC GCGAGACCCA CGCTCACCGG

3601 CTCCAGATTT ATCAGCAATA AACCAGCCAG CCGGAAGGGC CGAGCGCAGA AGTGGTCCTG

3661 CAACTTTATC CGCCTCCATC CAGTCTATTA ATTGTTGCCG GGAAGCTAGA GTAAGTAGTT

3721 CGCCAGTTAA TAGTTTGCGC AACGTTGTTG CCATTGCTAC AGGCATCGTG GTGTCACGCT

3781 CGTCGTTTGG TATGGCTTCA TTCAGCTCCG GTTCCCAACG ATCAAGGCGA GTTACATGAT

3841 CCCCCATGTT GTGCAAAAAA GCGGTTAGCT CCTTCGGTCC TCCGATCGTT GTCAGAAGTA

3901 AGTTGGCCGC AGTGTTATCA CTCATGGTTA TGGCAGCACT GCATAATTCT CTTACTGTCA

3961 TGCCATCCGT AAGATGCTTT TCTGTGACTG GTGAGTACTC AACCAAGTCA TTCTGAGAAT

4021 AGTGTATGCG GCGACCGAGT TGCTCTTGCC CGGCGTCAAT ACGGGATAAT ACCGCGCCAC

4081 ATAGCAGAAC TTTAAAAGTG CTCATCATTG GAAAACGTTC TTCGGGGCGA AAACTCTCAA

4141 GGATCTTACC GCTGTTGAGA TCCAGTTCGA TGTAACCCAC TCGTGCACCC AACTGATCTT

4201 CAGCATCTTT TACTTTCACC AGCGTTTCTG GGTGAGCAAA AACAGGAAGG CAAAATGCCG

4261 CAAAAAAGGG AATAAGGGCG ACACGGAAAT GTTGAATACT CATACTCTTC CTTTTTCAAT

4321 ATTA

//
